# Pan-Cancer and Single-Cell modelling of genomic alterations through gene expression

**DOI:** 10.1101/492561

**Authors:** Daniele Mercatelli, Forest Ray, Federico M. Giorgi

## Abstract

Cancer is a disease often characterized by the presence of multiple genomic alterations, which trigger altered transcriptional patterns and gene expression, which in turn sustain the processes of tumorigenesis, tumor progression and tumor maintenance. The links between genomic alterations and gene expression profiles can be utilized as the basis to build specific molecular tumorigenic relationships. In this study we perform pan-cancer predictions of the presence of single somatic mutations and copy number variations using machine learning approaches on gene expression profiles. We show that gene expression can be used to predict genomic alterations in every tumor type, where some alterations are more predictable than others. We propose gene aggregation as a tool to improve the accuracy of alteration prediction models from gene expression profiles. Ultimately, we show how this principle can be beneficial in intrinsically noisy datasets, such as those based on single cell sequencing.

**Author Summary:** In this article we show that transcript abundance can be used to predict the presence or absence of the majority of genomic alterations present in human cancer. We also show how these predictions can be improved by aggregating genes into small networks to counteract the effects of transcript measurement noise.

## Introduction

Cancer is a molecular disease occurring when a cell or group of cells acquire uncontrolled proliferative behavior, conferred by a multitude of deregulations in specific pathways [1]. As is implied by such a broad definition, cancer is a highly heterogeneous disease, showing remarkably different molecular, histological, genetic and clinical properties, even when comparing tumors originating from the same tissue [2], Many cancers are characterized by the presence of single nucleotide or short indel mutations and/or copy number alterations, which appear somatically at the early stages of oncogenesis and can drive tumor progression [3]. Cancers can be broadly divided in two classes: the M class, where point mutations are prevalent, and the C class, where copy number variations (CNVs) are more numerous and are often associated with TP53 mutations. Tumor class influences anatomic location. Most ovarian cancers, for example, belong to the C class, while most colorectal cancers belong to the M class, although many exceptions do exist [4].

The Cancer Genome Atlas (TCGA) project [5] has recently undergone a major effort to collect vast amounts of information on thousands of distinct tumor samples. The TCGA data collection, commonly referred to as the “Pan-cancer” dataset, provided the scientific community with an avalanche of data on DNA alterations, gene expression, methylation status and protein abundances among others, with the critical mass necessary to identify rarer driver tumorigenesis effects in many types of cancers [6–8]. By combining all 33 TCGA datasets, Bailey and colleagues [9] recently outlined a pan-cancer map of which mutations can be drivers for the progression of cancer.

The availability of thousands of samples measuring many different variables in cancer has allowed scientists to generate statistical models of relationships between different molecular species. A pan-cancer correlation network between coding genes and long noncoding RNAs, for example, sheds light on the function of non-coding parts of the transcriptome [10]. More recently, mutations on transcription factors (TFs) have been linked to altered gene expressions and phosphoprotein levels in 12 TCGA tumor type datasets [11]. Network approaches have been applied to identify clusters of coexpressed genes, shared by multiple cancer types [12], Several studies have sought to characterize the relationships between genomic status and expression levels in cancer, trying to identify commonalities across different cancer types [13,14], In particular, Alvarez and colleagues [15] have postulated that the effect of genomic alterations in cancer can be more readily assessed by aggregating gene expression profiles into transcriptional networks, rather than by profiles taken separately.

While the association between genomic events and gene expression is proven in several scenarios, it remains to be seen if it can be assessed in scenarios where fully quantitative readouts are unavailable, such as low coverage samples. One of these scenarios is Single Cell Sequencing [16], often carried out in experiments where thousands of mutations are generated via a system of pooled CRISPR-Cas9 knockouts [17].

To our knowledge there is no study trying to identify relationships between all genomic alteration events (somatic mutations/indels and CNVs) and global gene expression across cancers. In this study, we use 24 TCGA tumor datasets to investigate whether gene expression can be used to predict the presence of specific genomic alterations in several cancer tissue contexts. To this end, we leverage the current availability of a vast family of machine learning algorithms [18]. We investigate whether some gene alterations can be better modelled than others, and whether using grouped gene expression profiles as aggregated variables can effectively identify specific genomic alterations. Finally, we test whether predicting mutations and CNVs can be carried out in an intrinsically noisy single cell RNA-Seq (scRNA-Seq) transcriptomics datasets.

## Results

### Collection of Pan-Cancer Dataset

We downloaded the most recent version of the TCGA datasets available on Firehose (v2016_01_28), encompassing mutational, CNV and gene expression data. Using T-distributed stochastic neighbor embedding (TSNE [19]) clustering on gene expression data (9642 samples), we observed how different tumor types cluster separately from each other (Figure 1A). However, two tumour types segregate into two subgroups: breast cancer, which subdivides into a major luminal cluster and a smaller (in terms of samples collected) basal cluster [20]; and esophageal carcinoma, which roughly subdivides into adenocarcinomas and squamous cell carcinomas [21].

**Figure 1.**
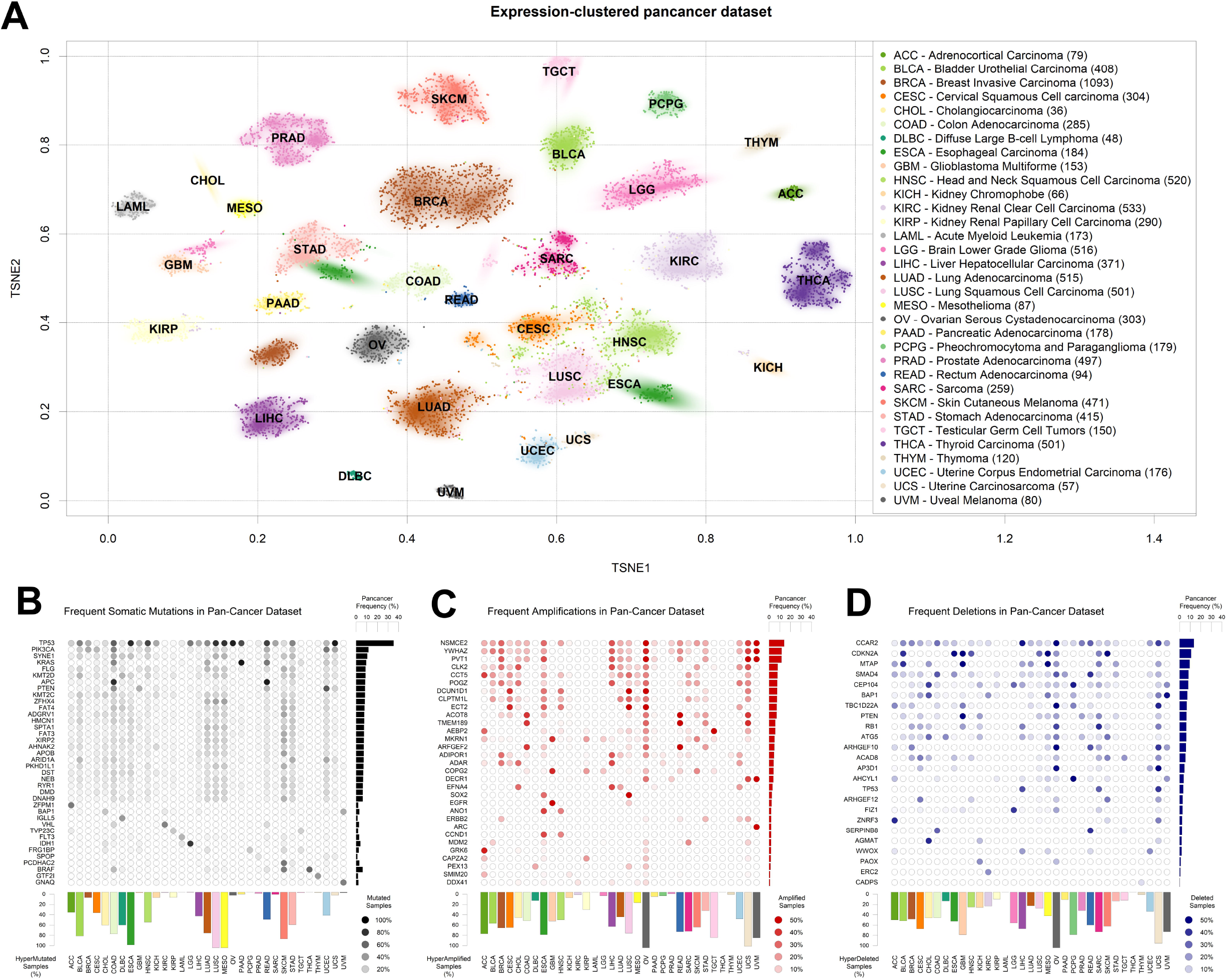
The TCGA dataset used. A: TSNE clustering of TCGA samples based on gene expression profiles. The 2D median of each tumor type is indicated using the TCGA tumor code. Subset size is indicated in brackets next to tumor type names to the right. B: table of most somatically mutated genes across TCGA tumor samples, in terms of number of samples where the gene is somatically mutated with altered protein product sequence. C: table of most amplified genes across TCGA tumor samples. D: table of most deleted genes across TCGA tumor samples. The fraction of total TCGA samples carrying a gene-targeting event is indicated to the right of panels B-D, and the fraction of samples where more than 0.5% of the genes is affected by the panel event type is indicated to the bottom of panels B-D.

We then aggregated the single nucleotide and short indel somatic mutation data from the same samples for which we had collected gene expression. As is widely known, TP53 is the most mutated gene in human cancer (Figure 1B), followed by PIK3CA, SYNE1 and KRAS. As shown before [4] some tumor types are characterized by a high presence of somatic mutations. In particular, lung squamous carcinoma, mesothelioma and esophageal cancer carry at least one of these events in almost 100% of the samples in the TCGA dataset. In the figure, we filtered out commonly known non-driver mutations [22], such as those happening in long genes like TTN and OBSCN, but we kept them in all following analyses for the sake of completion. A representation of all mutated genes, including blacklisted ones, is available in Figure SI. Some tumors are characterized by the prevalence of a mutation in a specific gene, such as the G-protein coding BRAF in thyroid carcinoma [23] or IDH1, translating into isocitrate dehydrogenase, in low grade glioma [24].

Finally, we obtained readouts of CNV status for all TCGA samples. CNVs can have different extensions in terms of nucleotides affected and can sometimes encompass entire chromosomes [25] and the thousands of genes therein. In order to limit the number of variables to a more meaningful subset, we assigned a CNV score to every gene, according the copy number score of the genomic region most overlapping with the UCSC annotated gene boundaries (genome version hg19). We then tested models for all genes affected by a CNV in at least 10 samples (extending what previously done in [26]). In order to make CNV variables comparable to the mutational ones, we defined a cut-off for presence or absence by using the log_2_(CNV) threshold of 0.5, which roughly corresponds to at least one copy gain for amplifications, and at least one copy loss for deletions (see Materials and Methods). We then reported their abundance in the pan-cancer dataset, distinguishing between amplifications (Figure 1C) and deletions (Figure 1D). As previously shown [4], virtually all ovarian cancer samples are characterized by at least one CNV event. Among the most amplified genes, we find the oncogenes SOX2 [27], EGFR [28] and MDM2 [29], and also a non-coding gene, PVT1, the most amplified gene in breast cancer, with proven but as-of-yet uncharacterized proto-oncogenic effects [30,31]. Amongst the most deleted genes (Fig. 1D) we observe well known tumor-suppressor genes, such as CDKN2A [32,33] and PTEN [34,35].

### Modelling Cancer Alterations with gene expression

After collecting all the expression and genomic alteration data from TCGA, we set out to generate models able to predict the presence or absence of each event by virtue of gene expression data in the contexts of all collected tumor types.

We tested several modelling algorithms for classification using the aggregator platform for machine learning caret [18] in the bladder cancer mutational dataset [36]. We observed that all models provide better-than-random predictions for the majority of mutational events, in terms of area under the ROC curve (AUROC)(Figure 2) [37], We chose the top-scoring algorithm in this test, the Gradient Boost Modelling algorithm (gbm), a robust tree-based boosting model [38], due to its robustness and speed of implementation.

**Figure 2.**
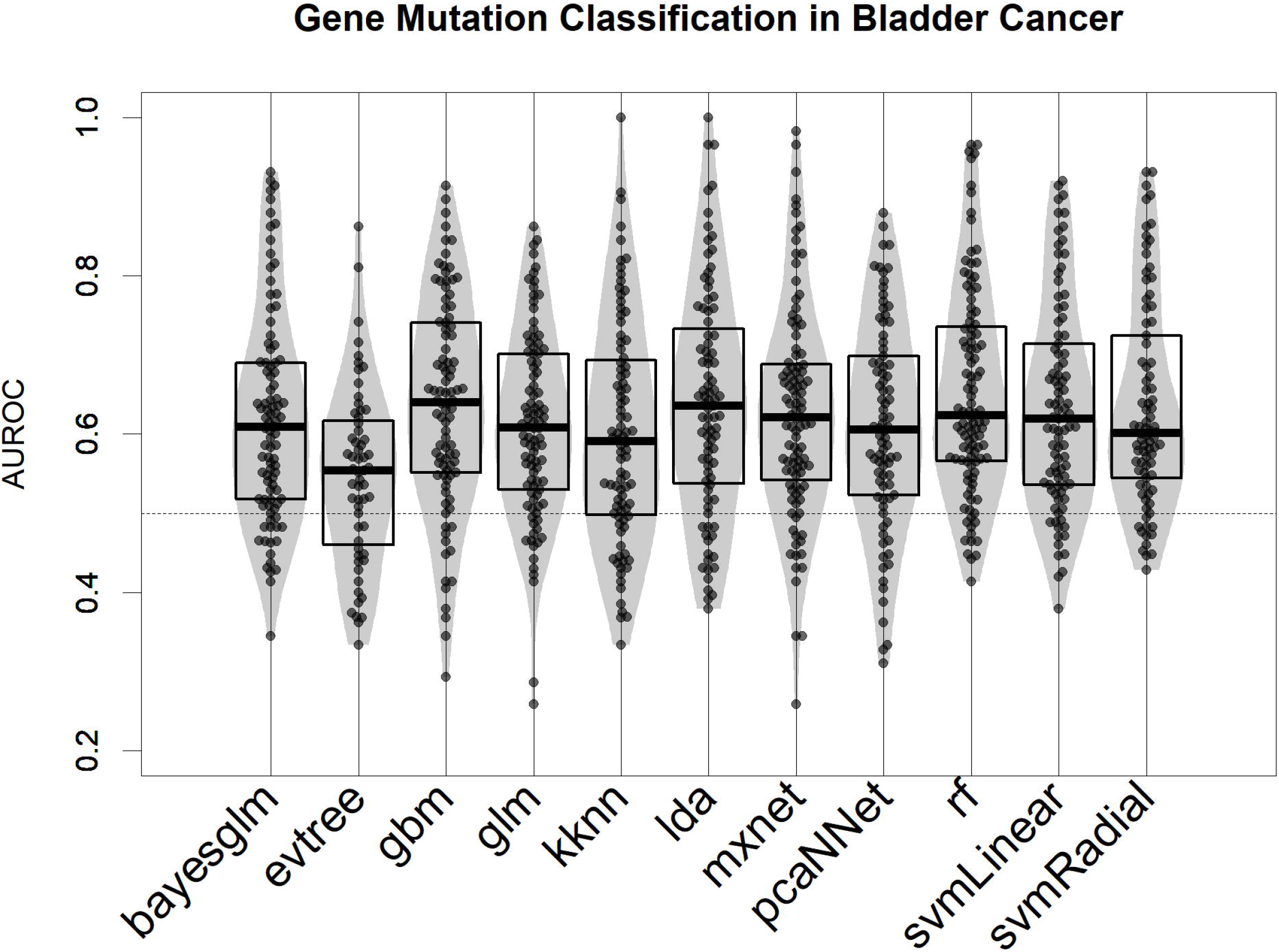
Performance of 11 machine learning algorithms in binary classification of mutated/nonmutated samples using gene expression predictor variables in the Bladder Cancer dataset. Each point corresponds to a specific mutation/model. Performance is indicated as AUROC: Area Under the Receiver Operating Characteristic curve.

We calculated gbm models for all tumour types of at least 100 samples with co-measured expression and CNV or mutations, which included 24 of the 33 TCGA tumor types. The models were predictive of genomic events observed in no less than 5% and no more than 95% of the patients in the dataset, and at least in 10 samples. Our results show that in all tumour types, a machine learning algorithm based on gene expression is consistently better than a random predictor (AUROC line at 0.5) at correctly classifying tumour samples for the presence or absence of specific genomic alteration events (Figure 3 and Supplementary Table S1). In particular, TP53 mutations are well modelled in many of these tumor types, being the most well predicted mutational event in both acute myeloid leukemia and low grade glioma. We could also model the presence of a copy loss of TP53 in sarcoma, which can be predicted with an accuracy of 70%. Ovarian and pancreatic cancer datasets presented exceptional cases, in that each contained such high TP53 mutation rates (next to 95% detected) [39,40] that our algorithms could not distinguish sufficient differences within each dataset to train a model. Also KRAS-targeting events are well modelled, specifically in colon, lung and stomach cancer, and cervical squamous carcinoma [41]. We noted a tendency where models for more frequent CNV events yielded a greater predictive power (Figure S2), a tendency not observed for somatic mutation models. We then tested if known tumor-related genes, such as those curated by the Cancer Gene Census [42] are better modelled than the rest of the genome. There is no difference in mutation and amplification results, but for deletion events, oncogenes yield weaker models (Wilcoxon Test, p=0.0037, figure S3) and tumor suppressor genes yield generally stronger models (p=0.00050). This is in agreement with the central paradigm of cancer, where a tumor suppressor gene deletion can be one of the driving events of tumorigenesis and tumor progression [43]. On the other hand, deletion of tumor-promoting oncogenes is generally unfavourable for tumor progression, and so, generally speaking it should be present only as a passenger event, unlikely to determine global gene expression and tumor fate.

**Figure 3.**
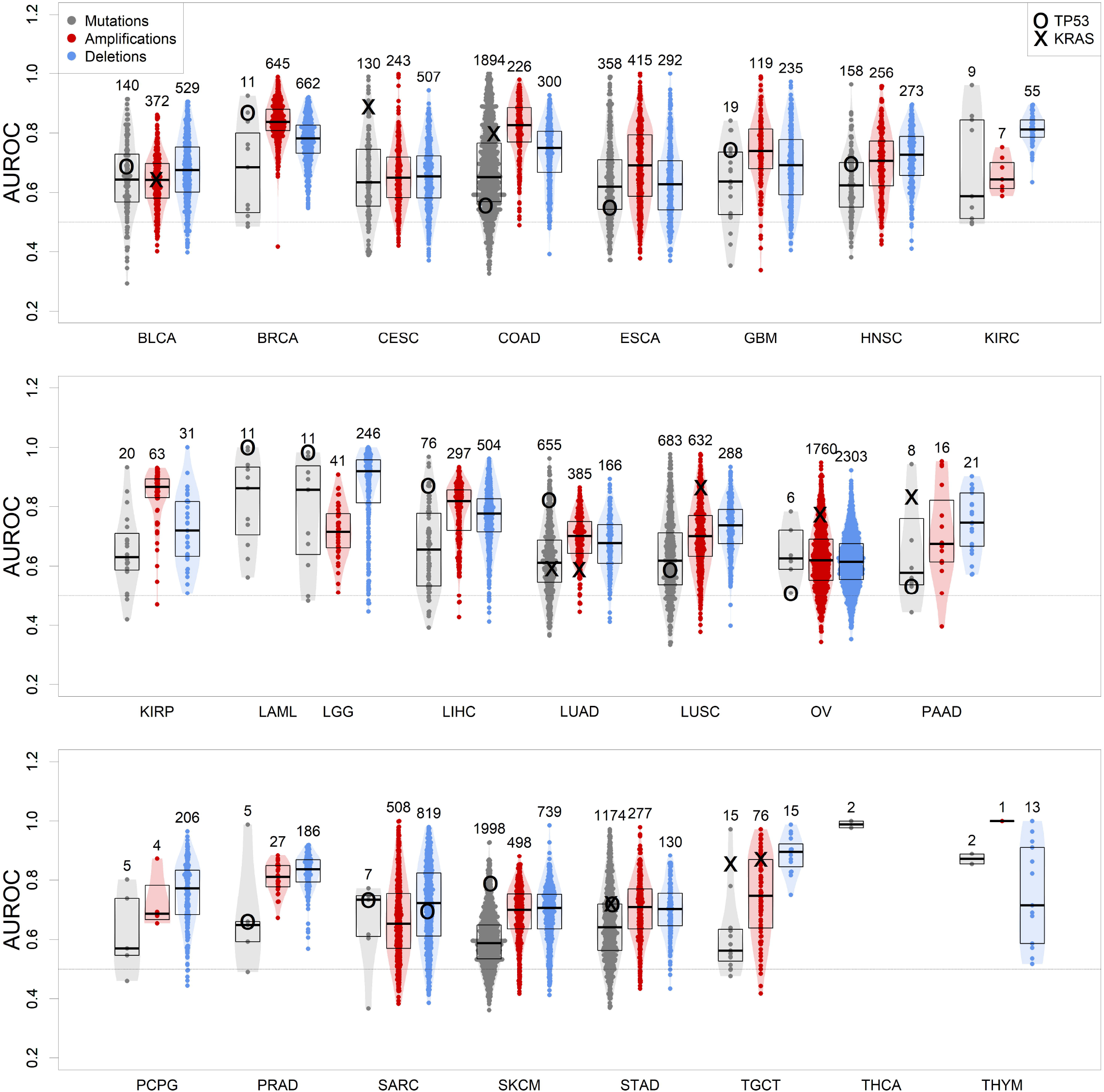
Performance of gbm models for each genomic alteration event in TCGA, predicted as a function of each tumor gene expression. Boxplots indicate distribution median, upper and lower quartile. Alterations targeting TP53 and KRAS are indicated.

### Modelling specific alterations with noise addition

In order to understand whether cancer-related genomic alterations can be modelled by gene expression in scenarios with lower signal-to-noise ratio, we artificially perturbed the TCGA gene expression dataset via the addition of Gaussian noise, and then proceeded to build models to predict the presence of TP53 mutations in breast cancer, the largest dataset in TCGA by number of samples.

As expected, the addition of uniform random gaussian noise to the gene expression matrix has a detrimental effect on the amount of information left for modelling the presence of TP53 somatic mutations (Figure 4A).

**Figure 4.**
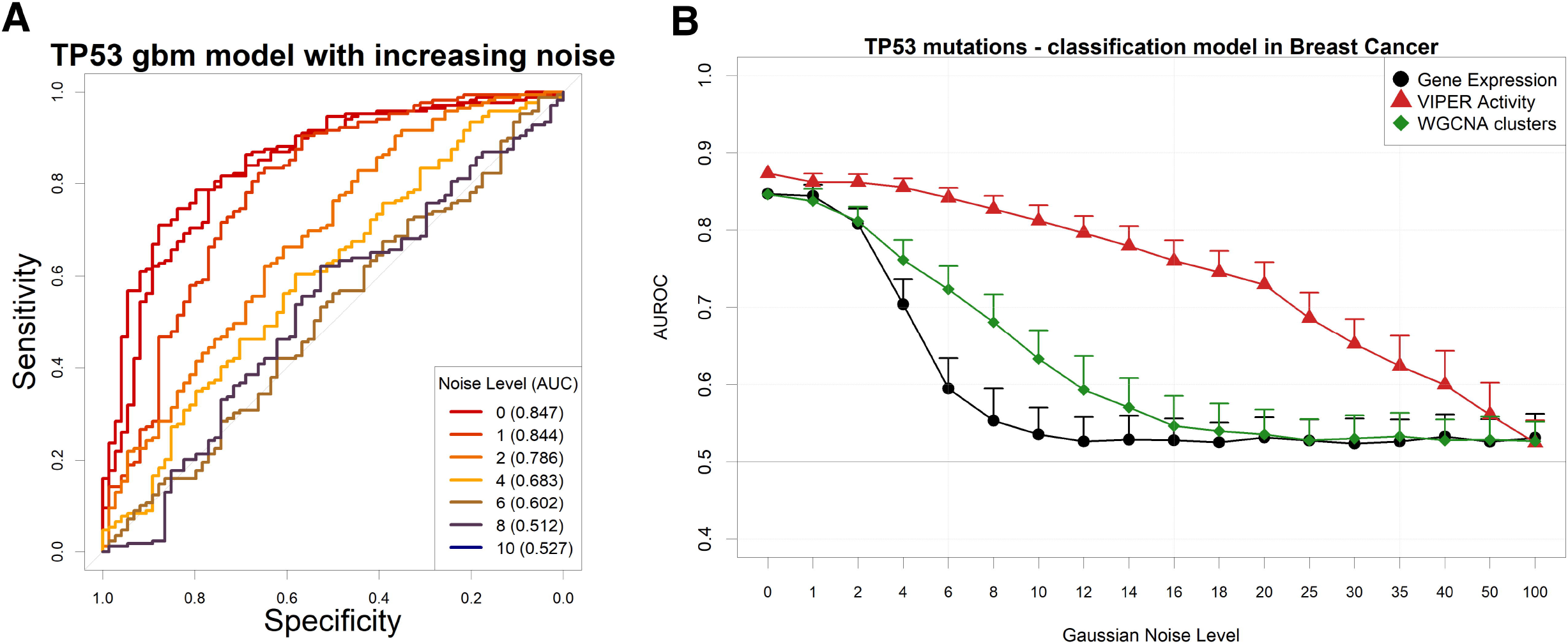
Performance of a TP53 somatic mutation gbm model upon gaussian noise addiction. A: ROC curves (and AUC) upon addition of increasing levels (in terms of SD of a gaussian distribution with mean=0) of gaussian noise. B: AUROCs of the model with increasing noise, calculated using gene expression (black line) or aggregated gene expression using the WGCNA (green line) or VIPER (red line) algorithms. Pseudocounts of 0.1 are added in order to show zero counts as −1 in log_10_ scale.

We then decided to test several permutations of noise addition on the same breast cancer expression data, by each time aggregating genes into networks defined a priori in the same context, using a Tukey Biweight Robust Average method [44] on Weighted Gene Correlation Network Analysis (WGCNA) clusters [45] and the VIPER algorithm [15] on ARACNe-AP networks [46]. It is important to note that WGCNA clusters are completely non-overlapping and yield generally a lower number of aggregated variables than VIPER clusters, which are groups of genes possibly shared by other transcription factor clusters and that collectively yield the global expression of a transcription factor target set (dubbed as a proxy for “TF activity” in the original VIPER manuscript [15]).

Our results show that gene expression, VIPER activity and WGCNA clusters yield very similar models for predicting TP53 mutations in breast cancer (figure S4). The amount of information contained in the input variables is therefore comparable. Adding noise to the input expression matrix, however, and then aggregating the resulting noise-burdened genes into VIPER or WGCNA clusters (see Materials and Methods), provides robustness to the models (Figure 4B). Similar results with higher variances (possibly due to the smaller size of the datasets) can be observed for EGFR amplifications in glioblastoma (Figure S5) and lung squamous carcinoma (Figure S6), for PVT1 amplifications in ovarian cancer (Figure S7) and for PTEN deletions in sarcoma (Figure S8). In all these examples, however, the performance of the simple WGCNA/Tukey aggregation is closer (if not worse) to that of simple gene expression.

An alternative way to reduce the information content from an NGS gene expression dataset is to reduce the number of read counts from each sample. This operation reflects either a low coverage bulk RNA-Seq experiment or an experiment arising from Single-Cell sequencing [47], In particular, single-cell RNA-Seq (scRNA-Seq) is characterized by the dropout phenomenon [48] wherein genes expressed in the cells are sometimes not detected at all. In order to simulate such scenarios, we down-sampled each RNA-Seq gene count profile from the largest TCGA dataset (Breast Cancer) to a target aligned read number using a beta function, which allows for reduction coupled with random complete gene dropouts (Figure 5A). We then modelled again the presence of TP53 mutations using gene expression (Figure 5B). We found out that models based on standard unaggregated gene expression experience an accuracy drop at around 30M reads, while aggregating genes using VIPER (but not with WGCNA) allows for better-than-random accuracies even at 3M reads, confirming the benefits of gene aggregation in low coverage RNA-Seq, as previously found e.g. for sample clustering [49].

**Figure 5.**
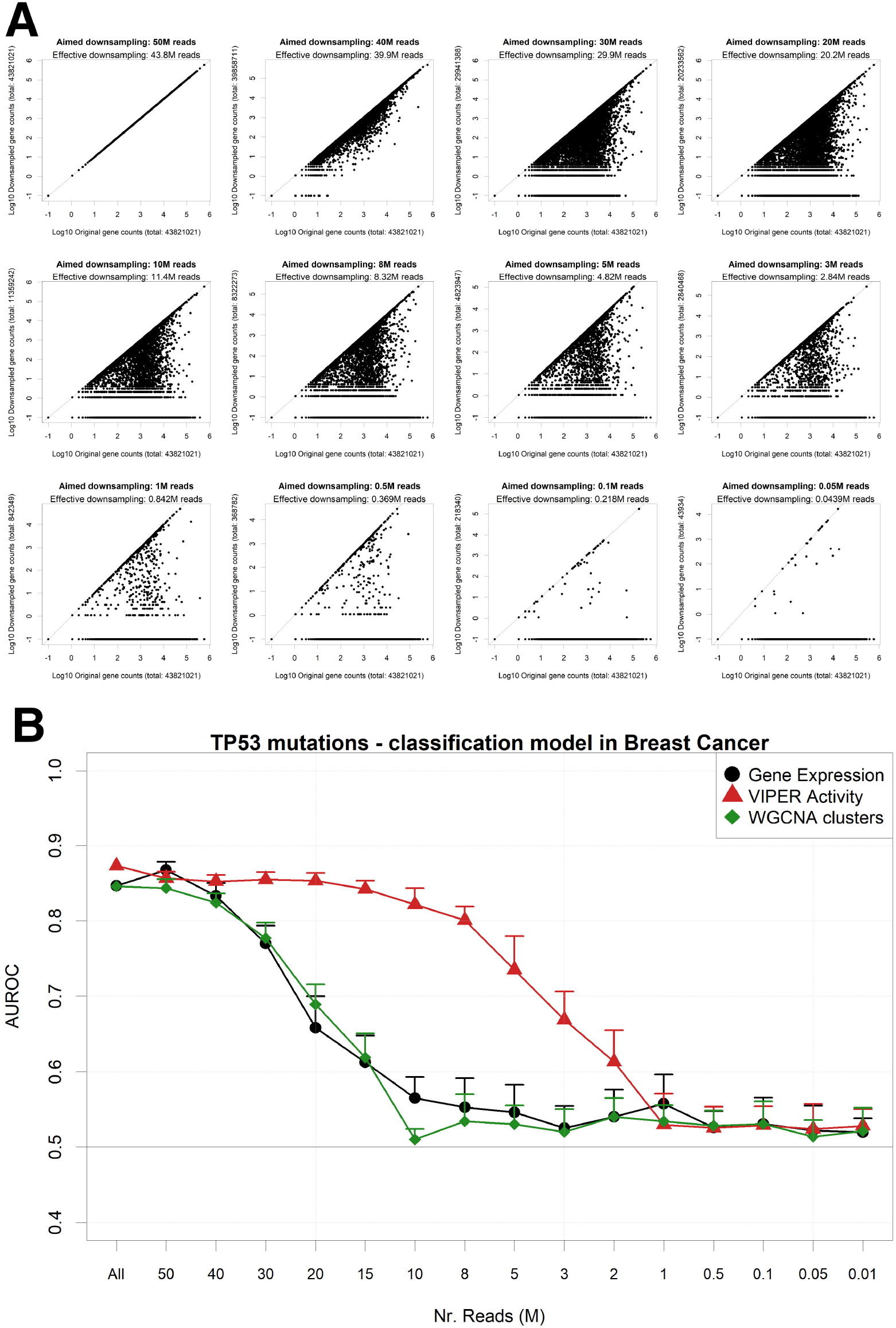
Performance of a TP53 mutation gbm model upon downsampling of the TCGA breast cancer RNA-Seq dataset. A: for a single TCGA sample (TCGA-A1-A0SB-01) with 43.8 gene mapping reads, the downsampling algorithm is applied for multiple target read quantities. X-axis shows the count for each gene in the original sample, and Y-axis in the downsampled output. B: AUROCs of the model with decreasing read numbers, calculated using gene expression (black line) or aggregated gene expression using the WGCNA (green line) or VIPER (red line) algorithms.

### Mutation prediction in single-cell data

Based on the results from the pan-cancer analysis, where we predicted sample mutations based on pooled RNA-Seq gene expression patterns, we decided to extend the same approach on single-cell datasets. Recently, the CROP-Seq methodology has been introduced [17], allowing for the measurement of cell-specific transcriptome-wide gene expression and mutations induced by CRISPR-Cas9 [50], thanks to the concurrent sequencing of CRISPR-Cas9 guide RNAs. We therefore tested the capability of gbm models to predict mutations using gene expression variables in two independent single-cell datasets. The first dataset (dubbed “Datlinger”) was extracted from the Jurkat cell line derived from human T lymphocytes [17], The second one (dubbed “Shifrut”) derive from primary unstimulated T cells from a human donor [51]. We removed cell UMI (Unique Molecular Identifier) counts and cell cycle as common confounding effects of single cell datasets [52] (Figure S10). We generated a regulatory transcription network using ARACNe-AP on the RNA-Seq Cancer Cell Line Encyclopedia dataset (CCLE [53]), which comprises 1021 distinct human cell lines. Using the CCLE network, we aggregated gene expression from the single cell datasets using the VIPER algorithm and implemented the resulting TF-centered VIPER activity profiles to build prediction models for the Crop-Seq detected mutations. Parallelly, we built models using un-aggregated VST-normalized gene expression data. Our results show that gbm models based on VIPER activity variables globally achieve a significantly higher performance in both the Datlinger (p=8.0xl0^-85^) and Shifrut datasets (p=2.2xl0^-117^) when compared with models obtained from gene expression data (Figure 6). For specific mutations (TUBB gene, CDKN1B) the VIPER aggregation based on CCLE ARACNe networks seem to be particularly beneficial to increase the performance of mutation prediction models based on gene expression, while for a few mutations, such as RUNX1, the CCLE-based networks significantly decrease the model performance.

**Figure 6.**
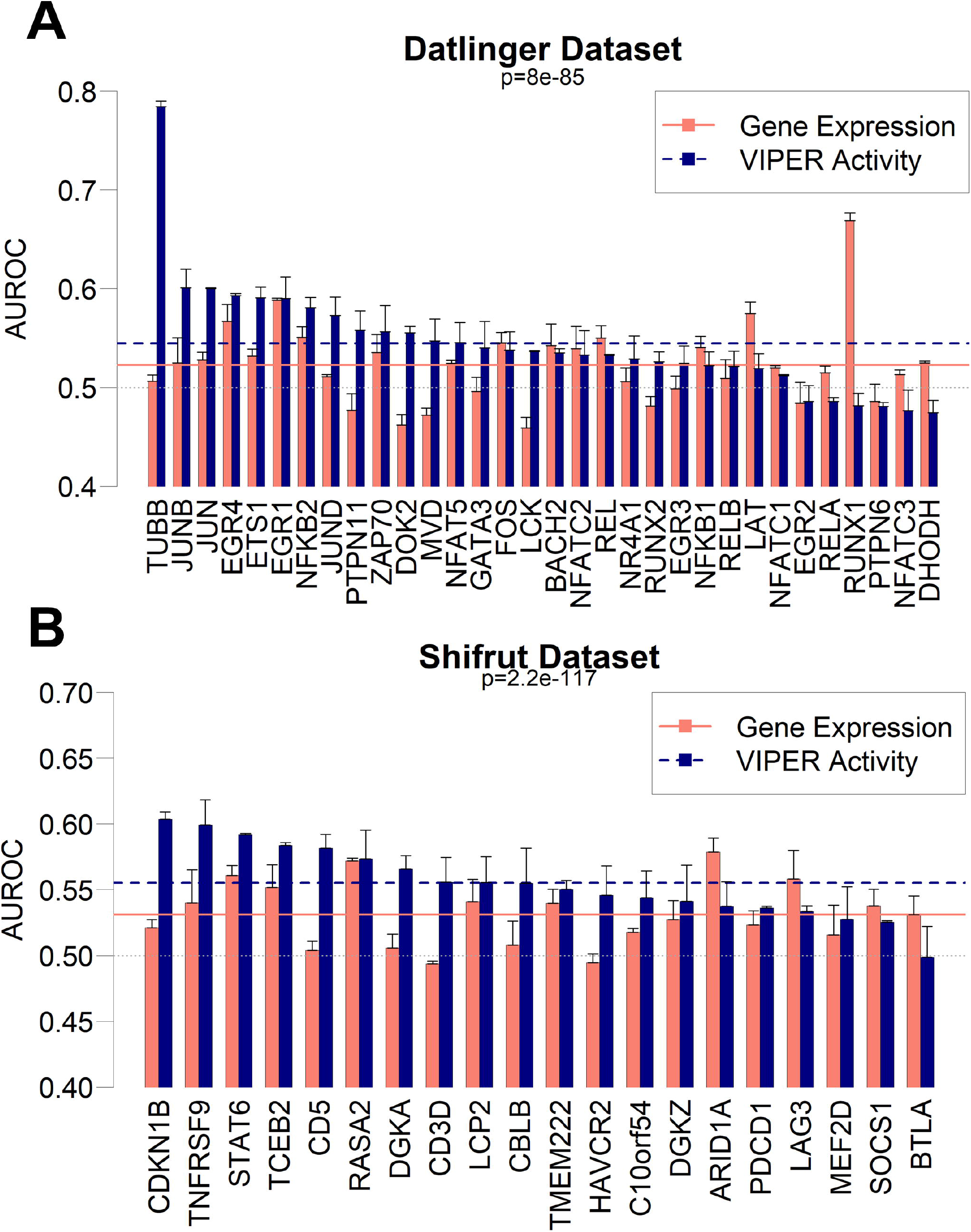
Performance as AUROC of gbm models to predict mutations in CROP-Seq datasets using gene expression (red bars) and VIPER activity (blue bars) derived from CCLE expression data in Datlinger (A) and Shifrut (B) datasets. The p-value of paired Wilcoxon tests between all VIPER and Expression AUROCs in each dataset is reported, as well as the average of all expression models (red solid line) and all VIPER activity models (blue dashed line). Error bars report the standard deviation of 100 AUROCs generated from multiple partitioning of training/test sets.

## Discussion

In this paper, we tested a framework to investigate the complex relationships between genetic events and transcriptional deregulation through machine learning approaches. We demonstrated as a generalized proof-of-principle that genomic alterations can be modeled by gene expression across several human cancers through several machine learning algorithms, and specifically that a gradient boost modeling approach seems optimal for the task. In the process, we generated a collection of models for each genomic alteration in each cancer context, showing that the best predicted alterations are not necessarily targeting known oncogenes or tumor suppressors. Interestingly, we show how the aggregation of gene expression profiles in groups of coexpressed genes, via the ARACNe/VIPER or WGCNA methods, makes the models more robust and more resistant to perturbations such as gaussian noise or artificial downsampling. Finally, we have shown how the same aggregation principle can have beneficial effects in predicting the presence of mutations in intrinsically noisy scenarios, both with artificial noise introduction and read reduction. At the same time, we have shown that expression-based mutation prediction can be modeled out in single-cell sequencing contexts, which can be considered as real cases of noisy datasets. The capability of predicting mutations based on single-cell RNA-Seq is however reduced when compared to datasets derived from pooled cells sequencing, as those provided by the TCGA dataset: the average performances of TCGA models (Figure 3) generally rest on a range between 0.6 and 0.9 area under the ROC curve, while the performance of CROP-Seq models fall on an average value of 0.55 (Figure 6).

The performance of gene aggregation methods has been tested before for sample clustering in RNA-Seq read reduction scenarios [15,49], but never in this specific task nor in a pancancer or a single-cell context. As a principle, the usage of robust averages of pre-defined co-expressed genes can be applied in any context where reliability of gene expression data is necessary, from differential expression to pathway enrichment analyses. The notion that relationships between genomic alterations and gene expression profiles can be robustly modelled across different cancer scenarios, as well as in single-cell and noisy contexts, can have important repercussions in diagnostics and quantification studies of heterogeneous cell populations, where theoretically a single quantitative expression experiment can be used to predict the presence or absence of a mutation.

## Materials and Methods

### Data processing

We obtained raw expression counts, mutation and CNV raw data from TCGA using the Firehose portal (gdac.broadinstitute.org). Raw counts were normalized using Variance Stabilizing Transformation as described before [54], Somatic mutations not changing the aminoacid sequence of the protein product were discarded. We flagged genes blacklisted by the MutSig project [22], such as TTN, ORs, MUCs as false positives, and removed them from further analysis (except the most mutated in the pan-cancer dataset, shown in Figure S1). CNV tracks were associated to the targeted gene using the GenomicRanges R package [55]. Gene-centered CNVs were then associated to the expression profile of the gene itself. Genes affected by a CNV in more than 10 samples were used in the rest of the analysis. Samples with more than 0.5% of the genes in the genome somatically amplified, deleted or mutated were deemed “hypermodified” and the total number was shown in Figure 1 bottom bars.

Clustering analysis was carried out on the TCGA tumor samples using the expression profiles of 1172 Transcription Factors defined by Gene Ontology terms “transcription factor activity, sequence-specific DNA binding” (GO:0003700) and “nuclear location” (GO:0005634) [56].

The dataset expression profiles were visualized after TSNE transformation [19] with 1000 iterations using a 2D kernel density estimate for coloring different tumor types [57], Oncogenes and Tumor Suppressor genes were obtained from the COSMIC Cancer Gene Census in October 2018 [42].

### Modeling

We used the R caret package [18] v 6.0-81 as the platform to run all our predictive models in a standardized and reproducible way. Default parameters for model training were used. Binary classifiers were built to predict the presence/absence of mutation, amplification and deletion events. The CNV value provided by TCGA corresponds to log2(tumor coverage) – genomic median coverage. The threshold for amplification/deletion presence was set to 0.5.

Data partitioning was performed once for each tumor type, with 75% of the samples used for training and 25% for test purposes. Training was performed using 10-fold Cross Validation. Recursive Feature Elimination was carried out by the default caret implementation on the 10,000 highest variance gene expression tracks. The algorithms used (and R packages implementing theme) were:

- Bayesian Generalized Linear Model (bayesglm)
- Tree Models from Genetic Algorithms (evtree)
- Gradient Boost Modeling (gbm)
- Generalized Linear Model (glm)
- k-Nearest Neighbors (kknn)
- Linear Discriminant Analysis (lda)
- Neural Networks (mxnet)
- Neural Networks with Feature Extraction (pcaNNet)
- Random Forest (rf)
- Linear Support Vector Machine (svmLinear)
- Radial Support Vector Machine (svmRadial)

In order to reduce information from the gene expression profiles, we adopted two strategies. The first, shown e.g. in Figure 4B, adds random gaussian noise to the expression tracks, with a variable standard deviation (indicated as “Gaussian Noise Level”). Each model run after noise addition was run 100 times to allow for various data partitions. The second strategy (Figure 5) reduced the number of reads mapped to each gene in order to obtain expression samples with decreased total gene counts. In order to do so, we applied to each gene in each sample a downsampling factor sample from a beta distribution:

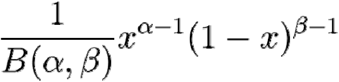

Where B is the Beta function, acting as a normalization constant, x is the raw gene expression count in a particular sample, α is the first shape parameter and β the second shape parameter. In order to reduce the total sample coverage to the desired level, β is set to 0.1 and α is set to:

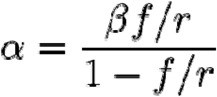

Where *f* is the desired number of reads and r is the total number of reads in the sample. A real case example of this beta distribution is shown in Figure S9.

### Aggregation algorithms

We used ARACNe-AP [46] to generate TF-centered networks on each of the VST-normalized TCGA expression datasets. TFs were selected via Gene Ontology as described before, with p-value for each network edge set to 10^-8^. ARACNe networks were then used to obtain an aggregated value of TF activity for each sample using the VIPER algorithm [15] which reports the collective gene expression level changes of each TF-centered network vs. the mean expression of each gene in the dataset. Only TF networks with at least 10 genes (excluding the TF) were included.

WGCNA clusters of genes were constructed using the wcgna package [45] with default parameters and minimum network size set to 10 To obtain a robust median expression value for each WGCNA cluster in each sample we used Tukey’s Biweight function as implemented by the R *affy* package [58].

### Single Cell analysis

We generated TF regulatory networks using ARACNe-AP as described before on the CCLE dataset available at https://portals.broadinstitute.org/ccle/data, raw counts version 2018-09-29, normalized by Variance-Stabilizing Transformation [58].

We downloaded raw RNA-Seq counts and gRNA mutation data from single-cell CROP-Seq datasets, specifically: 1) the Datlinger dataset available on Gene Expression Omnibus (GEO) series GSE92872 [17] and 2) the Shifrut dataset was obtained from a healthy donor and is available as raw counts and cell-sepcific gRNA from GEO sample GSM3375483 [51]. Both single cell CROP-Seq datasets were normalized using the R package Seurat with default parameters, as follows: a global-scaling normalization method (“LogNormalize”) was applied on raw gene counts for each cell, then the values were multiplied by a scale factor (10,000 by default) and the results were log-normalized. These values were then regressed by two variables: UMI counts and cell cycle, using cell cycle markers from [52], As an example of the Seurat regression, the TSNE representation of the Datlinger dataset before and after normalization clearly shows the removal of cell-cycle bias effects (Figure S10).

Gradient boost modelling (gbm) was applied to each CROP-Seq dataset by aggregating cells carrying mutations on the same genes and using wild-type cells as control. Performance of gbm models using VIPER and expression variables was compared using a two-tailed Wilcoxon test on 100 repetitions of training/test set splits before cross-validation for model testing [59].

### Methods Availability

All code used to generate the analysis and the figures of this paper is available in the online materials as Supplementary Code.

## Acknowledgments

We acknowledge the CINECA award (projects HP10CB1R7T and HP10CPQJBV) under the ISCRA initiative, for the support and availability of high-performance computing resources. We also thank Lupo Giorgi, Dr. Luca Pestarino and Jordan Pflugh Kraft for the fruitful discussions.

## Supporting Information Legends

**Figure SI.** Table of most somatically mutated genes across TCGA tumor samples, in terms of number of samples where the gene is somatically mutated with altered protein product sequence. This table includes also MutSig-blacklisted genes (in grey) such as Titin (TTN), Obscurin (OBSCN) and Mucin genes.

**Figure S2.** Relationship between alteration models and alteration frequency in the Pancancer dataset, for mutations (left), amplifications (center) and deletions (right).

**Figure S3.** Performance of Pan-cancer alterations models globally (left) and for MutSig genes, COSMIC oncogenes and COSMIC tumor suppressors. The Y-Axis indicates rank-transformed AUROC values. Asterisks indicate a significant (<0.01) difference between a distribution and the global “Other Genes” distribution according to Two-tailed Wilcoxon tests.

**Figure S4.** ROC curves for gbm TP53 models in Breast Cancer, using original expression data, VIPER aggregation (TF “activity”) and WGCNA aggregation (robust tukey biweight average of clusters).

**Figure S5.** AUROCs of EGFR amplication gbm prediction models in Glioblastoma with increasing noise, calculated using gene expression (black line) or aggregated gene expression using the WGCNA (green line) or VIPER (red line) algorithms.

**Figure S6.** AUROCs of EGFR amplication gbm prediction models in Lung Squamous Carcinoma (LUSC) with increasing noise, calculated using gene expression (black line) or aggregated gene expression using the WGCNA (green line) or VIPER (red line) algorithms.

**Figure S7.** AUROCs of PVT1 amplication gbm prediction models in Ovarian Cancer with increasing noise, calculated using gene expression (black line) or aggregated gene expression using the WGCNA (green line) or VIPER (red line) algorithms.

**Figure S8.** AUROCs of PTEN deletion gbm prediction models in Sarcoma with increasing noise, calculated using gene expression (black line) or aggregated gene expression using the WGCNA (green line) or VIPER (red line) algorithms.

**Figure S9.** Beta distribution used to down-sample the 43.8M reads breast cancer sample TCGA-A1-A0SB-01 to 10M reads. The grey line shows the ratio between the target coverage and the original coverage.

**Figure S10.** TSNE representation of the Datlinger CROP-Seq dataset before (A) and after (B) removal of cell-cycle specific markers. Colors indicated the predicted cell cycle phase according to the Seurat pipeline [60].

**Supplementary Table S1.** AUROCs for each event in the Pan-Cancer TCGA dataset (24 tumor types with at least 100 samples with co-measured genomic and expression data. The Sheet name indicates the tumor type and genomic alteration type (mut: somatic mutation, amp: amplification, del: deletion).

**Supplementary Code.** R and bash code snippets used in this study.

