## Supplementary Code for "Pan-Cancer and Single-Cell modelling of genomic alterations through gene expression": 000a_getRNASeq_Clinical.R

```
### Get stuff from http://gdac.broadinstitute.org/
### Nice guidelines on https://www.biostars.org/p/153013/
### https://confluence.broadinstitute.org/display/GDAC/Download

cd $DROPBOX/projects/pancancer/rawtums010
wget http://gdac.broadinstitute.org/runs/code/firehose_get_latest.zip
unzip firehose_get_latest.zip
./firehose_get --help
./firehose_get -c

# ./firehose_get analyses latest # 2016_01_28 run
cd $DROPBOX/projects/pancancer/rawtums010
./firehose_get -o Merge_Clinical.Level_1 data latest # 2016_07_15 run
./firehose_get -o Merge_rnaseqv2__illuminahiseq_rnaseqv2__unc_edu__Level_3__RSEM_genes_normalized__data.Level_3 data latest
./firehose_get -o Merge_rnaseqv2__illuminahiseq_rnaseqv2__unc_edu__Level_3__RSEM_genes__data.Level_3 data latest

# The loop to extract these very long names
cd $DROPBOX/projects/pancancer/rawtums010
for tum in ACC BLCA BRCA CESC CHOL COAD DLBC ESCA GBM HNSC KICH KIRC KIRP LAML LGG LIHC LUAD LUSC MESO OV PAAD PCPG PRAD READ SARC SKCM STAD TGCT THCA THYM UCEC UCS UVM
do

tarname=stddata__2016_07_15/${tum}/20160715/gdac.broadinstitute.org_${tum}.Merge_rnaseqv2__illuminahiseq_rnaseqv2__unc_edu__Level_3__RSEM_genes__data.Level_3.2016071500.0.0.tar.gz
tar xvzf $tarname
cd gdac.broadinstitute.org_${tum}.Merge_rnaseqv2__illuminahiseq_rnaseqv2__unc_edu__Level_3__RSEM_genes__data.Level_3.2016071500.0.0
mv ${tum}.rnaseqv2__illuminahiseq_rnaseqv2__unc_edu__Level_3__RSEM_genes__data.data.txt ../${tum}_RSEM_genes.txt
cd ..

tarname=stddata__2016_07_15/${tum}/20160715/gdac.broadinstitute.org_${tum}.Merge_rnaseqv2__illuminahiseq_rnaseqv2__unc_edu__Level_3__RSEM_genes_normalized__data.Level_3.2016071500.0.0.tar.gz
tar xvzf $tarname
cd gdac.broadinstitute.org_${tum}.Merge_rnaseqv2__illuminahiseq_rnaseqv2__unc_edu__Level_3__RSEM_genes_normalized__data.Level_3.2016071500.0.0
mv ${tum}.rnaseqv2__illuminahiseq_rnaseqv2__unc_edu__Level_3__RSEM_genes_normalized__data.data.txt ../${tum}_RSEM_genes_normalized.txt
cd ..

tarname=stddata__2016_07_15/${tum}/20160715/gdac.broadinstitute.org_${tum}.Merge_Clinical.Level_1.2016071500.0.0.tar.gz
tar xvzf $tarname
cd gdac.broadinstitute.org_${tum}.Merge_Clinical.Level_1.2016071500.0.0
mv ${tum}.merged_only_clinical_clin_format.txt ../${tum}_clinical.txt
cd ..
done


#### Format nicely in R
if(TRUE){
    raw<-matrix(c(
        "ACC","Adrenocortical Carcinoma",
        "BLCA","Bladder Urothelial Carcinoma",
        "BRCA","Breast Invasive Carcinoma",
        "CESC","Cervical Squamous Cell carcinoma",
        "CHOL","Cholangiocarcinoma",
        "COAD","Colon Adenocarcinoma",
        "DLBC","Diffuse Large B-cell Lymphoma",
        "ESCA","Esophageal Carcinoma",
        "GBM","Glioblastoma Multiforme",
        "HNSC","Head and Neck Squamous Cell Carcinoma",
        "KICH","Kidney Chromophobe",
        "KIRC","Kidney Renal Clear Cell Carcinoma",
        "KIRP","Kidney Renal Papillary Cell Carcinoma",
        "LAML","Acute Myeloid Leukemia",
        "LGG","Brain Lower Grade Glioma",
        "LIHC","Liver Hepatocellular Carcinoma",
        "LUAD","Lung Adenocarcinoma",
        "LUSC","Lung Squamous Cell Carcinoma",
        "MESO","Mesothelioma",
        "OV","Ovarian Serous Cystadenocarcinoma",
        "PAAD","Pancreatic Adenocarcinoma",
        "PCPG","Pheochromocytoma and Paraganglioma",
        "PRAD","Prostate Adenocarcinoma",
        "READ","Rectum Adenocarcinoma",
        "SARC","Sarcoma",
        "SKCM","Skin Cutaneous Melanoma",
        "STAD","Stomach Adenocarcinoma",
        "TGCT","Testicular Germ Cell Tumors",
        "THCA","Thyroid Carcinoma",
        "THYM","Thymoma",
        "UCEC","Uterine Corpus Endometrial Carcinoma",
        "UCS","Uterine Carcinosarcoma",
        "UVM","Uveal Melanoma"
    ),ncol=2,byrow = TRUE
    )
    tums<-setNames(raw[,2],raw[,1])
}
# Generate files
for(tum in names(tums)){
    message("Doing ",tum)
    dir.create(paste0("tums010/",tum))
    
    # Raw counts
    fname<-paste0("rawtums010/",tum,"_RSEM_genes.txt")
    superaw<-read.delim(fname,as.is=TRUE,row.names=1)
    sampleids<-gsub("\\.","-",substr(colnames(superaw),1,16))
    rawcounts<-superaw[2:nrow(superaw),(0:((ncol(superaw)/3)-1)*3)+1]
    colnames(rawcounts)<-sampleids[(0:((ncol(superaw)/3)-1)*3)+1]
    rownames(rawcounts)<-gsub(".+\\|","",rownames(rawcounts))
    for(j in 1:ncol(rawcounts)){
        rawcounts[,j]<-round(as.numeric(rawcounts[,j]))
    }
    save(rawcounts,file=paste0("tums010/",tum,"/",tum,"-rawcounts.rda"))
    rm(superaw)
    
    # RSEM normalized
    fname<-paste0("rawtums010/",tum,"_RSEM_genes_normalized.txt")
    superaw<-read.delim(fname,as.is=TRUE,row.names=1)
    normalized<-superaw[2:nrow(superaw),]
    colnames(normalized)<-gsub("\\.","-",substr(colnames(normalized),1,16))
    rownames(normalized)<-gsub(".+\\|","",rownames(normalized))
    save(normalized,file=paste0("tums010/",tum,"/",tum,"-normalized.rda"))
    rm(superaw)
    
    # Clinical
    fname<-paste0("rawtums010/",tum,"_clinical.txt")
    clinical<-as.data.frame(t(read.delim(fname,header=TRUE,row.names=1,sep="\t",as.is=TRUE)))
    rownames(clinical)<-clinical$IDs<-toupper(clinical$patient.bcr_patient_barcode)
    save(clinical,file=paste0("tums010/",tum,"/",tum,"-clinical.rda"))
    
    rm(clinical,rawcounts,normalized)
}


# Survival
library(survival)
for(tum in names(tums)){
    message("Doing ",tum)
    load(paste0("tums010/",tum,"/",tum,"-clinical.rda"))
    
    
    # Get survival info
    rawsurv<-clinical[,colnames(clinical)%in%c(
        "patient.days_to_birth",
        "patient.days_to_last_followup",
        "patient.days_to_death"
    )]
    for(j in 1:3){
        rawsurv[,j]<-as.integer(as.character(rawsurv[,j]))
    }
    
    # Create a survival object
    rawsurvtime<-rawsurv[,colnames(rawsurv)%in%c(
        "patient.days_to_last_followup",
        "patient.days_to_death"
    )
    ]
    rawsurvtime[is.na(rawsurvtime)]<-0
    eventTime<-apply(rawsurvtime,1,function(x){
        as.numeric(max(as.numeric(x),na.rm=TRUE))
    })
    survival<-Surv(
        time=eventTime,
        event=clinical$patient.vital_status=="dead"
    )
    rownames(survival)<-rownames(clinical)
    save(survival,file=paste0("tums010/",tum,"/",tum,"-survival.rda"))
}


# VST Normalization
library("DESeq")
for(tum in names(tums)){
    message("Doing ",tum)
    load(paste0("tums010/",tum,"/",tum,"-rawcounts.rda"))
    
    # Load count table into a cds object
    conditions<-factor(colnames(rawcounts))
    cds<-newCountDataSet(rawcounts,conditions)
    cds<-estimateSizeFactors(cds)
    
    ### Variance Stabilizing Transformation
    #This function calculates a variance stabilizing transformation
    #(VST) from the fitted dispersion-mean relation(s) and then
    #transforms the count data (normalized by division by the size
    #factor), yielding a matrix of values which are now approximately
    #homoskedastic. This is useful as input to statistical analyses
    #requiring homoskedasticity.
    
    #RNA-Seq measurements of expression differ from those of other platforms
    #in several ways, but one of the most important is that even after normalizing
    #for length and total read yield (RPKM) the variance on a gene's expression across
    #replicates is largely a function of its overall expression (at least with current protocols).
    
    cdsBlind<-estimateDispersions(cds,method="blind")
    vsd<-varianceStabilizingTransformation(cdsBlind)
    expmat<-exprs(vsd)
    
    save(expmat,file=paste0("tums010/",tum,"/",tum,"-expmat.rda"))
    rm(expmat,rawcounts)
}

# Files for ARACNe
for(tum in names(tums)){
    message("Doing ",tum)
    load(paste0("tums010/",tum,"/",tum,"-expmat.rda"))
    cat("gene\t",file=paste0("tums010/",tum,"/",tum,"-expmat.dat"))
    write.table(expmat,
                file=paste0("tums010/",tum,"/",tum,"-expmat.dat"),
                col.names=TRUE,
                sep="\t",
                quote=FALSE,
                append=TRUE
    )
}

cd $DROPBOX/projects/pancancer
scp -r tums010/:/gpfs/work/IscrC_tumornet/pancancer/
    
    
    
    
    
```
