## Supplementary Code for "Pan-Cancer and Single-Cell modelling of genomic alterations through gene expression": 000b_getMutations.R

```
### Get stuff from http://gdac.broadinstitute.org/
### Nice guidelines on https://www.biostars.org/p/153013/
### https://confluence.broadinstitute.org/display/GDAC/Download

### Coding Mutations
cd $DROPBOX/projects/pancancer/rawtums010
wget http://gdac.broadinstitute.org/runs/code/firehose_get_latest.zip
unzip firehose_get_latest.zip
./firehose_get --help
./firehose_get -c
#./firehose_get -tasks data latest prad # 2016_07_15 run
./firehose_get -o Mutation_Packager_Calls.Level_3 data latest # 2016_07_15 run
./firehose_get -o Mutation_Packager_Raw_Calls.Level_3 data latest coad read # 2016_07_15 run

# Unzip
cd $DROPBOX/projects/pancancer/rawtums010
for tum in ACC BLCA BRCA CESC CHOL COAD DLBC ESCA GBM HNSC KICH KIRC KIRP LAML LGG LIHC LUAD LUSC MESO OV PAAD PCPG PRAD READ SARC SKCM STAD TGCT THCA THYM UCEC UCS UVM
do
echo $tum
tarname=stddata__2016_07_15/${tum}/20160715/gdac.broadinstitute.org_${tum}.Mutation_Packager_Calls.Level_3.2016071500.0.0.tar.gz
tar xvzf $tarname
done

for tum in COAD READ
do
echo $tum
tarname=stddata__2016_07_15/${tum}/20160715/gdac.broadinstitute.org_${tum}.Mutation_Packager_Raw_Calls.Level_3.2016071500.0.0.tar.gz
tar xvzf $tarname
done


# Format nicely in R
if(TRUE){
raw<-matrix(c(
    "ACC","Adrenocortical Carcinoma",
    "BLCA","Bladder Urothelial Carcinoma",
    "BRCA","Breast Invasive Carcinoma",
    "CESC","Cervical Squamous Cell carcinoma",
    "CHOL","Cholangiocarcinoma",
    "COAD","Colon Adenocarcinoma",
    "DLBC","Diffuse Large B-cell Lymphoma",
    "ESCA","Esophageal Carcinoma",
    "GBM","Glioblastoma Multiforme",
    "HNSC","Head and Neck Squamous Cell Carcinoma",
    "KICH","Kidney Chromophobe",
    "KIRC","Kidney Renal Clear Cell Carcinoma",
    "KIRP","Kidney Renal Papillary Cell Carcinoma",
    "LAML","Acute Myeloid Leukemia",
    "LGG","Brain Lower Grade Glioma",
    "LIHC","Liver Hepatocellular Carcinoma",
    "LUAD","Lung Adenocarcinoma",
    "LUSC","Lung Squamous Cell Carcinoma",
    "MESO","Mesothelioma",
    "OV","Ovarian Serous Cystadenocarcinoma",
    "PAAD","Pancreatic Adenocarcinoma",
    "PCPG","Pheochromocytoma and Paraganglioma",
    "PRAD","Prostate Adenocarcinoma",
    "READ","Rectum Adenocarcinoma",
    "SARC","Sarcoma",
    "SKCM","Skin Cutaneous Melanoma",
    "STAD","Stomach Adenocarcinoma",
    "TGCT","Testicular Germ Cell Tumors",
    "THCA","Thyroid Carcinoma",
    "THYM","Thymoma",
    "UCEC","Uterine Corpus Endometrial Carcinoma",
    "UCS","Uterine Carcinosarcoma",
    "UVM","Uveal Melanoma"
),ncol=2,byrow = TRUE
)
tums<-setNames(raw[,2],raw[,1])
rm(raw)
} # Tumor Acronyms

# Generate raw mutations files
for(tum in names(tums)){
    message("Doing ",tum)
    dirname<-paste0("rawtums010/gdac.broadinstitute.org_",tum,".Mutation_Packager_Calls.Level_3.2016071500.0.0")
    files<-dir(dirname)
    files<-setdiff(files,"MANIFEST.txt")
    if(length(files)<1){next}
    # Create a global MAF
    rawmuts<-read.delim(paste0(dirname,"/",files[1]),as.is=TRUE)
    for(i in 2:length(files)){
        raw<-read.delim(paste0(dirname,"/",files[i]),as.is=TRUE)
        rawmuts<-rbind(rawmuts,raw)
    }
    save(rawmuts,file=paste0("tums010/",tum,"/",tum,"-rawmuts.rda"))
    
    # Add raw mutation calls for more samples
    dirname<-paste0("rawtums010/gdac.broadinstitute.org_",tum,".Mutation_Packager_Raw_Calls.Level_3.2016071500.0.0")
    files2<-dir(dirname)
    files2<-setdiff(files2,"MANIFEST.txt")
    files2<-setdiff(files2,files)
    if(length(files2)<1){next}
    for(i in 1:length(files2)){
        raw<-read.delim(paste0(dirname,"/",files2[i]),as.is=TRUE)
        int<-intersect(colnames(raw),colnames(rawmuts))
        rawmuts<-rbind(rawmuts[,int],raw[,int])
    }
    save(rawmuts,file=paste0("tums010/",tum,"/",tum,"-rawmuts.rda"))
    
}


# Generate coding mutations files
source("../shared/functions/geneids.R")
for(tum in names(tums)){
    message("Doing ",tum)
    input<-paste0("tums010/",tum,"/",tum,"-rawmuts.rda")
    if(file.exists(input)){
        load(input)
        rawsnp<-rawmuts
        
        # Remove entrezs equaling 0
        zeroids<-which(rawsnp[,"Entrez_Gene_Id"]==0)
        if(length(zeroids)>0){
            rawsnp[zeroids,"Entrez_Gene_Id"]<-sym2eg(rawsnp[zeroids,"Hugo_Symbol"])
            zeroids<-grep("unknown",rawsnp[,"Entrez_Gene_Id"])
            if(length(zeroids)>0){rawsnp<-rawsnp[-zeroids,]}
            rawsnp<-rawsnp[rownames(rawsnp)!="0",]
        }

        # Shorten sample id
        rawsnp[, "Tumor_Sample_Barcode"] <- substr(rawsnp[, "Tumor_Sample_Barcode"], 1, 15)

        # Remove Silent mutations
        rawsnp <- rawsnp[!(rawsnp[, "Variant_Classification"] %in% c("Silent", "RNA")), ]
        
        # Summarize genes mutated more than once in the same patient
        rawsnp <- rawsnp[!duplicated(paste(rawsnp[,"Tumor_Sample_Barcode"], rawsnp[,"Entrez_Gene_Id"], sep="-")), ]
        
        # Format the table more nicely (1/0 format)
        snp<-matrix(0,
                    ncol=length(unique(rawsnp[,"Tumor_Sample_Barcode"])),
                    nrow=length(unique(rawsnp[,"Entrez_Gene_Id"]))
        )
        rownames(snp)<-as.character(unique(rawsnp[,"Entrez_Gene_Id"]))
        colnames(snp)<-unique(rawsnp[,"Tumor_Sample_Barcode"])
        
        # Assign 1/0 to each mutation/patient combination
        for(i in as.character(unique(rawsnp$Entrez_Gene_Id))){
            snpsamples<-rawsnp$Tumor_Sample_Barcode[as.character(rawsnp$Entrez_Gene_Id)==i]
            snpsamples<-snpsamples[!is.na(snpsamples)]
            if(length(snpsamples)>0){
                snp[i,snpsamples]<-1
            }
        }
        mutations<-snp
        save(mutations,file=paste0("tums010/",tum,"/",tum,"-mutations.rda"))
        rm(rawsnp,mutations,snp)
    }
}
```
