## Supplementary Code for "Pan-Cancer and Single-Cell modelling of genomic alterations through gene expression": 000c_getCNVs.R

```
### Get stuff from http://gdac.broadinstitute.org/
### Nice guidelines on https://www.biostars.org/p/153013/
### https://confluence.broadinstitute.org/display/GDAC/Download

### Copy Number Variations
cd $DROPBOX/projects/pancancer/rawtums010
wget http://gdac.broadinstitute.org/runs/code/firehose_get_latest.zip
unzip firehose_get_latest.zip
./firehose_get --help
./firehose_get -c
./firehose_get -tasks data latest prad # 2016_07_15 run
./firehose_get -o Merge_cna__illuminahiseq_dnaseqc__hms_harvard_edu__Level_3__segmentation__seg data latest # 2016_07_15 run
./firehose_get -o Merge_snp__genome_wide_snp_6__broad_mit_edu__Level_3__segmented_scna_minus_germline_cnv_hg19__seg data latest # 2016_07_15 run


# The loop to extract these very long names
cd $DROPBOX/projects/pancancer/rawtums010
for tum in ACC BLCA BRCA CESC CHOL COAD DLBC ESCA GBM HNSC KICH KIRC KIRP LAML LGG LIHC LUAD LUSC MESO OV PAAD PCPG PRAD READ SARC SKCM STAD TGCT THCA THYM UCEC UCS UVM
do

tarname=stddata__2016_07_15/${tum}/20160715/gdac.broadinstitute.org_${tum}.Merge_cna__illuminahiseq_dnaseqc__hms_harvard_edu__Level_3__segmentation__seg.Level_3.2016071500.0.0.tar.gz
tar xvzf $tarname
cd gdac.broadinstitute.org_${tum}.Merge_cna__illuminahiseq_dnaseqc__hms_harvard_edu__Level_3__segmentation__seg.Level_3.2016071500.0.0
mv ${tum}.cna__illuminahiseq_dnaseqc__hms_harvard_edu__Level_3__segmentation__seg.seg.txt ../${tum}_cna_dnaseq.txt
cd $DROPBOX/projects/pancancer/rawtums010

tarname=stddata__2016_07_15/${tum}/20160715/gdac.broadinstitute.org_${tum}.Merge_snp__genome_wide_snp_6__broad_mit_edu__Level_3__segmented_scna_minus_germline_cnv_hg19__seg.Level_3.2016071500.0.0.tar.gz
tar xvzf $tarname
cd gdac.broadinstitute.org_${tum}.Merge_snp__genome_wide_snp_6__broad_mit_edu__Level_3__segmented_scna_minus_germline_cnv_hg19__seg.Level_3.2016071500.0.0
mv ${tum}.snp__genome_wide_snp_6__broad_mit_edu__Level_3__segmented_scna_minus_germline_cnv_hg19__seg.seg.txt ../${tum}_cna_snparray.txt
cd $DROPBOX/projects/pancancer/rawtums010

done


# Format nicely in R
setwd("D:/Dropbox/projects/pancancer")

library(GenomicFeatures)
library(Biobase)
library(TxDb.Hsapiens.UCSC.hg19.knownGene)
hg19genes<-transcriptsBy(TxDb.Hsapiens.UCSC.hg19.knownGene, by="gene")


if(TRUE){
    raw<-matrix(c(
        "ACC","Adrenocortical Carcinoma",
        "BLCA","Bladder Urothelial Carcinoma",
        "BRCA","Breast Invasive Carcinoma",
        "CESC","Cervical Squamous Cell carcinoma",
        "CHOL","Cholangiocarcinoma",
        "COAD","Colon Adenocarcinoma",
        "DLBC","Diffuse Large B-cell Lymphoma",
        "ESCA","Esophageal Carcinoma",
        "GBM","Glioblastoma Multiforme",
        "HNSC","Head and Neck Squamous Cell Carcinoma",
        "KICH","Kidney Chromophobe",
        "KIRC","Kidney Renal Clear Cell Carcinoma",
        "KIRP","Kidney Renal Papillary Cell Carcinoma",
        "LAML","Acute Myeloid Leukemia",
        "LGG","Brain Lower Grade Glioma",
        "LIHC","Liver Hepatocellular Carcinoma",
        "LUAD","Lung Adenocarcinoma",
        "LUSC","Lung Squamous Cell Carcinoma",
        "MESO","Mesothelioma",
        "OV","Ovarian Serous Cystadenocarcinoma",
        "PAAD","Pancreatic Adenocarcinoma",
        "PCPG","Pheochromocytoma and Paraganglioma",
        "PRAD","Prostate Adenocarcinoma",
        "READ","Rectum Adenocarcinoma",
        "SARC","Sarcoma",
        "SKCM","Skin Cutaneous Melanoma",
        "STAD","Stomach Adenocarcinoma",
        "TGCT","Testicular Germ Cell Tumors",
        "THCA","Thyroid Carcinoma",
        "THYM","Thymoma",
        "UCEC","Uterine Corpus Endometrial Carcinoma",
        "UCS","Uterine Carcinosarcoma",
        "UVM","Uveal Melanoma"
    ),ncol=2,byrow = TRUE
    )
    tums<-setNames(raw[,2],raw[,1])
    rm(raw)
}


cnvProcess <- function(input) {
    # Fix X and Y
    cnvd<-input
    cnvd[,2]<-paste0("chr",cnvd[,2])
    cnvd[cnvd[,2]=="chr23",2]<-"chrX"
    cnvd[cnvd[,2]=="chr24",2]<-"chrY"
    
    # The next cryptic block converts chromosome coordinates into entrez ids
    cnvd1<-GRanges(
        seqnames=cnvd$Chromosome,
        ranges=IRanges(start=cnvd$Start, end=cnvd$End),
        strand=rep("*", length(cnvd$Chromosome)),
        sample=cnvd$Sample
    )
    # Find overlaps between CNV regions and genes
    overlap<-as.matrix(findOverlaps(cnvd1, hg19genes))
    overlap[,2]<-names(hg19genes)[overlap[,2]]
    overlap<-cbind(overlap,
                   cnvd$Segment_Mean[as.numeric(overlap[,1])],
                   cnvd$Sample[as.numeric(overlap[,1])]
    )
    overlap<-overlap[order(abs(as.numeric(overlap[,3])),decreasing=T),]
    colnames(overlap)<-c("SegmentNr","EntrezId","CNVlevel","Sample")
    overlap<-overlap[!duplicated(paste0(overlap[,2],"_",overlap[,4])),]
    rm(cnvd,cnvd1)
    
    # CNV object: gene-level CN levels
    genes<-unique(overlap[,2])
    samples<-unique(overlap[,4])
    cnv<-matrix(0,nrow=length(genes),ncol=length(samples))
    rownames(cnv)<-genes
    colnames(cnv)<-samples
    
    # Loop
    pb<-txtProgressBar(0,nrow(overlap),style=3)
    for(i in 1:nrow(overlap)){
        gene<-overlap[i,2]
        cnvlevel<-overlap[i,3]
        sample<-overlap[i,4]
        cnv[gene,sample]<-as.numeric(cnvlevel)
        setTxtProgressBar(pb,i)
    }
    return(cnv)
}


# Generate files
for(tum in names(tums)){
    message("Doing ",tum)
    
    # Files (not used, but just to know we have them)
    fname1<-paste0("rawtums010/",tum,"_cna_dnaseq.txt")
    fname2<-paste0("rawtums010/",tum,"_cna_snparray.txt")
    
    if(file.exists(fname1)){
        raw1<-read.delim(fname1,as.is=TRUE)
        if(file.exists(fname2)){
            raw2<-read.delim(fname2,as.is=TRUE)
            new<-setdiff(raw1[,1],raw2[,1])
            raw1<-raw1[raw1[,1]%in%new,]
            raw<-rbind(raw2,raw1)
            rm(raw1,raw2)
        } else {
            raw<-raw1
            rm(raw1)
        }
    } else if(file.exists(fname2)){
        raw2<-read.delim(fname2,as.is=TRUE)
        raw<-raw2
        rm(raw2)
    } else {
        next
    }
    raw[,1]<-substr(raw[,1],1,16)
    input<-raw;rm(raw,fname1,fname2)
    
    # Create a list, rawcnv and linkage
    rawcnv<-cnvProcess(input)
    
    # Save the file (linkagelev is useless)
    save(rawcnv,file=paste0("tums010/",tum,"/",tum,"-rawcnv.rda"))
    rm(rawcnv)
    
}
```
