## Supplementary Code for "Pan-Cancer and Single-Cell modelling of genomic alterations through gene expression": 000d_aracne.R

```
#### Aracne pipeline on the cluster
# scancel -u fgiorgi0


############ Calculating thresholds
cd /gpfs/work/IscrC_tumornet/pancancer/tums010
aracne=/galileo/home/userexternal/fgiorgi0/programs/Aracne.jar # Aracne-AP v1.4 (runs with Java 8)
tfs=/gpfs/work/IscrC_tumornet/shared/genelists/tfgenes_2018_08_06.dat
cotfs=/gpfs/work/IscrC_tumornet/shared/genelists/cotfgenes_2018_08_06.dat
sig=/gpfs/work/IscrC_tumornet/shared/genelists/signaling_2018_08_06.dat
module load jre

# Calculate threshold (cotfs)
for tumAcro in ACC BLCA BRCA CESC CHOL COAD DLBC ESCA GBM HNSC KICH KIRC KIRP LAML LGG LIHC LUAD LUSC MESO OV PAAD PCPG PRAD READ SARC SKCM STAD TGCT THCA THYM UCEC UCS UVM
do
qlogo=/gpfs/scratch/userexternal/fgiorgi0/qlogs/ar_thr_${tumAcro}_out.txt
qloge=/gpfs/scratch/userexternal/fgiorgi0/qlogs/ar_thr_${tumAcro}_err.txt
command="java -Xmx5500M -jar $aracne \
-e $tumAcro/${tumAcro}-expmat.dat \
-o $tumAcro/aracne-cotfs/ \
--tfs $cotfs \
--pvalue 0.00000001 --seed 1 \
--calculateThreshold"
sbatch --wrap "$command" -o $qlogo -e $qloge -J th_${tumAcro} -t 2:00:00 --mem=8000M -p gll_usr_prod -A iscrc_tumornet
done

# The same threshold is recycled for TF and signaling networks
cd /gpfs/work/IscrC_tumornet/pancancer/tums010
for tumAcro in ACC BLCA BRCA CESC CHOL COAD DLBC ESCA GBM HNSC KICH KIRC KIRP LAML LGG LIHC LUAD LUSC MESO OV PAAD PCPG PRAD READ SARC SKCM STAD TGCT THCA THYM UCEC UCS UVM
do
mkdir ${tumAcro}/aracne-tfs/
mkdir ${tumAcro}/aracne-sig/
cp ${tumAcro}/aracne-cotfs/miThreshold* ${tumAcro}/aracne-tfs/
cp ${tumAcro}/aracne-cotfs/miThreshold* ${tumAcro}/aracne-sig/
done


##### Run 100 bootstraps
cd /gpfs/work/IscrC_tumornet/pancancer/tums010
aracne=/galileo/home/userexternal/fgiorgi0/programs/Aracne.jar # Aracne-AP v1.4 (runs with Java 8)
tfs=/gpfs/work/IscrC_tumornet/shared/genelists/tfgenes_2018_08_06.dat
cotfs=/gpfs/work/IscrC_tumornet/shared/genelists/cotfgenes_2018_08_06.dat
sig=/gpfs/work/IscrC_tumornet/shared/genelists/signaling_2018_08_06.dat
module load jre


# cotfs
for tumAcro in ACC BLCA BRCA CESC CHOL COAD DLBC ESCA GBM HNSC KICH KIRC KIRP LAML LGG LIHC LUAD LUSC MESO OV PAAD PCPG PRAD READ SARC SKCM STAD TGCT THCA THYM UCEC UCS UVM
do
for i in {1..100}
do
qlogo=/gpfs/scratch/userexternal/fgiorgi0/qlogs/cotf_${tumAcro}_b${i}_out.txt
qloge=/gpfs/scratch/userexternal/fgiorgi0/qlogs/cotf_${tumAcro}_b${i}_err.txt
command="java -Xmx7000M -jar $aracne \
-e $tumAcro/${tumAcro}-expmat.dat \
-o $tumAcro/aracne-cotfs/ \
--tfs $cotfs \
--pvalue 0.00000001 \
--seed $i \
--threads 8"
sbatch --wrap "$command" -o $qlogo -e $qloge -J c_${tumAcro}_b$i -t 12:00:00 --mem=9000M -p gll_usr_prod -A iscrc_tumornet
done
done


# tfs
for tumAcro in ACC BLCA BRCA CESC CHOL COAD DLBC ESCA GBM HNSC KICH KIRC KIRP LAML LGG LIHC LUAD LUSC MESO OV PAAD PCPG PRAD READ SARC SKCM STAD TGCT THCA THYM UCEC UCS UVM
do
for i in {1..100}
do
qlogo=/gpfs/scratch/userexternal/fgiorgi0/qlogs/tf_${tumAcro}_b${i}_out.txt
qloge=/gpfs/scratch/userexternal/fgiorgi0/qlogs/tf_${tumAcro}_b${i}_err.txt
command="java -Xmx8000M -jar $aracne \
-e $tumAcro/${tumAcro}-expmat.dat \
-o $tumAcro/aracne-tfs/ \
--tfs $tfs \
--pvalue 0.00000001 \
--seed $i \
--threads 8"
sbatch --wrap "$command" -o $qlogo -e $qloge -J t_${tumAcro}_b$i -t 12:00:00 --mem=10000M -p gll_usr_prod -A iscrc_tumornet
done
done


# signaling
for tumAcro in ACC BLCA BRCA CESC CHOL COAD DLBC ESCA GBM HNSC KICH KIRC KIRP LAML LGG LIHC LUAD LUSC MESO OV PAAD PCPG PRAD READ SARC SKCM STAD TGCT THCA THYM UCEC UCS UVM
do
for i in {1..100}
do
qlogo=/gpfs/scratch/userexternal/fgiorgi0/qlogs/sig_${tumAcro}_b${i}_out.txt
qloge=/gpfs/scratch/userexternal/fgiorgi0/qlogs/sig_${tumAcro}_b${i}_err.txt
command="java -Xmx8000M -jar $aracne \
-e $tumAcro/${tumAcro}-expmat.dat \
-o $tumAcro/aracne-sig/ \
--tfs $sig \
--pvalue 0.00000001 \
--seed $i \
--threads 8"
sbatch --wrap "$command" -o $qlogo -e $qloge -J s_${tumAcro}_b$i -t 12:00:00 --mem=10000M -p gll_usr_prod -A iscrc_tumornet
done
done


### Check for completeness
cd /gpfs/work/IscrC_tumornet/pancancer/tums010
for tumAcro in ACC BLCA BRCA CESC CHOL COAD DLBC ESCA GBM HNSC KICH KIRC KIRP LAML LGG LIHC LUAD LUSC MESO OV PAAD PCPG PRAD READ SARC SKCM STAD TGCT THCA THYM UCEC UCS UVM
do
o1=`ls ${tumAcro}/aracne-cotfs/bootstrapNetwork* | wc -l` 
o2=`ls ${tumAcro}/aracne-tfs/bootstrapNetwork* | wc -l` 
o3=`ls ${tumAcro}/aracne-sig/bootstrapNetwork* | wc -l` 
echo "${tumAcro} COTF ${o1}    TF $o2   SIG $o3"
done


##### Consolidate
cd /gpfs/work/IscrC_tumornet/pancancer/tums010
aracne=/galileo/home/userexternal/fgiorgi0/programs/Aracne.jar # Aracne-AP v1.4 (runs with Java 8)
tfs=/gpfs/work/IscrC_tumornet/shared/genelists/tfgenes_2018_08_06.dat
cotfs=/gpfs/work/IscrC_tumornet/shared/genelists/cotfgenes_2018_08_06.dat
sig=/gpfs/work/IscrC_tumornet/shared/genelists/signaling_2018_08_06.dat
module load jre

for tumAcro in ACC BLCA BRCA CESC CHOL COAD DLBC ESCA GBM HNSC KICH KIRC KIRP LAML LGG LIHC LUAD LUSC MESO OV PAAD PCPG PRAD READ SARC SKCM STAD TGCT THCA THYM UCEC UCS UVM
do
qlogo=/gpfs/scratch/userexternal/fgiorgi0/qlogs/konsolidate_${tumAcro}_out.txt
qloge=/gpfs/scratch/userexternal/fgiorgi0/qlogs/konsolidate_${tumAcro}_err.txt
command="java -Xmx12000M -jar $aracne \
-o $tumAcro/aracne-cotfs/ \
--consolidate"
sbatch --wrap "$command" -o $qlogo -e $qloge -J kc_${tumAcro} -t 2:00:00 --mem=12000M -p gll_usr_prod -A iscrc_tumornet
command="java -Xmx12000M -jar $aracne \
-o $tumAcro/aracne-tfs/ \
--consolidate"
sbatch --wrap "$command" -o $qlogo -e $qloge -J kt_${tumAcro} -t 2:00:00 --mem=12000M -p gll_usr_prod -A iscrc_tumornet
command="java -Xmx12000M -jar $aracne \
-o $tumAcro/aracne-sig/ \
--consolidate"
sbatch --wrap "$command" -o $qlogo -e $qloge -J ks_${tumAcro} -t 2:00:00 --mem=12000M -p gll_usr_prod -A iscrc_tumornet
done


########## Convert into a regulon file (R)
### Convert from 4col to 3col format
cd /gpfs/work/IscrC_tumornet/pancancer/tums010/rawaracne
for tumAcro in ACC BLCA BRCA CESC CHOL COAD DLBC ESCA GBM HNSC KICH KIRC KIRP LAML LGG LIHC LUAD LUSC MESO OV PAAD PCPG PRAD READ SARC SKCM STAD TGCT THCA THYM UCEC UCS UVM
do
cut -f1-3 /gpfs/work/IscrC_tumornet/pancancer/tums010/${tumAcro}/aracne-tfs/network.txt > finalNetwork_tfs_${tumAcro}_3col.tsv
cut -f1-3 /gpfs/work/IscrC_tumornet/pancancer/tums010/${tumAcro}/aracne-cotfs/network.txt > finalNetwork_cotfs_${tumAcro}_3col.tsv
done

#scp -r:/gpfs/work/IscrC_tumornet/pancancer/tums010/rawaracne /mnt/d/Dropbox/projects/pancancer/tums010/

##### Local
library(viper) # Publicly available on Bioconductor
library(mixtools)
source("../shared/functions/aracne2regulon.R")
panFolder<-"D:/Dropbox/projects/pancancer/tums010"
setwd(panFolder)

if(TRUE){
    raw<-matrix(c(
        "ACC","Adrenocortical Carcinoma",
        "BLCA","Bladder Urothelial Carcinoma",
        "BRCA","Breast Invasive Carcinoma",
        "CESC","Cervical Squamous Cell carcinoma",
        "CHOL","Cholangiocarcinoma",
        "COAD","Colon Adenocarcinoma",
        "DLBC","Diffuse Large B-cell Lymphoma",
        "ESCA","Esophageal Carcinoma",
        "GBM","Glioblastoma Multiforme",
        "HNSC","Head and Neck Squamous Cell Carcinoma",
        "KICH","Kidney Chromophobe",
        "KIRC","Kidney Renal Clear Cell Carcinoma",
        "KIRP","Kidney Renal Papillary Cell Carcinoma",
        "LAML","Acute Myeloid Leukemia",
        "LGG","Brain Lower Grade Glioma",
        "LIHC","Liver Hepatocellular Carcinoma",
        "LUAD","Lung Adenocarcinoma",
        "LUSC","Lung Squamous Cell Carcinoma",
        "MESO","Mesothelioma",
        "OV","Ovarian Serous Cystadenocarcinoma",
        "PAAD","Pancreatic Adenocarcinoma",
        "PCPG","Pheochromocytoma and Paraganglioma",
        "PRAD","Prostate Adenocarcinoma",
        "READ","Rectum Adenocarcinoma",
        "SARC","Sarcoma",
        "SKCM","Skin Cutaneous Melanoma",
        "STAD","Stomach Adenocarcinoma",
        "TGCT","Testicular Germ Cell Tumors",
        "THCA","Thyroid Carcinoma",
        "THYM","Thymoma",
        "UCEC","Uterine Corpus Endometrial Carcinoma",
        "UCS","Uterine Carcinosarcoma",
        "UVM","Uveal Melanoma"
    ),ncol=2,byrow = TRUE
    )
    tums<-setNames(raw[,2],raw[,1])
    rm(raw)
} # Tumor Acronyms

tumAcros<-names(tums)
for(tumAcro in tumAcros){
    message("Doing ",tumAcro)
    nettypes<-c("tfs","cotfs")
    for(nettype in nettypes){
        input<-paste0("rawaracne/","finalNetwork_",nettype,"_",tumAcro,"_3col.tsv")
        load(paste0(tumAcro,"/",tumAcro,"-expmat.rda"))
        regul <- aracne2regulon(afile=input,eset=expmat)
        class(regul) <- "regulon"
        
        nettype<-sub("tfs","tf",nettype)
        output<-paste0(tumAcro,"/",tumAcro,"-",nettype,"-regulon.rda")
        save(regul,file=output)
    }
}
```
