## Supplementary Code for "Pan-Cancer and Single-Cell modelling of genomic alterations through gene expression": 000e_viper.R

```
rootFolder<-"D:/Dropbox/projects/pancancer/tums010/"
if(TRUE){
    raw<-matrix(c(
        "ACC","Adrenocortical Carcinoma",
        "BLCA","Bladder Urothelial Carcinoma",
        "BRCA","Breast Invasive Carcinoma",
        "CESC","Cervical Squamous Cell carcinoma",
        "CHOL","Cholangiocarcinoma",
        "COAD","Colon Adenocarcinoma",
        "DLBC","Diffuse Large B-cell Lymphoma",
        "ESCA","Esophageal Carcinoma",
        "GBM","Glioblastoma Multiforme",
        "HNSC","Head and Neck Squamous Cell Carcinoma",
        "KICH","Kidney Chromophobe",
        "KIRC","Kidney Renal Clear Cell Carcinoma",
        "KIRP","Kidney Renal Papillary Cell Carcinoma",
        "LAML","Acute Myeloid Leukemia",
        "LGG","Brain Lower Grade Glioma",
        "LIHC","Liver Hepatocellular Carcinoma",
        "LUAD","Lung Adenocarcinoma",
        "LUSC","Lung Squamous Cell Carcinoma",
        "MESO","Mesothelioma",
        "OV","Ovarian Serous Cystadenocarcinoma",
        "PAAD","Pancreatic Adenocarcinoma",
        "PCPG","Pheochromocytoma and Paraganglioma",
        "PRAD","Prostate Adenocarcinoma",
        "READ","Rectum Adenocarcinoma",
        "SARC","Sarcoma",
        "SKCM","Skin Cutaneous Melanoma",
        "STAD","Stomach Adenocarcinoma",
        "TGCT","Testicular Germ Cell Tumors",
        "THCA","Thyroid Carcinoma",
        "THYM","Thymoma",
        "UCEC","Uterine Corpus Endometrial Carcinoma",
        "UCS","Uterine Carcinosarcoma",
        "UVM","Uveal Melanoma"
    ),ncol=2,byrow = TRUE
    )
    tums<-setNames(raw[,2],raw[,1])
    rm(raw)
} # Tumor Acronyms

## Initialize
setwd(rootFolder)
library(viper)
library(dynamicTreeCut)


for(tumAcro in names(tums)){
    message("Doing ",tumAcro)
    
    ################## PANVIPER (vs normal and vs. mean)
    ### This will run 2 VIPER runs for each tumor type
    # Tumor vs. Tumor mean
    # Tumor vs. Normal

    
    ## Fix random seed for reproducibility
    set.seed(1)
    
    ### Run if all interactomes are ready
    if(
        !file.exists(paste0(tumAcro,"/",tumAcro,"-cotf-regulon.rda")) |
        !file.exists(paste0(tumAcro,"/",tumAcro,"-tf-regulon.rda"))
    ) next
    
    ### Networks
    load(paste0(tumAcro,"/",tumAcro,"-cotf-regulon.rda"))
    regul1<-regul
    load(paste0(tumAcro,"/",tumAcro,"-tf-regulon.rda"))
    regul2<-regul
    
    regulon<-c(regul1,regul2)
    rm(regul1,regul2,regul)
    
    ### Expression data
    load(paste0(tumAcro,"/",tumAcro,"-expmat.rda"))
    
    # Tumor/Metastatic/Normal expression
    # https://gdc.cancer.gov/resources-tcga-users/tcga-code-tables/sample-type-codes
    t<-colnames(expmat)[grep("01",substr(colnames(expmat),14,15))] # Primary Solid Tumor
    m<-colnames(expmat)[grep("06",substr(colnames(expmat),14,15))] # Metastasic
    n<-colnames(expmat)[grep("11",substr(colnames(expmat),14,15))] # Solid Tissue Normal
    l<-colnames(expmat)[grep("03",substr(colnames(expmat),14,15))] # Primary Blood Derived Cancer

    texpmat<-expmat[,t]
    mexpmat<-expmat[,m]
    nexpmat<-expmat[,n]
    lexpmat<-expmat[,l]
    if(tumAcro=="SKCM") texpmat<-cbind(texpmat,mexpmat)
    if(tumAcro=="LAML") texpmat<-lexpmat
    
    ### Tumor vs. Tumor viper (mean, without SD correction)
    signature <- viperSignature(texpmat, texpmat, method="mean")
    vipermat <- viper(signature, regulon, pleiotropy=FALSE, minsize=10)
    save(vipermat,file=paste0(tumAcro,"/",tumAcro,"-vipermat_tumor.rda"))
    rm(vipermat)
    
    ### Tumor vs. Normal viper
    if(is.matrix(nexpmat)&&ncol(nexpmat)>=5){
        signature <- viperSignature(texpmat, nexpmat)
        vipermat <- viper(signature, regulon, pleiotropy=FALSE, minsize=10)
        save(vipermat,file=paste0(tumAcro,"/",tumAcro,"-vipermat_normal.rda"))
        rm(vipermat)
    }
    
    
    
    
    
    
    
    
    
    
    ################## PANVIPER GROUPS (integration of signature vs. cluster groups)
    # Viper using clusters as reference
    
    ## Fix random seed for reproducibility
    set.seed(1)
    
    ### Networks
    load(paste0(tumAcro,"/",tumAcro,"-cotf-regulon.rda"))
    regulcotf<-regul
    load(paste0(tumAcro,"/",tumAcro,"-tf-regulon.rda"))
    regultf<-regul
    
    regulon<-c(regulcotf,regultf)
    rm(regul)
    
    ### Expression data
    load(paste0(tumAcro,"/",tumAcro,"-expmat.rda"))
    
    # Tumor/Metastatic/Normal expression
    # https://gdc.cancer.gov/resources-tcga-users/tcga-code-tables/sample-type-codes
    t<-colnames(expmat)[grep("01",substr(colnames(expmat),14,15))] # Primary Solid Tumor
    m<-colnames(expmat)[grep("06",substr(colnames(expmat),14,15))] # Metastasic
    n<-colnames(expmat)[grep("11",substr(colnames(expmat),14,15))] # Solid Tissue Normal
    l<-colnames(expmat)[grep("03",substr(colnames(expmat),14,15))] # Primary Blood Derived Cancer
    
    texpmat<-expmat[,t]
    mexpmat<-expmat[,m]
    nexpmat<-expmat[,n]
    lexpmat<-expmat[,l]
    if(tumAcro=="SKCM") texpmat<-cbind(texpmat,mexpmat)
    if(tumAcro=="LAML") texpmat<-lexpmat
    
    
    ### Defining the groups to use as reference, based on the TF-only viper matrix
    vp1 <- viper(texpmat-rowMeans(texpmat), regultf, pleiotropy=FALSE, method="none", verbose=FALSE, minsize=15)
    rvp1 <- apply(vp1, 2, function(x) sample(x))
    rownames(rvp1) <- rownames(vp1)
    svd1 <- svd(scale(t(vp1)), nv=0)
    pvar <- svd1$d^2/sum(svd1$d^2)
    pc1 <- diag(svd1$d) %*% t(svd1$u)
    colnames(pc1) <- colnames(texpmat)
    rsvd1 <- svd(scale(t(rvp1)), nv=0)
    rpvar <- rsvd1$d^2/sum(rsvd1$d^2)
    nopt <- which((pvar-rpvar)<0)[1]-1
    dd <- dist(t(pc1[1:nopt, ]))
    hc <- hclust(dd, "average")
    cuts <- cutreeDynamic(hc, minClusterSize=max(20, round(ncol(pc1)*.05)), distM=as.matrix(dd), deepSplit=TRUE)
    cuts[cuts==0] <- max(cuts)
    clus <- split(colnames(pc1), cuts)
    
    ### Running viper using the different clusters as reference
    # Also the cluster where the sample belongs is used as a reference, in order to prevent e.g. wrong cluster assignments
    vp1 <- mclapply(clus, function(ref, expset, regulon) {
        ss <- viperSignature(expset, expset[, colnames(expset) %in% ref], method="zscore", verbose=FALSE)
        viper(ss, regulon, method="none", pleiotropy=FALSE, minsize=10)
    }, expset=texpmat, regulon=regulon)
    tmp <- sapply(vp1, function(x) as.vector(x))
    vipermat <- matrix(rowSums(tmp^3)/rowSums(tmp^2), nrow(vp1[[1]]), ncol(vp1[[1]]), dimnames=list(rownames(vp1[[1]]), colnames(texpmat)))
    save(vipermat, file=paste(tumAcro, "/", tumAcro, "-vipermat_groups.rda", sep=""))
    
    
    
    
    
    
    
}
```
