## Supplementary Code for "Pan-Cancer and Single-Cell modelling of genomic alterations through gene expression": 000f_wgcna.R

```
setwd("D:/Dropbox/projects/pancancer")
library(WGCNA)
library(affy)

if(TRUE){
    raw<-matrix(c(
        "ACC","Adrenocortical Carcinoma",
        "BLCA","Bladder Urothelial Carcinoma",
        "BRCA","Breast Invasive Carcinoma",
        "CESC","Cervical Squamous Cell carcinoma",
        "CHOL","Cholangiocarcinoma",
        "COAD","Colon Adenocarcinoma",
        "DLBC","Diffuse Large B-cell Lymphoma",
        "ESCA","Esophageal Carcinoma",
        "GBM","Glioblastoma Multiforme",
        "HNSC","Head and Neck Squamous Cell Carcinoma",
        "KICH","Kidney Chromophobe",
        "KIRC","Kidney Renal Clear Cell Carcinoma",
        "KIRP","Kidney Renal Papillary Cell Carcinoma",
        "LAML","Acute Myeloid Leukemia",
        "LGG","Brain Lower Grade Glioma",
        "LIHC","Liver Hepatocellular Carcinoma",
        "LUAD","Lung Adenocarcinoma",
        "LUSC","Lung Squamous Cell Carcinoma",
        "MESO","Mesothelioma",
        "OV","Ovarian Serous Cystadenocarcinoma",
        "PAAD","Pancreatic Adenocarcinoma",
        "PCPG","Pheochromocytoma and Paraganglioma",
        "PRAD","Prostate Adenocarcinoma",
        "READ","Rectum Adenocarcinoma",
        "SARC","Sarcoma",
        "SKCM","Skin Cutaneous Melanoma",
        "STAD","Stomach Adenocarcinoma",
        "TGCT","Testicular Germ Cell Tumors",
        "THCA","Thyroid Carcinoma",
        "THYM","Thymoma",
        "UCEC","Uterine Corpus Endometrial Carcinoma",
        "UCS","Uterine Carcinosarcoma",
        "UVM","Uveal Melanoma"
    ),ncol=2,byrow = TRUE
    )
    tums<-setNames(raw[,2],raw[,1])
    rm(raw)
} # Tumor Acronyms


### Generate clusters with WGCNA
if(TRUE){
    for(tum in names(tums)[1:length(tums)]){
        load(paste0("tums010/",tum,"/",tum,"-expmat.rda"))
        datExpr<-t(expmat)
        rm(expmat)
        
        ### Select power threshold
        if(TRUE){
            png(paste0("plots/000f_wgcna/power-",tum,".png"),w=1000,h=800,p=30)
            # Choose a set of soft-thresholding powers
            powers = c(c(1:10), seq(from = 12, to=20, by=2))
            # Call the network topology analysis function
            sft = pickSoftThreshold(datExpr, powerVector = powers, verbose = 5)
            # Plot the results:
            par(mfrow = c(1,2))
            # Scale-free topology fit index as a function of the soft-thresholding power
            plot(sft$fitIndices[,1], -sign(sft$fitIndices[,3])*sft$fitIndices[,2],
                 xlab="Soft Threshold (power)",ylab="Scale Free Topology Model Fit,signed R^2",type="n",
                 main = paste("Scale independence"));
            text(sft$fitIndices[,1], -sign(sft$fitIndices[,3])*sft$fitIndices[,2],
                 labels=powers,cex=0.9,col="red");
            # this line corresponds to using an R^2 cut-off of h
            abline(h=0.90,col="red")
            # Mean connectivity as a function of the soft-thresholding power
            plot(sft$fitIndices[,1], sft$fitIndices[,5],
                 xlab="Soft Threshold (power)",ylab="Mean Connectivity", type="n",
                 main = paste("Mean connectivity"))
            text(sft$fitIndices[,1], sft$fitIndices[,5], labels=powers, cex=0.9,col="red")
            dev.off()
            
            # The result is shown in Fig. 1. We choose the power 6, which is the lowest power for which the scale-free topology fit
            # index curve flattens out upon reaching a high value (in this case, roughly 0.90).
            topology<-(-sign(sft$fitIndices[,3])*sft$fitIndices[,2])
            subtopology<-which(topology>=0.90)
            threshold<-subtopology[1]
            if(is.na(threshold)){
                subtopology<-which(topology>=0.80)
                threshold<-subtopology[1]
            }
            if(is.na(threshold)){
                subtopology<-which(topology>=0.70)
                threshold<-subtopology[1]
            }
            message(tum," has soft threshold power of ", threshold)
        }
        
        ### One-step network construction and module detection
        ## Minimum module size should be the same as VIPER
        net<-blockwiseModules(datExpr,power=threshold,minModuleSize=10,
                              numericLabels=TRUE,pamRespectsDendro=FALSE,
                              verbose=3,maxBlockSize=10000)
        
        ## Plot clusters
        png(paste0("plots/000f_wgcna/dendro-",tum,".png"),w=1000,h=800,p=30)
        mergedColors<-labels2colors(net$colors)
        plotDendroAndColors(net$dendrograms[[1]], mergedColors[net$blockGenes[[1]]],
                            "Module colors",
                            dendroLabels = FALSE, hang = 0.03,
                            addGuide = TRUE, guideHang = 0.05)
        dev.off()    
        
        
        ## Save clusters
        wgcna<-setNames(net$colors,colnames(datExpr))
        message(tum," has this many clusters: ",length(unique(wgcna)))
        save(wgcna,file=paste0("tums010/",tum,"/",tum,"-wgcna.rda"))
    }
}


## To obtain activity, we do a robust mean of the genes in the cluster
## 1) median polish (I have a paper on it)
## 2) tukey biweight (cute)
## 3) eigengenes (in the default wgcna pipeline)


### Generate wgcnamat
for(tum in names(tums)){
    ### Take vipermat (tumor samples)
    load(paste0("tums010/",tum,"/",tum,"-vipermat_tumor.rda"))
    
    ### Take expmat
    load(paste0("tums010/",tum,"/",tum,"-expmat.rda"))
    expmat<-expmat[,colnames(vipermat)]
    
    message("Doing ",tum)
    fname<-paste0("tums010/",tum,"/",tum,"-wgcnamat.rda")
    # unlink(fname)}
    if(!file.exists(fname)){
        ## Create wgcnamat by tukey biweight
        load(paste0("tums010/",tum,"/",tum,"-wgcna.rda"))
        wgcna<-sort(setNames(paste0("w",wgcna),names(wgcna)))
        wgcnamat<-matrix(NA,nrow=length(unique(wgcna)),ncol=ncol(vipermat))
        colnames(wgcnamat)<-colnames(vipermat)
        rownames(wgcnamat)<-unique(wgcna)
        pb<-txtProgressBar(0,nrow(wgcnamat),style=3)
        il<-0
        for(i in unique(wgcna)){
            wgenes<-names(which(wgcna==i))
            #wpolish<-medpolish(t(wsubmat))
            if(length(wgenes)==1){
                wtukey<-expmat[wgenes,]
            } else {
                wsubmat<-expmat[wgenes,]
                wtukey<-apply(wsubmat,2,tukey.biweight)
            }
            wgcnamat[i,]<-wtukey
            il<-il+1
            setTxtProgressBar(pb,il)
        }
        save(wgcnamat,file=fname)
    } else {
        load(fname)
    }
}
```
