## Supplementary Code for "Pan-Cancer and Single-Cell modelling of genomic alterations through gene expression": 000h_ccle.R

```
# Downloaded on 20-Feb-2019 from https://portals.broadinstitute.org/ccle/data
# File version is 02-Jan-2019
raw<-read.delim("../shared/ccle/raw/CCLE_RNAseq_genes_counts_20180929.gct.gz",skip=2)
dim(raw) # 56202 1021
rawcounts<-raw[,3:ncol(raw)]
rownames(rawcounts)<-raw[,1]
raw<-rawcounts
rm(rawcounts)

# Format and normalize
raw<-as.matrix(raw)
source("../shared/functions/geneids.R")
fname<-"../shared/ccle/ccle-rawcounts.rda"
if(!file.exists(fname)){
    ensids<-gsub("\\.[0-9]+","",rownames(raw))
    rownames(raw)<-ensids
    egs<-setNames(eg2sym(ens2eg(ensids)),ensids)
    egs<-egs[!is.na(egs)]
    rawcounts<-matrix(0,nrow=length(unique(egs)),ncol=ncol(raw))
    rownames(rawcounts)<-unique(egs)
    colnames(rawcounts)<-colnames(raw)
    
    # Sum counts of ENSG models mapping the same Entrez Gene
    for(i in 1:nrow(rawcounts)){
        eghere<-rownames(rawcounts)[i]
        enshere<-names(which(egs==eghere))
        message(i,"/",nrow(rawcounts),": ",length(enshere))
        if(length(enshere)==1){
            rawcounts[eghere,]<-raw[enshere,]
        } else {
            mysum<-apply(raw[enshere,],2,sum,na.rm=TRUE)
            rawcounts[eghere,]<-mysum
        }
    }
    save(rawcounts,file=fname)
} else {load(fname)}

# VST-normalize
#biocLite(c("DESeq","Biobase"))
library(DESeq)
library(Biobase)
fname<-"../shared/ccle/ccle-expmat.rda"
if(!file.exists(fname)){
    conditions <- c()
    samples<-colnames(rawcounts)
    conditions <- factor(samples)
    cds <- DESeq::newCountDataSet(rawcounts, conditions)
    cds <- DESeq::estimateSizeFactors(cds)
    cds <- DESeq::estimateDispersions(cds, method="blind")
    vsd <- DESeq::varianceStabilizingTransformation(cds)
    expmat <- Biobase::exprs(vsd)
    rownames(expmat) <- rownames(rawcounts)
    save(expmat,file=fname)
    
} else {load(fname)}


# File for ARACNe
cat("gene\t",file="../shared/ccle/ccle-expmat.dat")
write.table(expmat,
            file="../shared/ccle/ccle-expmat.dat",
            col.names=TRUE,
            sep="\t",
            quote=FALSE,
            append=TRUE
)
```
