## Supplementary Code for "Pan-Cancer and Single-Cell modelling of genomic alterations through gene expression": 000i_ccleAracne.sh

```
scp /mnt/d/Dropbox/projects/shared/ccle/ccle-expmat.dat \
:/gpfs/work/IscrC_tumornet/ccle/


ssh


# Run ARACNe

############ Calculating thresholds
cd /gpfs/work/IscrC_tumornet/ccle
aracne=/galileo/home/userexternal/fgiorgi0/programs/Aracne.jar # Aracne-AP v1.4 (runs with Java 8)
tfs=/gpfs/work/IscrC_tumornet/shared/genelists/tfgenes_2018_08_06.symbol.dat
cotfs=/gpfs/work/IscrC_tumornet/shared/genelists/cotfgenes_2018_08_06.symbol.dat
sig=/gpfs/work/IscrC_tumornet/shared/genelists/signaling_2018_08_06.symbol.dat
module load jre

# Calculate threshold (cotfs)
cd /gpfs/work/IscrC_tumornet/ccle
mkdir /gpfs/scratch/userexternal/fgiorgi0/qlogs
qlogo=/gpfs/scratch/userexternal/fgiorgi0/qlogs/thr_out.txt
qloge=/gpfs/scratch/userexternal/fgiorgi0/qlogs/thr_err.txt
command="java -Xmx10000M -jar $aracne \
-e ccle-expmat.dat \
-o aracne-cotfs/ \
--tfs $cotfs \
--pvalue 0.00000001 --seed 1 \
--calculateThreshold"
sbatch --wrap "$command" -o $qlogo -e $qloge -J th -t 2:00:00 --mem=16000M -p gll_usr_prod -A iscrc_tumornet

# The same threshold is recycled for TF and signaling networks
cd /gpfs/work/IscrC_tumornet/ccle
mkdir aracne-tfs/
mkdir aracne-sig/
cp aracne-cotfs/miThreshold* aracne-tfs/
cp aracne-cotfs/miThreshold* aracne-sig/


##### Run 100 bootstraps
cd /gpfs/work/IscrC_tumornet/ccle
aracne=/galileo/home/userexternal/fgiorgi0/programs/Aracne.jar # Aracne-AP v1.4 (runs with Java 8)
tfs=/gpfs/work/IscrC_tumornet/shared/genelists/tfgenes_2018_08_06.symbol.dat
cotfs=/gpfs/work/IscrC_tumornet/shared/genelists/cotfgenes_2018_08_06.symbol.dat
sig=/gpfs/work/IscrC_tumornet/shared/genelists/signaling_2018_08_06.symbol.dat
module load jre

# cotfs
for i in {1..100}
do
qlogo=/gpfs/scratch/userexternal/fgiorgi0/qlogs/cotf_out_b{i}.txt
qloge=/gpfs/scratch/userexternal/fgiorgi0/qlogs/cotf_err_b{i}.txt
command="java -Xmx7000M -jar $aracne \
-e ccle-expmat.dat \
-o aracne-cotfs/ \
--tfs $cotfs \
--pvalue 0.00000001 \
--seed $i \
--threads 8"
sbatch --wrap "$command" -o $qlogo -e $qloge -J c_$i -t 12:00:00 --mem=9000M -p gll_usr_prod -A iscrc_tumornet --ntasks-per-node=8
done


# tfs
for i in {1..100}
do
qlogo=/gpfs/scratch/userexternal/fgiorgi0/qlogs/cotf_out_b{i}.txt
qloge=/gpfs/scratch/userexternal/fgiorgi0/qlogs/cotf_err_b{i}.txt
command="java -Xmx7000M -jar $aracne \
-e ccle-expmat.dat \
-o aracne-tfs/ \
--tfs $tfs \
--pvalue 0.00000001 \
--seed $i \
--threads 8"
sbatch --wrap "$command" -o $qlogo -e $qloge -J t_$i -t 12:00:00 --mem=9000M -p gll_usr_prod -A iscrc_tumornet --ntasks-per-node=8
done


# signaling
for i in {1..100}
do
qlogo=/gpfs/scratch/userexternal/fgiorgi0/qlogs/sig_out_b{i}.txt
qloge=/gpfs/scratch/userexternal/fgiorgi0/qlogs/sig_err_b{i}.txt
command="java -Xmx12000M -jar $aracne \
-e ccle-expmat.dat \
-o aracne-sig/ \
--tfs $sig \
--pvalue 0.00000001 \
--seed $i \
--threads 8"
sbatch --wrap "$command" -o $qlogo -e $qloge -J s_$i -t 12:00:00 --mem=16000M -p gll_usr_prod -A iscrc_tumornet --ntasks-per-node=8
done


### Check for completeness
cd /gpfs/work/IscrC_tumornet/ccle
o1=`ls aracne-cotfs/bootstrapNetwork* | wc -l` 
o2=`ls aracne-tfs/bootstrapNetwork* | wc -l` 
o3=`ls aracne-sig/bootstrapNetwork* | wc -l` 
echo "${tumAcro} COTF ${o1}    TF $o2   SIG $o3"


##### Consolidate
cd /gpfs/work/IscrC_tumornet/ccle
aracne=/galileo/home/userexternal/fgiorgi0/programs/Aracne.jar # Aracne-AP v1.4 (runs with Java 8)
module load jre

qlogo=/gpfs/scratch/userexternal/fgiorgi0/qlogs/konsolidate_out.txt
qloge=/gpfs/scratch/userexternal/fgiorgi0/qlogs/konsolidate_err.txt
command="java -Xmx10000M -jar $aracne -o aracne-cotfs/ --consolidate"
sbatch --wrap "$command" -o $qlogo -e $qloge -J kc -t 2:00:00 --mem=12000M -p gll_usr_prod -A iscrc_tumornet
command="java -Xmx10000M -jar $aracne -o aracne-tfs/ --consolidate"
sbatch --wrap "$command" -o $qlogo -e $qloge -J kt -t 2:00:00 --mem=12000M -p gll_usr_prod -A iscrc_tumornet
command="java -Xmx10000M -jar $aracne -o aracne-sig/ --consolidate"
sbatch --wrap "$command" -o $qlogo -e $qloge -J ks -t 2:00:00 --mem=12000M -p gll_usr_prod -A iscrc_tumornet


########## Convert into a regulon file (R)
### Convert from 4col to 3col format
cd /gpfs/work/IscrC_tumornet/ccle
cut -f1-3 aracne-tfs/network.txt > finalNetwork_tfs_ccle_3col.tsv
cut -f1-3 aracne-cotfs/network.txt > finalNetwork_cotfs_ccle_3col.tsv

#scp -r:/gpfs/work/IscrC_tumornet/ccle/finalNetwork* /mnt/d/Dropbox/projects/shared/ccle/

##### Local
library(viper) # Publicly available on Bioconductor
library(mixtools)
source("../shared/functions/aracne2regulon.R")

nettypes<-c("tfs","cotfs")
for(nettype in nettypes){
    input<-paste0("../shared/ccle/finalNetwork_",nettype,"_ccle_3col.tsv")
    load("../shared/ccle/ccle-expmat.rda")
    regul <- aracne2regulon(afile=input,eset=expmat)
    class(regul) <- "regulon"
    nettype<-sub("tfs","tf",nettype)
    output<-paste0("../shared/ccle/ccle-",nettype,"-regulon.rda")
    save(regul,file=output)
}
```
