## Supplementary Code for "Pan-Cancer and Single-Cell modelling of genomic alterations through gene expression": 000j_aracnetxt.R

```
load("tums.rda")
source("../shared/functions/geneids.R")

for(tum in names(tums)){
    if(tum=="NBL"){next}
    message("Doing ",tum)
    load(paste0("tums010/",tum,"/",tum,"-tf-regulon.rda"))
    fname<-paste0("aracnetxt/",tum,"-tf-regulon.txt")
    cat("Regulator\tTarget\tMoA\tlikelihood\tRegulator_Entrez\tTarget_Entrez\n",file=fname)
    for(tf in names(regul)){
        sub<-regul[[tf]]
        toprint<-cbind(
            rep(eg2sym(tf),length(sub$tfmode)),
            eg2sym(names(sub$tfmode)),
            sub$tfmode,
            sub$likelihood,
            rep(tf,length(sub$tfmode)),
            names(sub$tfmode)
        )
        write.table(toprint,file=fname,append=TRUE,col=FALSE,row=FALSE,quote=FALSE,sep="\t")
    }
    
    load(paste0("tums010/",tum,"/",tum,"-cotf-regulon.rda"))
    fname<-paste0("aracnetxt/",tum,"-cotf-regulon.txt")
    cat("Regulator\tTarget\tMoA\tlikelihood\tRegulator_Entrez\tTarget_Entrez\n",file=fname)
    for(tf in names(regul)){
        sub<-regul[[tf]]
        toprint<-cbind(
            rep(eg2sym(tf),length(sub$tfmode)),
            eg2sym(names(sub$tfmode)),
            sub$tfmode,
            sub$likelihood,
            rep(tf,length(sub$tfmode)),
            names(sub$tfmode)
        )
        write.table(toprint,file=fname,append=TRUE,col=FALSE,row=FALSE,quote=FALSE,sep="\t")
    }
}
```
