## Supplementary Code for "Pan-Cancer and Single-Cell modelling of genomic alterations through gene expression": 001_fCNVs.R

```
### Basically, just load CNVs and expression, and keep those that cis-correlate >= 0.5

if(TRUE){
    raw<-matrix(c(
        "ACC","Adrenocortical Carcinoma",
        "BLCA","Bladder Urothelial Carcinoma",
        "BRCA","Breast Invasive Carcinoma",
        "CESC","Cervical Squamous Cell carcinoma",
        "CHOL","Cholangiocarcinoma",
        "COAD","Colon Adenocarcinoma",
        "DLBC","Diffuse Large B-cell Lymphoma",
        "ESCA","Esophageal Carcinoma",
        "GBM","Glioblastoma Multiforme",
        "HNSC","Head and Neck Squamous Cell Carcinoma",
        "KICH","Kidney Chromophobe",
        "KIRC","Kidney Renal Clear Cell Carcinoma",
        "KIRP","Kidney Renal Papillary Cell Carcinoma",
        "LAML","Acute Myeloid Leukemia",
        "LGG","Brain Lower Grade Glioma",
        "LIHC","Liver Hepatocellular Carcinoma",
        "LUAD","Lung Adenocarcinoma",
        "LUSC","Lung Squamous Cell Carcinoma",
        "MESO","Mesothelioma",
        "OV","Ovarian Serous Cystadenocarcinoma",
        "PAAD","Pancreatic Adenocarcinoma",
        "PCPG","Pheochromocytoma and Paraganglioma",
        "PRAD","Prostate Adenocarcinoma",
        "READ","Rectum Adenocarcinoma",
        "SARC","Sarcoma",
        "SKCM","Skin Cutaneous Melanoma",
        "STAD","Stomach Adenocarcinoma",
        "TGCT","Testicular Germ Cell Tumors",
        "THCA","Thyroid Carcinoma",
        "THYM","Thymoma",
        "UCEC","Uterine Corpus Endometrial Carcinoma",
        "UCS","Uterine Carcinosarcoma",
        "UVM","Uveal Melanoma"
    ),ncol=2,byrow = TRUE
    )
    tums<-setNames(raw[,2],raw[,1])
    rm(raw)
}


plotmat<-matrix(0,nrow=2,ncol=length(tums))
colnames(plotmat)<-names(tums)
rownames(plotmat)<-c("cnv","fcnv")

for(tum in names(tums)){
    message("Doing ",tums[tum])
    load(paste0("tums010/",tum,"/",tum,"-rawcnv.rda"))
    load(paste0("tums010/",tum,"/",tum,"-expmat.rda"))
    colnames(rawcnv)<-substr(colnames(rawcnv),1,15)
    colnames(expmat)<-substr(colnames(expmat),1,15)
    common<-intersect(colnames(expmat),colnames(rawcnv))
    rawcnv<-rawcnv[,common]
    expmat<-expmat[,common]
    common<-intersect(rownames(expmat),rownames(rawcnv))
    rawcnv<-rawcnv[common,]
    expmat<-expmat[common,]

    cors<-rep(0,nrow(rawcnv))
    for(i in 1:nrow(expmat)){
        cors[i]<-cor(rawcnv[i,],expmat[i,],method="spearman")
    }
    names(cors)<-rownames(expmat)
    functional<-names(cors[abs(cors)>=0.5])
    functional<-functional[!is.na(functional)]
    fcnv<-rawcnv[functional,]
    if(length(functional)==1){
        fcnv<-t(as.matrix(fcnv))
        rownames(fcnv)<-functional
    }
    save(fcnv,file=paste0("tums010/",tum,"/",tum,"-fcnv.rda"))
    
    plotmat["cnv",tum]<-nrow(rawcnv)
    plotmat["fcnv",tum]<-nrow(fcnv)
}


if(TRUE){
    # https://stackoverflow.com/questions/15282580/how-to-generate-a-number-of-most-distinctive-colors-in-r
    library(RColorBrewer)
    brewer.pal.info
    n<-length(tums)
    qual_col_pals<-brewer.pal.info[brewer.pal.info$category == 'qual',]
    cols<-unlist(mapply(brewer.pal, qual_col_pals$maxcolors, rownames(qual_col_pals)))
    set.seed(3)
    tumcols<-setNames(sample(cols, n),names(tums))
    #pie(rep(1,n),col=tumcols)
} # Tumor Colors

png("plots/001_fCNV_barplot.png",w=1200,h=1000,p=30)
par(las=2)
x<-sort(plotmat["fcnv",],dec=TRUE)
barplot(x,col=tumcols[names(x)],ylab="Genes Targeted by fCNV",main="Functional CNVs in TCGA")
dev.off()
```
