## Supplementary Code for "Pan-Cancer and Single-Cell modelling of genomic alterations through gene expression": 010_PaperStructure.R

```
### TCGA description (3 figures)
# Cancer sizes (nice cloud TSNE)
# Most mutated genes
# Most (functional) CNVs-affected genes (amp+del)


### Proof of concept
# Expression can predict TP53 mutations in breast cancer (GBM model)
# Test all mutations in breast cancer (CDH1 another good example)
# Most affected genes by TP53

### Pan-cancer mutation/expression relationships
# Test all somatic mutations in each cancer, test which ones are most predictive
# Show how some classes of genes (e.g. cancer drivers) are more predictive than others (within similar ranges of mutated samples number)
# Show differences in C/M classes of tumors


### Test different classifiers with caret
methods<-c("bayesglm","gbm","glmnet","lda","nnet","rf","svmLinear","svmRadial")
# Bayesian Generalized Linear Model: bayesglm
# Gradient Boost Modeling: gbm
# Generalized Linear Model: glmnet
# Linear Discriminant Analysis: lda
# Neural Network: nnet
# Random Forest: rf
# Support Vector Machine: svmLinear
# Support Vector Machine: svmRadial


### Model robustness
# To gaussian noise
# To read decrease
# Test Expression vs. WGCNA vs. VIPER


### Single Cell
# Circulating prostate cells dataset (Miyamoto et al.,), Patient CTCs and Primary Tumor Cells
# Predict the presence of TP53 mutations vs Test them via RNASeq?
# CCLE vs. itself (proof: it works on patient and non-patient data)
# Cross-dataset prediction
```
