## Supplementary Code for "Pan-Cancer and Single-Cell modelling of genomic alterations through gene expression": 011_testGenes.R

```
setwd("D:/Dropbox/projects/pancancer/")


library(caret)
library(pROC)
if(TRUE){
    raw<-matrix(c(
        "ACC","Adrenocortical Carcinoma",
        "BLCA","Bladder Urothelial Carcinoma",
        "BRCA","Breast Invasive Carcinoma",
        "CESC","Cervical Squamous Cell carcinoma",
        "CHOL","Cholangiocarcinoma",
        "COAD","Colon Adenocarcinoma",
        "DLBC","Diffuse Large B-cell Lymphoma",
        "ESCA","Esophageal Carcinoma",
        "GBM","Glioblastoma Multiforme",
        "HNSC","Head and Neck Squamous Cell Carcinoma",
        "KICH","Kidney Chromophobe",
        "KIRC","Kidney Renal Clear Cell Carcinoma",
        "KIRP","Kidney Renal Papillary Cell Carcinoma",
        "LAML","Acute Myeloid Leukemia",
        "LGG","Brain Lower Grade Glioma",
        "LIHC","Liver Hepatocellular Carcinoma",
        "LUAD","Lung Adenocarcinoma",
        "LUSC","Lung Squamous Cell Carcinoma",
        "MESO","Mesothelioma",
        "OV","Ovarian Serous Cystadenocarcinoma",
        "PAAD","Pancreatic Adenocarcinoma",
        "PCPG","Pheochromocytoma and Paraganglioma",
        "PRAD","Prostate Adenocarcinoma",
        "READ","Rectum Adenocarcinoma",
        "SARC","Sarcoma",
        "SKCM","Skin Cutaneous Melanoma",
        "STAD","Stomach Adenocarcinoma",
        "TGCT","Testicular Germ Cell Tumors",
        "THCA","Thyroid Carcinoma",
        "THYM","Thymoma",
        "UCEC","Uterine Corpus Endometrial Carcinoma",
        "UCS","Uterine Carcinosarcoma",
        "UVM","Uveal Melanoma"
    ),ncol=2,byrow = TRUE
    )
    tums<-setNames(raw[,2],raw[,1])
    rm(raw)
} # Tumor Acronyms
source("../shared/functions/geneids.R")
if(TRUE){
    # https://stackoverflow.com/questions/15282580/how-to-generate-a-number-of-most-distinctive-colors-in-r
    library(RColorBrewer)
    brewer.pal.info
    n<-length(tums)
    qual_col_pals<-brewer.pal.info[brewer.pal.info$category == 'qual',]
    cols<-unlist(mapply(brewer.pal, qual_col_pals$maxcolors, rownames(qual_col_pals)))
    set.seed(1)
    tumcols<-setNames(sample(cols, n),names(tums))
    #pie(rep(1,n),col=tumcols)
} # Tumor Colors


### Occurrence in TCGA of examples
# Coding mutations of TP53
if(TRUE){
    for(gene in any2entrez(c("NFKB1","TP53"))){
        res<-matrix(0,ncol=length(tums),nrow=2);dimnames(res)<-list(c("n","perc"),names(tums))
        for(tum in names(tums)){
            load(paste0("tums010/",tum,"/",tum,"-mutations.rda"))
            if(gene%in%rownames(mutations)){
                track<-mutations[gene,]
                res["n",tum]<-sum(track)
                res["perc",tum]<-100*sum(track)/length(track)
            }
        }
        png(paste0("plots/011a_mutations",eg2sym(gene),".png"),w=1400,h=1000,p=35)
        par(las=2)
        res<-res[,order(-res["perc",])]
        y<-res["perc",]
        bp<-barplot(y,col=tumcols[names(y)],main=paste0(eg2sym(gene)," Somatic Mutations"),ylim=c(0,max(y)*1.1))
        par(las=3)
        grid(col="darkgray")
        mtext("% Samples",2,line=2)
        text(bp,y,res["n",],pos=3,offset=0.3,cex=0.5,font=2)
        dev.off()
    }
}

# Amplifications
if(TRUE){
    for(gene in any2entrez(c("PVT1","EGFR"))){
        res<-matrix(0,ncol=length(tums),nrow=2);dimnames(res)<-list(c("n","perc"),names(tums))
        for(tum in names(tums)){
            load(paste0("tums010/",tum,"/",tum,"-fcnv.rda"))
            if(gene%in%rownames(fcnv)){
                track<-fcnv[gene,]
                track<-(track>=0.5)
                res["n",tum]<-sum(track)
                res["perc",tum]<-100*sum(track)/length(track)
            }
        }
        png(paste0("plots/011b_amplifications",eg2sym(gene),".png"),w=1400,h=1000,p=35)
        par(las=2)
        res<-res[,order(-res["perc",])]
        y<-res["perc",]
        bp<-barplot(y,col=tumcols[names(y)],main=paste0(eg2sym(gene)," Functional Amplifications"),ylim=c(0,max(y)*1.1))
        par(las=3)
        grid(col="darkgray")
        mtext("% Samples",2,line=2)
        text(bp,y,res["n",],pos=3,offset=0.3,cex=0.5,font=2)
        dev.off()
    }
}


# DELETIONS
if(TRUE){
    for(gene in any2entrez(c("PTEN","NFKB1"))){
        res<-matrix(0,ncol=length(tums),nrow=2);dimnames(res)<-list(c("n","perc"),names(tums))
        for(tum in names(tums)){
            load(paste0("tums010/",tum,"/",tum,"-fcnv.rda"))
            if(gene%in%rownames(fcnv)){
                track<-fcnv[gene,]
                track<-(track<=(-0.5))
                res["n",tum]<-sum(track)
                res["perc",tum]<-100*sum(track)/length(track)
            }
        }
        png(paste0("plots/011b_deletions",eg2sym(gene),".png"),w=1400,h=1000,p=35)
        par(las=2)
        res<-res[,order(-res["perc",])]
        y<-res["perc",]
        bp<-barplot(y,col=tumcols[names(y)],main=paste0(eg2sym(gene)," Functional Deletions"),ylim=c(0,max(y)*1.1))
        par(las=3)
        grid(col="darkgray")
        mtext("% Samples",2,line=2)
        text(bp,y,res["n",],pos=3,offset=0.3,cex=0.5,font=2)
        dev.off()
    }
}


### Binary Classifiers
## Specify tumor type and gene entrez id 
tum<-"BRCA"
gene<-"7157"

## Methods to be used
methods<-c("bayesglm","gbm","glmnet","lda","nnet","rf","svmLinear","svmRadial")
# Bayesian Generalized Linear Model: bayesglm
# Gradient Boost Modeling: gbm
# Generalized Linear Model: glmnet
# Linear Discriminant Analysis: lda
# Neural Network: nnet
# Random Forest: rf
# Support Vector Machine: svmLinear
# Support Vector Machine: svmRadial
method<-"glmnet"

# Loads the two data files from the tumor type inputted to the function
load(paste0("tums010/",tum,"/",tum,"-mutations.rda"))
load(paste0("tums010/",tum,"/",tum,"-expmat.rda"))
colnames(mutations)<-substr(colnames(mutations),1,15)
colnames(expmat)<-substr(colnames(expmat),1,15)

# Creates a data frame with variables (genes) and observations (samples) that includes the mutations data for the specific mutation
int.samples<-intersect(colnames(expmat),colnames(mutations))
mutations.a<-mutations[,int.samples]
gene.mut<-as.vector(mutations.a[as.character(gene),])
gene.mut<-ifelse(gene.mut==1,"mut","wt")
expmat.a<-expmat[,int.samples]
# Filter by Variance (Top 1000)
evars<-apply(expmat.a,1,var)
expmat.a<-expmat.a[evars>0,]
expmat.a<-expmat.a[names(sort(evars,dec=TRUE)[1:1000]),]
expmat.t<-cbind(t(expmat.a),gene.mut)
df<-as.data.frame(expmat.t)
df$gene.mut<-as.factor(df$gene.mut)
percmut<-signif(100*sum(as.vector(mutations.a[as.character(gene),]))/nrow(df),4)

# Many caret functions model the probability of the first factor level
df$gene.mut<-relevel(df$gene.mut,ref="mut")

# TODO avoid test if samples are too few or if percmut is not within optimal range (5-95%)
message(tum,": testing on ",nrow(df)," samples with a % of mutations for ",eg2sym(gene)," of ",percmut)

# We want to predict the outcome (the mutations), from a set of predictors (gene expression data), which we code here
prop.outcome<-prop.table(table(df[,"gene.mut"]))
outcomeName<-"gene.mut"
predictorsNames<-colnames(df)[colnames(df)!=outcomeName]

# Set a seed to use, to make the CV steps reproducible
set.seed(1234)

# Split the data into training and test sets (0.6, 0.8, try different ones?)
splitIndex<-createDataPartition(df[,outcomeName],p=0.75,list=FALSE,times=1)
trainDF<-df[splitIndex,]
testDF<-df[-splitIndex,]

# Train Model Parameters (10 is acceptable)
objControl<-trainControl(method="repeatedcv",number=10,returnResamp='none',summaryFunction=twoClassSummary,classProbs=TRUE)
objModel<-train(data.matrix(trainDF[,predictorsNames]),trainDF[,outcomeName], 
                method=method, 
                trControl=objControl,  
                metric="ROC"
)


# Validating the Model
# It will predict using the best model: objModel$finalModel
predictions<-predict(object=objModel,data.matrix(testDF[,predictorsNames]))
probabilities<-predict(object=objModel,data.matrix(testDF[,predictorsNames]),type="prob")
rocCurve<-roc(response=testDF$gene.mut,predictor=probabilities[,"mut"])

# Confusion matrix
confusionMatrix(predictions,testDF$gene.mut)

# Plot the ROC Curve with AUC
png("plots/011c_exampleROC_TP53.png",w=1400,h=1000,p=35)
plot(rocCurve,col="red3",cex.lab=2,cex.main=1.1,print.auc=FALSE,lwd=2,
     main=paste0(method," Model predicting ",eg2sym(gene)," Mutations in ",tum," dataset"))
mtext("10-fold CV",side=1)
grid(col="lightgrey")
legend(0.2,0.2,legend=paste0("AUC= ", round(rocCurve$auc,4)))
dev.off()

# Extract predictors from objModel$finalModel
if(method=="glmnet"){
    str(objModel$finalModel)
    objModel$finalModel$tuneValue # alpha=1 is lasso
    coefs<-coef(objModel$finalModel,objModel$bestTune$lambda)[,1]
    coefs<-coefs[coefs!=0]
}


## Test of difference btw two AUCs according to Hanley and McNeil (1983)
library(MKmisc)
?AUC.test

# Wilcoxon test is less refined, but it avoids the multiple pairwise tests required by the Hanley and McNeil test
```
