## Supplementary Code for "Pan-Cancer and Single-Cell modelling of genomic alterations through gene expression": 022_clouds.R

```
# TODO: Add comment
# 
# Author: fmg2117
###############################################################################


setwd("D:/Dropbox/projects/pancancer")

### Packages
library(ks)
library(tsne)

### Setup
tfs<-as.character(read.delim("../shared/genelists/tfgenes_2018_08_06.txt",as.is=TRUE)[,1])
cotfs<-as.character(read.delim("../shared/genelists/cotfgenes_2018_08_06.txt",as.is=TRUE)[,1])
sig<-as.character(read.delim("../shared/genelists/tfgenes_2018_08_06.txt",as.is=TRUE)[,1])
if(TRUE){
    raw<-matrix(c(
        "ACC","Adrenocortical Carcinoma",
        "BLCA","Bladder Urothelial Carcinoma",
        "BRCA","Breast Invasive Carcinoma",
        "CESC","Cervical Squamous Cell carcinoma",
        "CHOL","Cholangiocarcinoma",
        "COAD","Colon Adenocarcinoma",
        "DLBC","Diffuse Large B-cell Lymphoma",
        "ESCA","Esophageal Carcinoma",
        "GBM","Glioblastoma Multiforme",
        "HNSC","Head and Neck Squamous Cell Carcinoma",
        "KICH","Kidney Chromophobe",
        "KIRC","Kidney Renal Clear Cell Carcinoma",
        "KIRP","Kidney Renal Papillary Cell Carcinoma",
        "LAML","Acute Myeloid Leukemia",
        "LGG","Brain Lower Grade Glioma",
        "LIHC","Liver Hepatocellular Carcinoma",
        "LUAD","Lung Adenocarcinoma",
        "LUSC","Lung Squamous Cell Carcinoma",
        "MESO","Mesothelioma",
        "OV","Ovarian Serous Cystadenocarcinoma",
        "PAAD","Pancreatic Adenocarcinoma",
        "PCPG","Pheochromocytoma and Paraganglioma",
        "PRAD","Prostate Adenocarcinoma",
        "READ","Rectum Adenocarcinoma",
        "SARC","Sarcoma",
        "SKCM","Skin Cutaneous Melanoma",
        "STAD","Stomach Adenocarcinoma",
        "TGCT","Testicular Germ Cell Tumors",
        "THCA","Thyroid Carcinoma",
        "THYM","Thymoma",
        "UCEC","Uterine Corpus Endometrial Carcinoma",
        "UCS","Uterine Carcinosarcoma",
        "UVM","Uveal Melanoma"
    ),ncol=2,byrow = TRUE
    )
    tums<-setNames(raw[,2],raw[,1])
    rm(raw)
} # Tumor Acronyms
if(TRUE){
    # https://stackoverflow.com/questions/15282580/how-to-generate-a-number-of-most-distinctive-colors-in-r
    library(RColorBrewer)
    brewer.pal.info
    n<-length(tums)
    qual_col_pals<-brewer.pal.info[brewer.pal.info$category == 'qual',]
    cols<-unlist(mapply(brewer.pal, qual_col_pals$maxcolors, rownames(qual_col_pals)))
    set.seed(3)
    tumcols<-setNames(sample(cols, n),names(tums))
    #pie(rep(1,n),col=tumcols)
} # Tumor Colors


### Panmat
fname<-"results/022_panmat_expression_tfs.rda"
if(!file.exists(fname)){
    # Prepare the matrix
    matsamples<-c()
    matgenes<-c()
    mattypes<-c()
    tumorsizes<-matrix(NA,ncol=7,nrow=length(tums))
    colnames(tumorsizes)<-c("Acronym","Tumor","Primary Solid Tumor","Metastatic","Solid Tissue Normal","Primary Blood Derived Cancer","Other")
    tumorsizes[,1]<-names(tums);tumorsizes[,2]<-tums
    for(tum in names(tums)){
        message("Doing ",tum)
        filename<-paste0("tums010/",tum,"/",tum,"-expmat.rda")
        load(filename)
        
        # Only tumor samples
        # Tumor/Metastatic/Normal expression
        # https://gdc.cancer.gov/resources-tcga-users/tcga-code-tables/sample-type-codes
        t<-colnames(expmat)[grep("01",substr(colnames(expmat),14,15))] # Primary Solid Tumor
        m<-colnames(expmat)[grep("06",substr(colnames(expmat),14,15))] # Metastasic
        n<-colnames(expmat)[grep("11",substr(colnames(expmat),14,15))] # Solid Tissue Normal
        l<-colnames(expmat)[grep("03",substr(colnames(expmat),14,15))] # Primary Blood Derived Cancer
        texpmat<-expmat[,t]
        mexpmat<-expmat[,m]
        nexpmat<-expmat[,n]
        lexpmat<-expmat[,l]
        if(tum=="SKCM") texpmat<-cbind(texpmat,mexpmat)
        if(tum=="LAML") texpmat<-lexpmat
        tumorsizes[tumorsizes[,1]==tum,3]<-length(t)
        tumorsizes[tumorsizes[,1]==tum,4]<-length(m)
        tumorsizes[tumorsizes[,1]==tum,5]<-length(n)
        tumorsizes[tumorsizes[,1]==tum,6]<-length(l)
        tumorsizes[tumorsizes[,1]==tum,7]<-ncol(expmat)-length(t)-length(m)-length(n)-length(l)
        
        mat<-texpmat[intersect(tfs,rownames(texpmat)),]
        matgenes<-c(matgenes,rownames(mat))
        mattypes<-c(mattypes,rep(tum,ncol(mat)))
        matsamples<-c(matsamples,colnames(texpmat))
        rm(texpmat,mexpmat,nexpmat,lexpmat,expmat,mat,t,m,n,l)
    }
    write.csv(tumorsizes,file="results/022_tumorsizes.csv",quote=FALSE,row.names=FALSE)
    matgenes<-names(table(matgenes)[table(matgenes)==length(tums)])
    panmat<-matrix(NA,ncol=length(matsamples),nrow=length(matgenes))
    rownames(panmat)<-matgenes
    colnames(panmat)<-matsamples
    dim(panmat) # 1172 9642
    
    # Fill the matrix
    for(tum in names(tums)){
        message("Doing ",tum)
        filename<-paste0("tums010/",tum,"/",tum,"-expmat.rda")
        load(filename)
        
        # Only tumor samples
        # Only tumor samples
        # Tumor/Metastatic/Normal expression
        # https://gdc.cancer.gov/resources-tcga-users/tcga-code-tables/sample-type-codes
        t<-colnames(expmat)[grep("01",substr(colnames(expmat),14,15))] # Primary Solid Tumor
        m<-colnames(expmat)[grep("06",substr(colnames(expmat),14,15))] # Metastasic
        n<-colnames(expmat)[grep("11",substr(colnames(expmat),14,15))] # Solid Tissue Normal
        l<-colnames(expmat)[grep("03",substr(colnames(expmat),14,15))] # Primary Blood Derived Cancer
        texpmat<-expmat[,t]
        mexpmat<-expmat[,m]
        nexpmat<-expmat[,n]
        lexpmat<-expmat[,l]
        if(tum=="SKCM") texpmat<-cbind(texpmat,mexpmat)
        if(tum=="LAML") texpmat<-lexpmat
        
        mat<-texpmat[intersect(matgenes,rownames(texpmat)),]
        panmat[rownames(mat),colnames(mat)]<-mat[rownames(mat),colnames(mat)]
        rm(texpmat,mexpmat,nexpmat,lexpmat,expmat,mat,t,m,n,l)
    }
    
    save(panmat,mattypes,file=fname)
    
} else {load(fname)} # Generate a Pan-cancer matrix of all transcription factors expresison profiles

dim(panmat) # 1172 9642

### TSNE
fname<-"results/022_panmat_TSNE.rda"
if(!file.exists(fname)){
    set.seed(1)
    ttt<-tsne(t(panmat),max_iter=1000)
    rownames(ttt)<-mattypes
    samplenames<-colnames(panmat)
    save(ttt,samplenames,file=fname)
} else {
    load(fname)
}

##### Principal Components
#ppp<-prcomp(t(panmat))
#ttt<-ppp$x[,1:2]


####################################### PLOTS
### Parameters
margin<-5
gridsize<-1000

### Define plot locations
x<-ttt[,1]
y<-ttt[,2]
xlim<-range(x)
xlim<-xlim+diff(xlim)*c(-1,1)*margin/gridsize
ylim<-range(y)
ylim<-ylim+diff(ylim)*c(-1,1)*margin/gridsize
a<-c(xlim[1],ylim[1])
b<-1/c(diff(xlim),diff(ylim))
xnorm<-(x-a[1])*b[1]
ynorm<-(y-a[2])*b[2]


### Prepare 2d densities
den2d<-list()
for(tum in names(tums)){
    message("Estimating ",tum)
    tumden<-kde(cbind(x,y)[which(rownames(ttt)==tum),],gridsize=gridsize,xmin=c(xlim[1],ylim[1]),xmax=c(xlim[2],ylim[2]))$estimate
    den2d[[tum]]<-tumden
}
den2d<-lapply(den2d,function(x){x/max(x)})


# t1<-runif(10000,0,1)
# t2<-t1^10/2
# plot(t1,t2)
# 

##################### TSNE plot: pancancer ##################################
png("plots/022_TSNE_expmat.png",width=6000,height=3000,pointsize=60)
# Prepare plot area
plot(
    xnorm,
    ynorm,
    xlim=c(0,1.4),
    xlab="TSNE1",
    ylab="TSNE2",
    main="Expression-clustered pancancer dataset",
    type="n"
)
#rect(par("usr")[1], par("usr")[3], par("usr")[2], par("usr")[4], col = "grey98")
grid(col="black")
# Plot densities
for(itum in 1:length(tums)){
    message("Doing ",tums[itum])
    #alphas<-sort(0.5-as.vector(den2d[[itum]]^0.1/2))
    alphas<-(0:100)^2/100^2
    image(den2d[[itum]],add=TRUE,
          col=rgb(
              red=col2rgb(tumcols[itum])[1,1]/255,
              green=col2rgb(tumcols[itum])[2,1]/255,
              blue=col2rgb(tumcols[itum])[3,1]/255,
              alpha=alphas
          )
    )
}                     
legend("topright",pch=16,col=tumcols,legend=paste0(names(tums)," - ",tums," (",table(rownames(ttt)),")"),bg="white",cex=0.95)
# Plot points
points(
    xnorm,
    ynorm,
    col=tumcols[rownames(ttt)],
    pch=20,
    cex=0.5
)
# Centroid of tumor types
for(tum in names(tums)){
    xmean<-median(xnorm[rownames(ttt)==tum])
    ymean<-median(ynorm[rownames(ttt)==tum])
    text(xmean,ymean,labels=tum,font=2)
}
dev.off()


################ One TSNE per tumor type ###############
for(tum in names(tums)){
    ii<-which(names(tums)==tum)
    
    png(paste0("plots/022_tsne/022_TSNE_expmat_",tum,".png"),width=6000,height=3000,pointsize=60)
    # Prepare plot area
    plot(
        xnorm,
        ynorm,
        xlim=c(0,1.4),
        xlab="TSNE1",
        ylab="TSNE2",
        main="Expression-clustered pancancer dataset",
        type="n"
    )
    #rect(par("usr")[1], par("usr")[3], par("usr")[2], par("usr")[4], col = "grey98")
    grid(col="black")
    # Specific tumor density
    #alphas<-sort(0.5-as.vector(den2d[[ii]]^0.5/2))
    alphas<-(0:100)^2/100^2
    image(den2d[[ii]],add=TRUE,
          col=rgb(
              red=col2rgb(tumcols[ii])[1,1]/255,
              green=col2rgb(tumcols[ii])[2,1]/255,
              blue=col2rgb(tumcols[ii])[3,1]/255,
              alpha=alphas
          )
    )
    legendcols<-rep("grey95",length(tums))
    legendcols[ii]<-"black"
    legend("topright",pch=16,col=tumcols,legend=paste0(names(tums)," - ",tums," (",table(rownames(ttt)),")"),
           bg="white",cex=0.95,text.col=legendcols)
    # Plot points
    points(
        xnorm[which(rownames(ttt)==tum)],
        ynorm[which(rownames(ttt)==tum)],
        col="black",
        pch=20,
        cex=0.5
    )
    dev.off()
}
```
