## Supplementary Code for "Pan-Cancer and Single-Cell modelling of genomic alterations through gene expression": 023a_mostMutated.R

```
source("../shared/functions/geneids.R")
library(plotrix)
library(gplots)

matrix2col<-function(z,col1="white",col2="black",nbreaks=1000,center=FALSE,minmax=NULL){
    if(is.null(minmax)){
        if(center){
            extreme=max(abs(z))+0.001
            breaks <- seq(-extreme, extreme, length = nbreaks)
        }else {
            breaks <- seq(min(z), max(z), length = nbreaks)
        }
    } else {
        breaks <- seq(minmax[1],minmax[2], length = nbreaks)
    }
    
    ncol <- length(breaks) - 1
    col <- colorpanel(ncol,col1,col2)
    CUT <- cut(z, breaks=breaks,include.lowest = TRUE)
    colorlevels <- col[match(CUT, levels(CUT))] # assign colors to heights for each point
    names(colorlevels)<-rownames(z)
    
    colormatrix<-matrix(colorlevels,ncol=ncol(z),nrow=nrow(z))
    dimnames(colormatrix)<-dimnames(z)
    colormatrix[is.na(colormatrix)]<-col2 # Values above max values will not be included by CUT, so we color them as maximum values
    return(list(colormatrix=colormatrix,col=col))
}


### In the previous plot we showed the tumors in TCGA.
### Now we investigate the most mutated genes.


# Setup
if(TRUE){
    raw<-matrix(c(
        "ACC","Adrenocortical Carcinoma",
        "BLCA","Bladder Urothelial Carcinoma",
        "BRCA","Breast Invasive Carcinoma",
        "CESC","Cervical Squamous Cell carcinoma",
        "CHOL","Cholangiocarcinoma",
        "COAD","Colon Adenocarcinoma",
        "DLBC","Diffuse Large B-cell Lymphoma",
        "ESCA","Esophageal Carcinoma",
        "GBM","Glioblastoma Multiforme",
        "HNSC","Head and Neck Squamous Cell Carcinoma",
        "KICH","Kidney Chromophobe",
        "KIRC","Kidney Renal Clear Cell Carcinoma",
        "KIRP","Kidney Renal Papillary Cell Carcinoma",
        "LAML","Acute Myeloid Leukemia",
        "LGG","Brain Lower Grade Glioma",
        "LIHC","Liver Hepatocellular Carcinoma",
        "LUAD","Lung Adenocarcinoma",
        "LUSC","Lung Squamous Cell Carcinoma",
        "MESO","Mesothelioma",
        "OV","Ovarian Serous Cystadenocarcinoma",
        "PAAD","Pancreatic Adenocarcinoma",
        "PCPG","Pheochromocytoma and Paraganglioma",
        "PRAD","Prostate Adenocarcinoma",
        "READ","Rectum Adenocarcinoma",
        "SARC","Sarcoma",
        "SKCM","Skin Cutaneous Melanoma",
        "STAD","Stomach Adenocarcinoma",
        "TGCT","Testicular Germ Cell Tumors",
        "THCA","Thyroid Carcinoma",
        "THYM","Thymoma",
        "UCEC","Uterine Corpus Endometrial Carcinoma",
        "UCS","Uterine Carcinosarcoma",
        "UVM","Uveal Melanoma"
    ),ncol=2,byrow = TRUE
    )
    tums<-setNames(raw[,2],raw[,1])
    rm(raw)
} # Tumor Acronyms
if(TRUE){
    # https://stackoverflow.com/questions/15282580/how-to-generate-a-number-of-most-distinctive-colors-in-r
    library(RColorBrewer)
    brewer.pal.info
    n<-length(tums)
    qual_col_pals<-brewer.pal.info[brewer.pal.info$category == 'qual',]
    cols<-unlist(mapply(brewer.pal, qual_col_pals$maxcolors, rownames(qual_col_pals)))
    set.seed(3)
    tumcols<-setNames(sample(cols, n),names(tums))
    #pie(rep(1,n),col=tumcols)
} # Tumor Colors

# Load the mutSig blacklist
mutsig<-any2entrez(read.delim("../shared/genelists/mutSig_blacklist.txt",as.is=TRUE,header=FALSE)[,1])
# Check also http://massgenomics.org/2013/06/ngs-false-positives.html

# Most Mutated Genes in Individual Cancers
seengenes<-c()
for(tum in names(tums)){
    message("Doing ",tum)
    load(paste0("tums010/",tum,"/",tum,"-mutations.rda"))
    mutations<-mutations[!is.na(rownames(mutations)),]
    mutsums<-sort(apply(mutations,1,sum),dec=TRUE)
    x<-mutsums[1:30]
    names(x)<-eg2sym(names(x))
    png(paste0("plots/023a_mostMutated/mutations_",tum,".png"),w=3000,h=2000,p=40)
    par(las=2)
    bp<-barplot(100*x/ncol(mutations),ylab="% Mutated Samples",
                col=ifelse(names(x)%in%eg2sym(mutsig),"grey","black"),
                ylim=c(0,100),main=tums[tum]
    )
    par(las=1)
    mtext(paste0("Total samples: ",ncol(mutations)),cex=0.8)
    text(bp[,1],100*x/ncol(mutations),x,pos=3)
    dev.off()
    seengenes<-union(seengenes,rownames(mutations))
}
length(seengenes) # 20408 genes with at least one somatic mutations in at least one cancer (basically most of the genome)
if(TRUE){seengenes<-setdiff(seengenes,mutsig)} # Ignoring MutSig blacklisted genes
length(seengenes) # 20303 genes

### Prepare a panmut matrix
panmat<-matrix(0,nrow=length(seengenes),ncol=length(tums))
colnames(panmat)<-names(tums)
rownames(panmat)<-seengenes
dim(panmat) # 6626 genes with at least 10 mutations in at least one tumor type, 33 tumor types
hypermut<-setNames(rep(0,length(tums)),names(tums))
specialists<-setNames(rep(NA,length(tums)),names(tums))
for(tum in names(tums)){
    message("Doing ",tum)
    load(paste0("tums010/",tum,"/",tum,"-mutations.rda"))
    mutations<-mutations[!is.na(rownames(mutations)),]
    mutations<-mutations[intersect(rownames(mutations),seengenes),]
    freqhere<-apply(mutations,1,function(x){sum(x)/length(x)})
    seengenes_here<-intersect(seengenes,names(freqhere))
    panmat[seengenes_here,tum]<-freqhere[seengenes_here]

    sumhere<-apply(mutations,2,sum)
    hypermut[tum]<-sum(sumhere>=length(seengenes)*0.005)/ncol(mutations) # "Hypermutated" samples with more than 0.5% of the genes somatically mutated
    specialists[tum]<-names(freqhere)[which.max(freqhere)]
}
pancancer_frequency<-apply(panmat,1,function(x){sum(x)/length(x)})

### Genes to show
topgenes<-names(sort(pancancer_frequency,dec=TRUE))[1:25]
topgenes<-union(topgenes,specialists)
central<-panmat[topgenes,]
rownames(central)<-eg2sym(rownames(central))
rightbarplot<-pancancer_frequency[topgenes]
bottombarplot<-hypermut


###### Complicated layout ######
png("plots/023a_mostMutated.png",w=4000,h=4000,p=100)
lmatrix<-rbind(c(1,1,2),c(3,4,5),c(6,7,8))
layout(lmatrix,width=c(1,6,1),heights=c(1,6,2))

# 1- TITLE
par(mar = c(0,0,0,0))
texttitle<-"Frequent Somatic Mutations in Pan-Cancer Dataset"
plot(c(0, 1), c(0, 1), ann = F, bty = 'n', type = 'n', xaxt = 'n', yaxt = 'n')
text(x=0.5, y = 0.5, texttitle,cex = 1.6, col = "black")

# 2- TOP RULER
par(mar = c(0,0,0,0.4))
plot(1,xlim=c(0,0.4),ylim=c(0,0.4),ann=F,type="n",axes=FALSE)
segments(0,0.1,0.4,0.1)
segments(c(0:4)/10,0.1,c(0:4)/10,0)
text(0.2,0.3,labels="Pancancer\nFrequency (%)")
text(c(0:4)/10,0.15,labels=c(0,10,20,30,40),cex=0.9,xpd=NA)


# 3- LEFT NAMES
par(mar = c(0,0,0,0))
plot(1,xlim=c(0,1),ylim=c(0,1),ann=F,type="n",axes=FALSE)
text(1,0:(nrow(central)-1)/(nrow(central)-1),rev(rownames(central)),pos=2,col=ifelse(rev(rownames(central))%in%eg2sym(mutsig),"grey","black"))


# 4- CIRCLE MATRIX
par(mar = c(0,0,0,0))
plot(1,xlim=c(0,ncol(central)-1),ylim=c(0,nrow(central)-1),ann=F,type="n",axes=FALSE)
radius<-0.35
circlecols<-matrix2col(central,minmax=c(0,1))$colormatrix
circlecols<-circlecols[rev(rownames(circlecols)),]
for(i in 0:(nrow(central)-1)){
    for(j in 0:(ncol(central)-1)){
        draw.circle(j,i,radius=radius,col=circlecols[i+1,j+1])
    }
}


# 5- RIGHT BARPLOT
par(mar = c(0,0,0,0.4))
bp<-barplot(rev(rightbarplot),horiz=TRUE,add=TRUE,col="black",offset=0,axes=FALSE,plot=FALSE)
plot(1,xlim=c(0,0.4),ylim=c(min(bp),max(bp)),ann=F,type="n",axes=FALSE)
barplot(rev(rightbarplot),horiz=TRUE,add=TRUE,col="black",offset=0,axes=FALSE,names.arg=NA)


# 6- BOTTOM RULER
par(las=2,mar=c(4,0,0,0))
plot(1,xlim=c(0,1),ylim=c(0,1),ann=F,type="n",axes=FALSE)
segments(0.9,0,0.9,1)
segments(rep(0.9,6),0:5/5,rep(0.8,6),0:5/5)
text(0.5,-0.4,labels="HyperMutated\nSamples\n(%)",xpd=NA)
text(0.8,c(0:5)/5,labels=rev(c(0,20,40,60,80,100)),cex=0.9,xpd=NA,pos=2)


# 7- BOTTOM BARPLOT
par(las=2,mar=c(4,0,0,0))
bp<-barplot(bottombarplot,plot=FALSE)
barplot(bottombarplot,xlim=c(min(bp),max(bp)),ylim=c(max(bottombarplot),0),offset=0,axes=FALSE,col=tumcols)


# 8- DOT LEGEND
par(las=1,mar=c(0,0,0,0))
plot(1,xlim=c(0,5),ylim=c(0,13),ann=F,type="n",axes=FALSE)
text(2.5,12,"Mutated\nSamples")
ii<-0
icols<-colorpanel(6,"white","black")[2:6]
labels<-c("20%","40%","60%","80%","100%")
for(i in c(1,3,5,7,9)){
    ii<-ii+1
    draw.circle(1,i,radius=0.4,col=icols[ii])
    text(3,i,labels=labels[ii])
}


dev.off()
```
