## Supplementary Code for "Pan-Cancer and Single-Cell modelling of genomic alterations through gene expression": 024_models.R

```
# Idea
# Take all most frequent mutations/amplifications/deletions and model them with a simple glmnet binary classifier
# Plot AUROCs of results, grouped as mut/amp/del and tumor type (violin plot)
# Supp: correlation btw nr of occurrences and auroc (dot color is tumor type, pch is alteration type)
# Supp: mutsig blacklist vs. not blacklisted
options(java.parameters = "-Xmx16000m") # for printing via the xlsx package


# Setup
tfs<-as.character(read.delim("../shared/genelists/tfgenes_2018_08_06.txt",as.is=TRUE)[,1])
cotfs<-as.character(read.delim("../shared/genelists/cotfgenes_2018_08_06.txt",as.is=TRUE)[,1])
sig<-as.character(read.delim("../shared/genelists/tfgenes_2018_08_06.txt",as.is=TRUE)[,1])
if(TRUE){
    raw<-matrix(c(
        "ACC","Adrenocortical Carcinoma",
        "BLCA","Bladder Urothelial Carcinoma",
        "BRCA","Breast Invasive Carcinoma",
        "CESC","Cervical Squamous Cell carcinoma",
        "CHOL","Cholangiocarcinoma",
        "COAD","Colon Adenocarcinoma",
        "DLBC","Diffuse Large B-cell Lymphoma",
        "ESCA","Esophageal Carcinoma",
        "GBM","Glioblastoma Multiforme",
        "HNSC","Head and Neck Squamous Cell Carcinoma",
        "KICH","Kidney Chromophobe",
        "KIRC","Kidney Renal Clear Cell Carcinoma",
        "KIRP","Kidney Renal Papillary Cell Carcinoma",
        "LAML","Acute Myeloid Leukemia",
        "LGG","Brain Lower Grade Glioma",
        "LIHC","Liver Hepatocellular Carcinoma",
        "LUAD","Lung Adenocarcinoma",
        "LUSC","Lung Squamous Cell Carcinoma",
        "MESO","Mesothelioma",
        "OV","Ovarian Serous Cystadenocarcinoma",
        "PAAD","Pancreatic Adenocarcinoma",
        "PCPG","Pheochromocytoma and Paraganglioma",
        "PRAD","Prostate Adenocarcinoma",
        "READ","Rectum Adenocarcinoma",
        "SARC","Sarcoma",
        "SKCM","Skin Cutaneous Melanoma",
        "STAD","Stomach Adenocarcinoma",
        "TGCT","Testicular Germ Cell Tumors",
        "THCA","Thyroid Carcinoma",
        "THYM","Thymoma",
        "UCEC","Uterine Corpus Endometrial Carcinoma",
        "UCS","Uterine Carcinosarcoma",
        "UVM","Uveal Melanoma"
    ),ncol=2,byrow = TRUE
    )
    tums<-setNames(raw[,2],raw[,1])
    rm(raw)
} # Tumor Acronyms
if(TRUE){
    # https://stackoverflow.com/questions/15282580/how-to-generate-a-number-of-most-distinctive-colors-in-r
    library(RColorBrewer)
    brewer.pal.info
    n<-length(tums)
    qual_col_pals<-brewer.pal.info[brewer.pal.info$category == 'qual',]
    cols<-unlist(mapply(brewer.pal, qual_col_pals$maxcolors, rownames(qual_col_pals)))
    set.seed(3)
    tumcols<-setNames(sample(cols, n),names(tums))
    #pie(rep(1,n),col=tumcols)
} # Tumor Colors

library(beeswarm)
library(vioplot)
library(caret)
library(pROC)


# Load the mutSig blacklist
source("../shared/functions/geneids.R")
mutsig<-any2entrez(read.delim("../shared/genelists/mutSig_blacklist.txt",as.is=TRUE,header=FALSE)[,1])


### Mega model loop

# Sample sizes
nsamples<-matrix(0,nrow=length(tums),ncol=2)
rownames(nsamples)<-names(tums);colnames(nsamples)<-c("mut","cnv")
for(tum in names(tums)){
    message("Doing ",tum)
    load(paste0("tums010/",tum,"/",tum,"-mutations.rda"))
    load(paste0("tums010/",tum,"/",tum,"-fcnv.rda"))
    load(paste0("tums010/",tum,"/",tum,"-expmat.rda"))
    colnames(mutations)<-substr(colnames(mutations),1,15)
    colnames(fcnv)<-substr(colnames(fcnv),1,15)
    colnames(expmat)<-substr(colnames(expmat),1,15)
    nsamples[tum,"mut"]<-length(intersect(colnames(mutations),colnames(expmat)))
    nsamples[tum,"cnv"]<-length(intersect(colnames(fcnv),colnames(expmat)))
}    

# In order to have reliable models, we will limit our analysis to tumor datasets with at least 100 samples (intersection expression/mutation
# or expression/cnv) and to mutations/cnvs with at least 10 cases, present in at least 5% of the samples but no more than 95% of the samples
bigtums<-rownames(nsamples)[nsamples[,1]>=100&nsamples[,1]>=100]
write.csv(nsamples,file="results/024_intersectionExpmatWithAlterations.csv")

# Test models
method<-"gbm"


# Big model loop (run on the clusters)
if(TRUE){
    # Mutations
    for(tum in bigtums){
        fname<-paste0("results/024_aurocs/",tum,"-mut_aurocs.rda")
        if(!file.exists(fname)){
            message("Doing ",tum)
            load(paste0("tums010/",tum,"/",tum,"-mutations.rda"))
            load(paste0("tums010/",tum,"/",tum,"-expmat.rda"))
            colnames(mutations)<-substr(colnames(mutations),1,15)
            colnames(expmat)<-substr(colnames(expmat),1,15)
            
            # Creates a data frame with variables (genes) and observations (samples) that includes the mutations data for the specific mutation
            int.samples<-intersect(colnames(expmat),colnames(mutations))
            mutations.a<-mutations[,int.samples]
            expmat.a<-expmat[,int.samples]
            # Filter by Variance (Top 1000)
            evars<-apply(expmat.a,1,var)
            expmat.a<-expmat.a[evars>0,]
            expmat.a<-expmat.a[names(sort(evars,dec=TRUE)[1:1000]),]
            
            
            # Filter only genes with at least 10 events in dataset
            mutsum<-apply(mutations.a,1,sum)
            mutgenes<-names(mutsum[mutsum>=10])
            
            aurocs<-c()
            pb<-txtProgressBar(0,length(mutgenes),style=3)
            for(igene in 1:length(mutgenes)){
                setTxtProgressBar(pb,igene)
                gene<-mutgenes[igene]
                gene.mut<-as.vector(mutations.a[as.character(gene),])
                gene.mut<-ifelse(gene.mut==1,"mut","wt")
                
                # Build data object
                df<-cbind(t(expmat.a),gene.mut)
                df<-as.data.frame(df)
                df$gene.mut<-as.factor(df$gene.mut)
                
                
                # Avoid test if samples are too few or if percmut is not within optimal range (5-95%)
                percmut<-signif(100*sum(as.vector(mutations.a[as.character(gene),]))/nrow(df),4)
                #message(tum,": testing on ",nrow(df)," samples for ",eg2sym(gene)," (",percmut,"% mutated)")
                if(percmut<5|percmut>95){next}
                
                
                # Many caret functions model the probability of the first factor level
                df$gene.mut<-relevel(df$gene.mut,ref="mut")
                
                
                # We want to predict the outcome (the mutations), from a set of predictors (gene expression data), which we code here
                # prop.outcome<-prop.table(table(df[,"gene.mut"]))
                outcomeName<-"gene.mut"
                predictorsNames<-colnames(df)[colnames(df)!=outcomeName]
                
                # Set a seed to use, to make the CV steps reproducible
                set.seed(1)
                
                # Split the data into training and test sets (0.6, 0.8, try different ones?)
                splitIndex<-createDataPartition(df[,outcomeName],p=0.75,list=FALSE,times=1)
                trainDF<-df[splitIndex,]
                testDF<-df[-splitIndex,]
                
                # Train Model Parameters (10-fold CV is acceptable)
                objControl<-trainControl(method="repeatedcv",number=10,returnResamp='none',summaryFunction=twoClassSummary,classProbs=TRUE)
                objModel<-train(data.matrix(trainDF[,predictorsNames]),trainDF[,outcomeName], 
                                method=method, 
                                trControl=objControl,  
                                metric="ROC",verbose=FALSE
                )
                
                # Validating the Model
                # It will predict using the best model: objModel$finalModel
                predictions<-predict(object=objModel,data.matrix(testDF[,predictorsNames]))
                probabilities<-predict(object=objModel,data.matrix(testDF[,predictorsNames]),type="prob")
                rocCurve<-roc(response=testDF$gene.mut,predictor=probabilities[,"mut"])
                
                # Confusion matrix
                #confusionMatrix(predictions,testDF$gene.mut)
                
                # # Plot the ROC Curve with AUC
                # png(paste0("plots/024_modelROCs/ROC_",tum,"_MUT_",eg2sym(gene),".png"),w=1400,h=1000,p=35)
                # plot(rocCurve,col="red3",cex.lab=2,cex.main=1.1,print.auc=FALSE,lwd=2,
                #      main=paste0(method," Model predicting ",eg2sym(gene)," Mutations in ",tum," dataset"))
                # mtext("10-fold CV",side=1)
                # grid(col="lightgrey")
                # legend(0.2,0.2,legend=paste0("AUC= ", round(rocCurve$auc,4)))
                # dev.off()
                
                # Update auroc object
                aurocs<-c(aurocs,rocCurve$auc)
                names(aurocs)[length(aurocs)]<-gene
                
                # # Extract predictors from objModel$finalModel
                # if(method=="glmnet"){
                #     str(objModel$finalModel)
                #     objModel$finalModel$tuneValue # alpha=1 is lasso
                #     coefs<-coef(objModel$finalModel,objModel$bestTune$lambda)[,1]
                #     coefs<-coefs[coefs!=0]
                # }
                
                
                
                # ## Test of difference btw two AUCs according to Hanley and McNeil (1983)
                # library(MKmisc)
                # ?AUC.test
            }
            save(aurocs,file=fname)
        }
    }
    
    # Amplifications
    for(tum in rev(bigtums)){
        fname<-paste0("results/024_aurocs/",tum,"-amp_aurocs.rda")
        if(!file.exists(fname)){
            message("Doing ",tum)
            load(paste0("tums010/",tum,"/",tum,"-fcnv.rda"))
            mutations<-fcnv
            mutations[fcnv>=0.5]<-1
            mutations[fcnv<0.5]<-0
            mutations<-mutations[!is.na(rownames(mutations)),]
            
            # Single event case
            singlecase<-FALSE
            if(!is.matrix(mutations)){
                singlecase<-TRUE
                mutations<-t(as.matrix(mutations))
                rownames(mutations)<-rownames(fcnv)
            }
            if(nrow(mutations)==0){next}
            
            load(paste0("tums010/",tum,"/",tum,"-expmat.rda"))
            colnames(mutations)<-substr(colnames(mutations),1,15)
            colnames(expmat)<-substr(colnames(expmat),1,15)
            
            
            # Creates a data frame with variables (genes) and observations (samples) that includes the mutations data for the specific mutation
            int.samples<-intersect(colnames(expmat),colnames(mutations))
            mutations.a<-mutations[,int.samples]
            expmat.a<-expmat[,int.samples]
            # Filter by Variance (Top 1000)
            evars<-apply(expmat.a,1,var)
            expmat.a<-expmat.a[evars>0,]
            expmat.a<-expmat.a[names(sort(evars,dec=TRUE)[1:1000]),]
            
            if(singlecase){
                mutations.a<-t(as.matrix(mutations.a))
                rownames(mutations.a)<-rownames(mutations)
            }
            
            if(!singlecase){
                # Filter only genes with at least 10 events in dataset
                mutsum<-apply(mutations.a,1,sum)
                mutgenes<-names(mutsum[mutsum>=10])
                if(tum=="THYM"){
                    mutgenes<-names(mutsum[mutsum>=5]) # Otherwise we have zero events here
                }
                
                # Some genes have identical CNV profiles
                cormat<-cor(t(fcnv[mutgenes,]))
                threshold<-1
                hits<-which(cormat==threshold,arr.ind=TRUE)
                hits<-hits[apply(hits,1,function(x){if(x[1]==x[2]){return(FALSE)}else{return(TRUE)}}),]
                hits1<-rownames(cormat)[hits[,1]]
                hits2<-colnames(cormat)[hits[,2]]
                hits<-cbind(hits1,hits2)
                identinet<-list()
                for (u in unique(as.vector(hits))){
                    identinet[[u]]<-unique(c(hits[hits[,1]==u,2],hits[hits[,2]==u,1]))
                }
            } else {
                identinet<-list()
                mutgenes<-rownames(mutations)
            }
            
            # Fill aurocs vector
            aurocs<-c()
            pb<-txtProgressBar(0,length(mutgenes),style=3)
            for(igene in 1:length(mutgenes)){
                setTxtProgressBar(pb,igene)
                gene<-mutgenes[igene]
                otheridenticals<-identinet[[gene]]
                if(any(otheridenticals%in%names(aurocs))){
                    takeaurocfromthis<-aurocs[which(otheridenticals%in%names(aurocs))]
                    aurocs<-c(aurocs,takeaurocfromthis)
                    names(aurocs)[length(aurocs)]<-gene
                    next
                }
                
                gene.mut<-as.vector(mutations.a[as.character(gene),])
                gene.mut<-ifelse(gene.mut==1,"mut","wt")
                
                # Build data object
                df<-cbind(t(expmat.a),gene.mut)
                df<-as.data.frame(df)
                df$gene.mut<-as.factor(df$gene.mut)
                
                
                # Avoid test if samples are too few or if percmut is not within optimal range (5-95%)
                percmut<-signif(100*sum(as.vector(mutations.a[as.character(gene),]))/nrow(df),4)
                #message(tum,": testing on ",nrow(df)," samples for ",eg2sym(gene)," (",percmut,"% mutated)")
                if(tum!="THYM"){
                    if(percmut<5|percmut>95){next}
                }
                
                
                # Many caret functions model the probability of the first factor level
                df$gene.mut<-relevel(df$gene.mut,ref="mut")
                
                
                # We want to predict the outcome (the mutations), from a set of predictors (gene expression data), which we code here
                # prop.outcome<-prop.table(table(df[,"gene.mut"]))
                outcomeName<-"gene.mut"
                predictorsNames<-colnames(df)[colnames(df)!=outcomeName]
                
                # Set a seed to use, to make the CV steps reproducible
                set.seed(1)
                
                # Split the data into training and test sets (0.6, 0.8, try different ones?)
                splitIndex<-createDataPartition(df[,outcomeName],p=0.75,list=FALSE,times=1)
                trainDF<-df[splitIndex,]
                testDF<-df[-splitIndex,]
                
                # Train Model Parameters (10-fold CV is acceptable)
                objControl<-trainControl(method="repeatedcv",number=10,returnResamp='none',summaryFunction=twoClassSummary,classProbs=TRUE)
                objModel<-train(data.matrix(trainDF[,predictorsNames]),trainDF[,outcomeName], 
                                method=method, 
                                trControl=objControl,  
                                metric="ROC",verbose=FALSE
                )
                
                # Validating the Model
                # It will predict using the best model: objModel$finalModel
                predictions<-predict(object=objModel,data.matrix(testDF[,predictorsNames]))
                probabilities<-predict(object=objModel,data.matrix(testDF[,predictorsNames]),type="prob")
                rocCurve<-roc(response=testDF$gene.mut,predictor=probabilities[,"mut"])
                
                # Confusion matrix
                #confusionMatrix(predictions,testDF$gene.mut)
                
                # # Plot the ROC Curve with AUC
                # png(paste0("plots/024_modelROCs/ROC_",tum,"_AMP_",eg2sym(gene),".png"),w=1400,h=1000,p=35)
                # plot(rocCurve,col="red3",cex.lab=2,cex.main=1.1,print.auc=FALSE,lwd=2,
                #      main=paste0(method," Model predicting ",eg2sym(gene)," Amplifications in ",tum," dataset"))
                # mtext("10-fold CV",side=1)
                # grid(col="lightgrey")
                # legend(0.2,0.2,legend=paste0("AUC= ", round(rocCurve$auc,4)))
                # dev.off()
                
                # Update auroc object
                aurocs<-c(aurocs,rocCurve$auc)
                names(aurocs)[length(aurocs)]<-gene
                
                # # Extract predictors from objModel$finalModel
                # if(method=="glmnet"){
                #     str(objModel$finalModel)
                #     objModel$finalModel$tuneValue # alpha=1 is lasso
                #     coefs<-coef(objModel$finalModel,objModel$bestTune$lambda)[,1]
                #     coefs<-coefs[coefs!=0]
                # }
                
                
                
                # ## Test of difference btw two AUCs according to Hanley and McNeil (1983)
                # library(MKmisc)
                # ?AUC.test
            }
            save(aurocs,file=fname)
        }
    }   
    
    # Deletions
    for(tum in bigtums){
        fname<-paste0("results/024_aurocs/",tum,"-del_aurocs.rda")
        if(!file.exists(fname)){
            message("Doing ",tum)
            load(paste0("tums010/",tum,"/",tum,"-fcnv.rda"))
            mutations<-fcnv
            mutations[fcnv<=(-0.5)]<-1
            mutations[fcnv>(-0.5)]<-0
            mutations<-mutations[!is.na(rownames(mutations)),]
            
            
            # Single event case
            singlecase<-FALSE
            if(!is.matrix(mutations)){
                singlecase<-TRUE
                mutations<-t(as.matrix(mutations))
                rownames(mutations)<-rownames(fcnv)
            }
            
            if(nrow(mutations)==0){next}
            
            load(paste0("tums010/",tum,"/",tum,"-expmat.rda"))
            colnames(mutations)<-substr(colnames(mutations),1,15)
            colnames(expmat)<-substr(colnames(expmat),1,15)
            
            
            # Creates a data frame with variables (genes) and observations (samples) that includes the mutations data for the specific mutation
            int.samples<-intersect(colnames(expmat),colnames(mutations))
            mutations.a<-mutations[,int.samples]
            expmat.a<-expmat[,int.samples]
            # Filter by Variance (Top 1000)
            evars<-apply(expmat.a,1,var)
            expmat.a<-expmat.a[evars>0,]
            expmat.a<-expmat.a[names(sort(evars,dec=TRUE)[1:1000]),]
            
            if(singlecase){
                mutations.a<-t(as.matrix(mutations.a))
                rownames(mutations.a)<-rownames(mutations)
            }
            
            if(!singlecase){
                # Filter only genes with at least 10 events in dataset
                mutsum<-apply(mutations.a,1,sum)
                mutgenes<-names(mutsum[mutsum>=10])
                
                if(tum=="THYM"){
                    mutgenes<-names(mutsum[mutsum>=5]) # Otherwise we have zero events here
                }
                
                # Some genes have identical CNV profiles
                cormat<-cor(t(fcnv[mutgenes,]))
                threshold<-1
                hits<-which(cormat==threshold,arr.ind=TRUE)
                hits<-hits[apply(hits,1,function(x){if(x[1]==x[2]){return(FALSE)}else{return(TRUE)}}),]
                hits1<-rownames(cormat)[hits[,1]]
                hits2<-colnames(cormat)[hits[,2]]
                hits<-cbind(hits1,hits2)
                identinet<-list()
                for (u in unique(as.vector(hits))){
                    identinet[[u]]<-unique(c(hits[hits[,1]==u,2],hits[hits[,2]==u,1]))
                }
            } else {
                mutgenes<-rownames(mutations)
            }
            
            
            # Fill aurocs vector
            aurocs<-c()
            pb<-txtProgressBar(0,length(mutgenes),style=3)
            for(igene in 1:length(mutgenes)){
                setTxtProgressBar(pb,igene)
                gene<-mutgenes[igene]
                otheridenticals<-identinet[[gene]]
                if(any(otheridenticals%in%names(aurocs))){
                    takeaurocfromthis<-aurocs[which(otheridenticals%in%names(aurocs))]
                    aurocs<-c(aurocs,takeaurocfromthis)
                    names(aurocs)[length(aurocs)]<-gene
                    next
                }
                
                gene.mut<-as.vector(mutations.a[as.character(gene),])
                gene.mut<-ifelse(gene.mut==1,"mut","wt")
                
                # Build data object
                df<-cbind(t(expmat.a),gene.mut)
                df<-as.data.frame(df)
                df$gene.mut<-as.factor(df$gene.mut)
                
                
                # Avoid test if samples are too few or if percmut is not within optimal range (5-95%)
                percmut<-signif(100*sum(as.vector(mutations.a[as.character(gene),]))/nrow(df),4)
                #message(tum,": testing on ",nrow(df)," samples for ",eg2sym(gene)," (",percmut,"% mutated)")
                if(tum!="THYM"){
                    if(percmut<5|percmut>95){next}
                }
                
                
                # Many caret functions model the probability of the first factor level
                df$gene.mut<-relevel(df$gene.mut,ref="mut")
                
                
                # We want to predict the outcome (the mutations), from a set of predictors (gene expression data), which we code here
                # prop.outcome<-prop.table(table(df[,"gene.mut"]))
                outcomeName<-"gene.mut"
                predictorsNames<-colnames(df)[colnames(df)!=outcomeName]
                
                # Set a seed to use, to make the CV steps reproducible
                set.seed(1)
                
                # Split the data into training and test sets (0.6, 0.8, try different ones?)
                splitIndex<-createDataPartition(df[,outcomeName],p=0.75,list=FALSE,times=1)
                trainDF<-df[splitIndex,]
                testDF<-df[-splitIndex,]
                
                # Train Model Parameters (10-fold CV is acceptable)
                objControl<-trainControl(method="repeatedcv",number=10,returnResamp='none',summaryFunction=twoClassSummary,classProbs=TRUE)
                objModel<-train(data.matrix(trainDF[,predictorsNames]),trainDF[,outcomeName], 
                                method=method, 
                                trControl=objControl,  
                                metric="ROC",verbose=FALSE
                )
                
                # Validating the Model
                # It will predict using the best model: objModel$finalModel
                predictions<-predict(object=objModel,data.matrix(testDF[,predictorsNames]))
                probabilities<-predict(object=objModel,data.matrix(testDF[,predictorsNames]),type="prob")
                rocCurve<-roc(response=testDF$gene.mut,predictor=probabilities[,"mut"])
                
                # Confusion matrix
                #confusionMatrix(predictions,testDF$gene.mut)
                
                # # Plot the ROC Curve with AUC
                # png(paste0("plots/024_modelROCs/ROC_",tum,"_DEL_",eg2sym(gene),".png"),w=1400,h=1000,p=35)
                # plot(rocCurve,col="red3",cex.lab=2,cex.main=1.1,print.auc=FALSE,lwd=2,
                #      main=paste0(method," Model predicting ",eg2sym(gene)," Deletions in ",tum," dataset"))
                # mtext("10-fold CV",side=1)
                # grid(col="lightgrey")
                # legend(0.2,0.2,legend=paste0("AUC= ", round(rocCurve$auc,4)))
                # dev.off()
                
                # Update auroc object
                aurocs<-c(aurocs,rocCurve$auc)
                names(aurocs)[length(aurocs)]<-gene
                
                # # Extract predictors from objModel$finalModel
                # if(method=="glmnet"){
                #     str(objModel$finalModel)
                #     objModel$finalModel$tuneValue # alpha=1 is lasso
                #     coefs<-coef(objModel$finalModel,objModel$bestTune$lambda)[,1]
                #     coefs<-coefs[coefs!=0]
                # }
                
                
                
                # ## Test of difference btw two AUCs according to Hanley and McNeil (1983)
                # library(MKmisc)
                # ?AUC.test
            }
            save(aurocs,file=fname)
        }
    }   
}


### We plot the number of NONIDENTICAL CNV tracks


# Aggregate auroc results
results<-list()
for(tum in bigtums){
    subresults<-list()
    for(type in c("mut","amp","del")){
        fname<-paste0("results/024_aurocs/",tum,"-",type,"_aurocs.rda")
        if(file.exists(fname)){
            load(fname)
            if(type=="mut"){
                subresults[[type]]<-aurocs
            } else {
                subresults[[type]]<-aurocs[!duplicated(aurocs)]
            }
        }else{
            subresults[[type]]<-c(NA)
        }
    }
    results[[tum]]<-subresults
}
save(results,file="results/024_aurocs_results.rda")

# Print AUROCs as supplementary table
library(xlsx)
it<-1
for(tum in names(results)){
    for(type in names(results[[tum]])){
        message("Doing ",paste0(tum,"_",type))
        if(it==1){
            it<-2
            append<-FALSE
        } else {append<-TRUE}
        vector<-sort(results[[tum]][[type]],dec=TRUE)
        if(length(vector>0)){
            dataframe<-data.frame(Gene=eg2sym(names(vector)),AUROC=vector)
            write.xlsx(dataframe,file="results/024_aurocs.xlsx",append=append,
                       sheetName=paste0(tum,"_",type),row.names=FALSE)
        }
    }
}


# Show TP53
# Explain that in PAAD KRAS is the dominant mutation and so TP53 models suffer from that


# Violin plot function
# Featuring semitransparent colors, jittered beeswarmed points and intelligent optional labeling
# input<-list()
# for(tum in names(tums)[1:8]){
#     input[[tum]]<-list(mut=abs(rnorm(1000,mean=1)),amp=abs(rnorm(1000,mean=1)),del=abs(rnorm(1000,mean=1)))
# }
violin<-function(input,flag1=NULL,flag2=NULL,legend=TRUE){
    cols<-c("grey50","red3","cornflowerblue")
    transpcol<-rgb(255,255,255,max=255,alpha=0,names="blue50")
    uex<-unlist(input)
    ss<-length(input[[1]])
    
    par(mar=c(2,3.4,1,1))
    plot(1,xlim=c(0,(ss+1)*length(input)),ylim=c(0.2,1.2),ann=F,type="n",axes="FALSE")
    axis(2,at=pretty(0:1),labels=pretty(0:1))
    mtext("AUROC",side=2,line=2)
    ii<-1
    set.seed(1) # for beeswarm placement
    for(i in 1:length(input)){
        text(ii+1,0.11,names(input)[i],xpd=NA)
        subinput<-input[[i]]
        for(is in 1:length(subinput)){
            if(!is.na(subinput[[is]][1])){
                vioplot(subinput[[is]],add=TRUE,at=ii,drawRect=FALSE,wex=1.1,border=NA,
                        col=rgb(col2rgb(cols[is])[1]/255,col2rgb(cols[is])[2]/255,col2rgb(cols[is])[3]/255,alpha=0.2))
                bs<-beeswarm(subinput[[is]],pch=20,add=TRUE,at=ii,col=cols[is],spacing=1)
                boxplot(subinput[[is]],add=TRUE,at=ii,lwd=4,pch=NA,whisklty=0,staplelty=0,width=1.5,col=transpcol)
                if(!is.null(flag1)){
                    iflag<-which(names(subinput[[is]])==flag1)
                    points(bs$x[iflag],bs$y[iflag],pch="o",cex=2)
                }
                if(!is.null(flag2)){
                    iflag<-which(names(subinput[[is]])==flag2)
                    points(bs$x[iflag],bs$y[iflag],pch="x",cex=2)
                }
                text(ii,max(subinput[[is]]),labels=length(subinput[[is]]),pos=3)
            }
            ii<-ii+1
        }
        ii<-ii+1
    }
    abline(h=0.5,lty=2)
    if(legend){
        legend("topleft",col=cols,pch=20,pt.cex=2,legend=c("Mutations","Amplifications","Deletions"))
        if((!is.null(flag1))&(!is.null(flag2))){
            legend("topright",col="black",pch=c("o","x"),pt.cex=2,legend=eg2sym(c(flag1,flag2)))
        }
    }
}


flag1=any2entrez("TP53");flag2=any2entrez("KRAS")

png("plots/024_violin.png",w=6000,h=6000,p=120)
par(mfrow=c(3,1))
violin(results[c(1:8)],flag1=flag1,flag2=flag2)
violin(results[c(9:16)],flag1=flag1,flag2=flag2,legend=FALSE)
violin(results[c(17:24)],flag1=flag1,flag2=flag2,legend=FALSE)
dev.off()


# Highlight
# BRAF in THCA
# Whatever is in THYM
# Top genes in violin
```
