## Supplementary Code for "Pan-Cancer and Single-Cell modelling of genomic alterations through gene expression": 024b_models_oncineca.R

```
# We need expmat, mutations, fcnv, tfgenes, mutSigblacklist, geneids.R
# put everything on input


cd /gpfs/work/IscrC_tumornet/pancancer


# In order to have reliable models, we will limit our analysis to tumor datasets with at least 100 samples (intersection expression/mutation
# or expression/cnv) and to mutations/cnvs with at least 10 cases, present in at least 5% of the samples but no more than 95% of the samples

script=024b_models_script.R
for type in mut amp del
do
for tum in BLCA BRCA CESC COAD ESCA GBM HNSC KIRC KIRP LAML LGG LIHC LUAD LUSC OV PAAD PCPG PRAD SARC SKCM STAD TGCT THCA THYM
do
qlogo=/gpfs/scratch/userexternal/fgiorgi0/qlogs/${type}_${tum}_out.txt
qloge=/gpfs/scratch/userexternal/fgiorgi0/qlogs/${type}_${tum}_err.txt
command="Rscript $script --vanilla --args $tum $type"
sbatch --wrap "$command" -o $qlogo -e $qloge -J ${type}_${tum} -t 24:00:00 --mem=8000M -p gll_usr_prod -A iscrc_tumornet
done
done


scp:/gpfs/work/IscrC_tumornet/pancancer/output/*  /mnt/d/Dropbox/projects/pancancer/results/024_aurocs/
```
