## Supplementary Code for "Pan-Cancer and Single-Cell modelling of genomic alterations through gene expression": 025_compareModelTypes.R

```
# Run on galileo
cd /gpfs/work/IscrC_tumornet/pancancer
srun -J models -t 24:00:00 --mem=16000M -p gll_usr_prod -A iscrc_tumornet --pty /bin/bash
R

## Pick a mutation set from one of the cancer in the previous figure and rerun the analysis with 
## other method types
libraries<-c("caret","pROC")
lapply(libraries,library,character.only=TRUE)

# # Neural Networks mxnet
# cran <- getOption("repos")
# cran["dmlc"] <- "https://s3-us-west-2.amazonaws.com/apache-mxnet/R/CRAN/"
# options(repos = cran)
# install.packages("mxnet",dependencies = T)
# library(mxnet)


## Methods to be used
methods<-c("bayesglm","evtree","gbm","glm","kknn","lda","mxnet","pcaNNet","rf","svmLinear","svmRadial")

# Bayesian Generalized Linear Model: bayesglm
# Elasticnet: enet ### NOT FOR CLASSIFICATION
# Ensembles of Generalized Linear Models: randomGLM ### CRASH, RECHECK
# Gradient Boost Modeling: gbm
# Generalized Linear Model: glm
# Generalized Linear Model: glmnet ### CRASH
# Generalized Linear Model: glmnet_h2o ### CRASH
# k-Nearest Neighbors: kknn
# Linear Discriminant Analysis: lda
# Naive Bayes: nb # CRASH
# Neural Network: mxnet
# Neural Network: mxnetAdam
# Neural Network: neuralnet ### NOT FOR CLASSIFICATION
# Neural Network: nnet ### CRASH
# Neural Network with Feature Extraction: pcaNNet
# Non-Informative Movel: null ### CRASH
# Quadratic Discriminant Analysis: qda ### CRASH
# Random Forest: rf
# Robust Linear model: rlm ### NOT FOR CLASSIFICATION
# Support Vector Machine: svmLinear
# Support Vector Machine: svmRadial
# Tree Models from Genetic Algorithms: evtree

run<-function(){
    tum<-"BLCA"
    for(method in methods){
        fname<-paste0("results/025_compareModelTypes/",tum,"-mut_",method,"_aurocs.rda")
        if(!file.exists(fname)){
            message("Doing ",method)
            load(paste0("tums010/",tum,"/",tum,"-mutations.rda"))
            load(paste0("tums010/",tum,"/",tum,"-expmat.rda"))
            #load(paste0("input/",tum,"-mutations.rda"))
            #load(paste0("input/",tum,"-expmat.rda"))
            colnames(mutations)<-substr(colnames(mutations),1,15)
            colnames(expmat)<-substr(colnames(expmat),1,15)
            
            # Creates a data frame with variables (genes) and observations (samples) that includes the mutations data for the specific mutation
            int.samples<-intersect(colnames(expmat),colnames(mutations))
            mutations.a<-mutations[,int.samples]
            expmat.a<-expmat[,int.samples]
            # Filter by Variance (Top 1000)
            evars<-apply(expmat.a,1,var)
            expmat.a<-expmat.a[evars>0,]
            expmat.a<-expmat.a[names(sort(evars,dec=TRUE)[1:1000]),]
            
            
            # Filter only genes with at least 10 events in dataset
            mutsum<-apply(mutations.a,1,sum)
            mutgenes<-names(mutsum[mutsum>=10])
            
            aurocs<-c()
            pb<-txtProgressBar(0,length(mutgenes),style=3)
            for(igene in 1:length(mutgenes)){
                setTxtProgressBar(pb,igene)
                gene<-mutgenes[igene]
                gene.mut<-as.vector(mutations.a[as.character(gene),])
                gene.mut<-ifelse(gene.mut==1,"mut","wt")
                
                # Build data object
                df<-cbind(t(expmat.a),gene.mut)
                df<-as.data.frame(df)
                df$gene.mut<-as.factor(df$gene.mut)
                
                
                # Avoid test if samples are too few or if percmut is not within optimal range (5-95%)
                percmut<-signif(100*sum(as.vector(mutations.a[as.character(gene),]))/nrow(df),4)
                #message(tum,": testing on ",nrow(df)," samples for ",eg2sym(gene)," (",percmut,"% mutated)")
                if(percmut<5|percmut>95){next}
                
                
                # Many caret functions model the probability of the first factor level
                df$gene.mut<-relevel(df$gene.mut,ref="mut")
                
                
                # We want to predict the outcome (the mutations), from a set of predictors (gene expression data), which we code here
                # prop.outcome<-prop.table(table(df[,"gene.mut"]))
                outcomeName<-"gene.mut"
                predictorsNames<-colnames(df)[colnames(df)!=outcomeName]
                
                # Set a seed to use, to make the CV steps reproducible
                set.seed(1)
                
                # Split the data into training and test sets (0.6, 0.8, try different ones?)
                splitIndex<-createDataPartition(df[,outcomeName],p=0.75,list=FALSE,times=1)
                trainDF<-df[splitIndex,]
                testDF<-df[-splitIndex,]
                
                # Train Model Parameters (10-fold CV is acceptable)
                objControl<-trainControl(method="repeatedcv",number=10,returnResamp='none',summaryFunction=twoClassSummary,classProbs=TRUE)
                # methods<-c("bayesglm","gbm","glmnet","lda","nnet","rf","svmLinear","svmRadial")
                objModel<-train(data.matrix(trainDF[,predictorsNames]),trainDF[,outcomeName], 
                                method=method, 
                                trControl=objControl,  
                                metric="ROC"
                )
                
                # Validating the Model
                # It will predict using the best model: objModel$finalModel
                predictions<-predict(object=objModel,data.matrix(testDF[,predictorsNames]))
                probabilities<-predict(object=objModel,data.matrix(testDF[,predictorsNames]),type="prob")
                rocCurve<-roc(response=testDF$gene.mut,predictor=probabilities[,"mut"])
                
                # Update auroc object
                aurocs<-c(aurocs,rocCurve$auc)
                names(aurocs)[length(aurocs)]<-gene
            }
            save(aurocs,file=fname)
        }
    }
}

run()


cd /mnt/d/Dropbox/projects/pancancer/results/025_compareModelTypes
scp -r:/gpfs/work/IscrC_tumornet/pancancer/results/025_compareModelTypes/* .


### And then we plot
library(beeswarm)
library(vioplot)

methods<-sort(c("bayesglm","gbm","lda","rf","svmLinear","svmRadial",
           "mxnet","glm","kknn","pcaNNet","evtree"))
tum<-"BLCA"

## Load results
results<-list()
for(method in methods){
    fname<-paste0("results/025_compareModelTypes/",tum,"-mut_",method,"_aurocs.rda")
    if(file.exists(fname)){
        load(fname)
        results[[method]]<-aurocs        
    }
}


## Plot
png("plots/025_modelViolins.png",w=2000,h=1500,p=45)
plot(1,xlim=c(0,length(results)+1),ylim=c(0.2,1),type="n",xaxt="n",ylab="AUROC",main="Gene Mutation Classification in Bladder Cancer",xlab="")
for(is in 1:length(results)){
    plotme<-unique(results[[is]])
    boxplot(plotme,add=TRUE,at=is,lwd=4,pch=NA,whisklty=0,staplelty=0,width=1.5)
    vioplot(plotme,add=TRUE,at=is,drawRect=FALSE,wex=1.1,border=NA,
            col=rgb(col2rgb("black")[1],col2rgb("black")[2],col2rgb("black")[3],alpha=0.2))
    
    bs<-beeswarm(plotme,pch=20,add=TRUE,at=is,
                 col=rgb(col2rgb("black")[1],col2rgb("black")[2],col2rgb("black")[3],alpha=0.6),
                 spacing=1.5)
}
text(cex=1.5,x=1:length(results),y=0.15,names(results),xpd=TRUE,srt=45,pos=2,offset=0)
abline(h=0.5,lty=2)
abline(v=c(1:length(results)))
dev.off()
```
