## Supplementary Code for "Pan-Cancer and Single-Cell modelling of genomic alterations through gene expression": 026_addNoise_EGFR_GBM.R

```
setwd("D:/Dropbox/projects/pancancer")

### Setup
library(viper)
library(WGCNA)
library(caret)
library(pROC)
library(affy)
library(gplots)
source("../shared/functions/geneids.R")

### Function to model
model<-function(mutrack,varmat,plot=TRUE,method="gbm",main="",colroc="black"){
    # Convert 1/0 into mut/wt categories for binary classification
    mutrack<-as.vector(mutrack)
    mutrack<-ifelse(mutrack==1,"mut","wt")
    # Filter Variables by Variance
    evars<-apply(varmat,1,var)
    varmat<-varmat[evars>0,]
    varmat<-varmat[names(sort(evars,dec=TRUE)[1:min(c(1000,nrow(varmat)))]),]
    # Build data object
    df<-cbind(t(varmat),mutrack)
    df<-as.data.frame(df)
    df$mutrack<-as.factor(df$mutrack)
    # Many caret functions model the probability of the first factor level
    df$mutrack<-relevel(df$mutrack,ref="mut")
    # We want to predict the outcome (the mutations), from a set of predictors (gene expression data), which we code here
    outcomeName<-"mutrack"
    predictorsNames<-colnames(df)[colnames(df)!=outcomeName]
    # Set a seed to use, to make the CV steps reproducible
    set.seed(1)
    # Split the data into training and test sets (0.6, 0.8, try different ones?)
    splitIndex<-createDataPartition(df[,outcomeName],p=0.75,list=FALSE,times=1)
    trainDF<-df[splitIndex,]
    testDF<-df[-splitIndex,]
    # Train Model Parameters (10-fold CV is acceptable)
    objControl<-trainControl(method="repeatedcv",number=10,returnResamp='none',summaryFunction=twoClassSummary,classProbs=TRUE)
    objModel<-train(data.matrix(trainDF[,predictorsNames]),trainDF[,outcomeName], 
                    method=method, 
                    trControl=objControl,  
                    metric="ROC",verbose=FALSE
    )
    # Validating the Model
    # It will predict using the best model: objModel$finalModel
    predictions<-predict(object=objModel,data.matrix(testDF[,predictorsNames]))
    probabilities<-predict(object=objModel,data.matrix(testDF[,predictorsNames]),type="prob")
    rocCurve<-roc(response=testDF$mutrack,predictor=probabilities[,"mut"])
    # Confusion matrix
    confMatrix<-confusionMatrix(predictions,testDF$mutrack)
    # Plot the ROC Curve with AUC
    if(plot){
        plot(rocCurve,col=colroc,cex.lab=2,cex.main=2,print.auc=FALSE,lwd=6,main=main)
        mtext("10-fold CV",side=1)
        grid(col="lightgrey")
        legend(0.3,0.2,legend=paste0("AUC= ", signif(rocCurve$auc,4)))
    }
    # Extract predictors from objModel$finalModel
    if(method=="glmnet"){
        str(objModel$finalModel)
        objModel$finalModel$tuneValue # alpha=1 is lasso
        coefs<-coef(objModel$finalModel,objModel$bestTune$lambda)[,1]
        coefs<-coefs[coefs!=0]
    }
    
    return(rocCurve$auc)
}


################ EGFR amplifications in GBM #################
tum<-"GBM"
gene<-any2entrez("EGFR")

### Take inputs
load(paste0("tums010/",tum,"/",tum,"-tf-regulon.rda"))
load(paste0("tums010/",tum,"/",tum,"-vipermat_tumor.rda"))
colnames(vipermat)<-substr(colnames(vipermat),1,15)
vipermat<-vipermat[intersect(names(regul),rownames(vipermat)),]
load(paste0("tums010/",tum,"/",tum,"-expmat.rda"))
colnames(expmat)<-substr(colnames(expmat),1,15)
load(paste0("tums010/",tum,"/",tum,"-wgcnamat.rda"))
colnames(wgcnamat)<-substr(colnames(wgcnamat),1,15)
load(paste0("tums010/",tum,"/",tum,"-wgcna.rda")) # Clusters
wgcna<-sort(setNames(paste0("w",wgcna),names(wgcna)))
load(paste0("tums010/",tum,"/",tum,"-fcnv.rda"))
mutations<-fcnv
mutations[fcnv>=0.5]<-1
mutations[fcnv<0.5]<-0
mutations<-mutations[!is.na(rownames(mutations)),]
### Keep same-samples (no normals)
commonsamples<-intersect(colnames(mutations),colnames(vipermat))
expmat<-expmat[,commonsamples]
vipermat<-vipermat[,commonsamples]
wgcnamat<-wgcnamat[,commonsamples]
mutations<-mutations[,commonsamples]
mutrack<-mutations[gene,]

### Test Models with expression, viper and wgcna variables (no noise)
fname<-paste0("results/026_",tum,"_",eg2sym(gene),"_amp_aurocs0.rda")
if(!file.exists(fname)){
    aurocs0<-setNames(rep(NA,3),c("exp","viper","wgcna"))
    png(paste0("plots/026_aurocs0_",tum,"_",eg2sym(gene),".png"),w=3000,h=1000,p=40)
    par(mfrow=c(1,3))
    varmat<-expmat
    auc<-model(mutrack=mutrack,varmat=varmat,method="gbm",plot=TRUE,main="Standard Gene Expression",colroc="black")
    aurocs0["exp"]<-auc
    
    varmat<-vipermat
    auc<-model(mutrack=mutrack,varmat=varmat,method="gbm",plot=TRUE,main="VIPER Activity",colroc="firebrick3")
    aurocs0["viper"]<-auc
    
    varmat<-wgcnamat
    auc<-model(mutrack=mutrack,varmat=varmat,method="gbm",plot=TRUE,main="WGCNA clusters",colroc="forestgreen")
    aurocs0["wgcna"]<-auc
    dev.off()
    save(aurocs0,file=fname)
} else {load(fname)}

### Then we shall add noise to the expression matrix
### and we will generate viper and wgcna matrices on the noisy expmat
### until we see a drop in AUROC. Hopefully it will be faster for expmat
noises<-c(0.01,0.1,0.5,1,2,4,6,8,10,12,14,16,18,20,25,30,35,40,50,100)
nperm<-100

# Computation of output!!!
for(noise in rev(noises)){
    for(inp in 1:nperm){
        fname<-paste0("results/026_addNoise/",gene,"_",tum,"_noise",noise,"_aurocs_",inp,".rda")
        if(!file.exists(fname)){
            aurocs<-c()
            message("Doing perm ",inp," of noise ",noise)
            # Add noise (alternative: remove reads from raw counts and then VST)
            set.seed(inp)
            expmat2<-expmat+rnorm(nrow(expmat)*ncol(expmat),sd=noise)
            
            # Generate noise vipermat
            signature<-viperSignature(expmat2,expmat2,method="mean")
            vipermat2<-viper(signature,regul,pleiotropy=FALSE,minsize=10)
            
            # Generate noise wgcnamat
            wgcnamat2<-matrix(NA,nrow=length(unique(wgcna)),ncol=ncol(vipermat))
            colnames(wgcnamat2)<-colnames(vipermat)
            rownames(wgcnamat2)<-unique(wgcna)
            for(iw in unique(wgcna)){
                wgenes<-names(which(wgcna==iw))
                if(length(wgenes)==1){
                    wtukey<-expmat2[wgenes,]
                } else {
                    wsubmat<-expmat2[wgenes,]
                    wtukey<-apply(wsubmat,2,tukey.biweight)
                }
                wgcnamat2[iw,]<-wtukey
            }
            
            # Generate models on these noisy matrices
            varmat<-expmat2
            auc<-model(mutrack=mutrack,varmat=varmat,method="gbm",plot=FALSE)
            aurocs<-auc
            
            varmat<-vipermat2
            auc<-model(mutrack=mutrack,varmat=varmat,method="gbm",plot=FALSE)
            aurocs<-c(aurocs,auc)
            
            varmat<-wgcnamat2
            auc<-model(mutrack=mutrack,varmat=varmat,method="gbm",plot=FALSE)
            aurocs<-c(aurocs,auc)
            
            names(aurocs)<-c("exp","viper","wgcna")
            save(aurocs,file=fname)
        }
    }
}


# Load results
nperm<-100
#noises<-c(0.01,0.1,0.5,1,2,4,6,8,10,12,14,16,18,20,25,30,35,40,50,100)
noises<-c(1,2,4,6,8,10,12,14,16,18,20,25,30,35,40,50,100)
results_wgcna<-matrix(NA,nrow=nperm,ncol=length(noises))
colnames(results_wgcna)<-as.character(noises)
results_exp<-results_viper<-results_wgcna
for(noise in as.character(noises)){
    for(inp in 1:nperm){
        fname<-paste0("results/026_addNoise/",gene,"_",tum,"_noise",noise,"_aurocs_",inp,".rda")
        if(file.exists(fname)){
            load(fname)
            results_exp[inp,noise]<-aurocs["exp"]
            results_viper[inp,noise]<-aurocs["viper"]
            results_wgcna[inp,noise]<-aurocs["wgcna"]
        }
    }
}


# Plot
png(paste0("plots/026_noise_",tum,"_amp_",eg2sym(gene),".png"),w=3000,h=2000,p=60)
par(mar=c(4,4,1,0))
results<-results_exp
means<-c(aurocs0[1],apply(results,2,mean,na.rm=TRUE));names(means)[1]<-"0"
sds<-c(0,apply(results,2,sd,na.rm=TRUE));names(means)[1]<-"0"
plotCI(means,uiw=sds,liw=0,pch=16,type="o",lwd=6,col="black",cex=1.5,xaxt="n",ylab="AUROC",xlab="Gaussian Noise Level",ylim=c(0.4,1.0),gap=0,
       main="EGFR amplifications - classification model in Glioblastoma",add=FALSE)

results<-results_viper
means<-c(aurocs0[2],apply(results,2,mean,na.rm=TRUE));names(means)[1]<-"0"
sds<-c(0,apply(results,2,sd,na.rm=TRUE));names(means)[1]<-"0"
plotCI(means,uiw=sds,liw=0,pch=17,type="o",lwd=6,col="firebrick3",cex=1.5,gap=0,add=TRUE)

results<-results_wgcna
means<-c(aurocs0[3],apply(results,2,mean,na.rm=TRUE));names(means)[1]<-"0"
sds<-c(0,apply(results,2,sd,na.rm=TRUE));names(means)[1]<-"0"
plotCI(means,uiw=sds,liw=0,pch=18,type="o",lwd=6,col="forestgreen",cex=1.5,gap=0,add=TRUE)

grid()
legend("topright",legend=c("Gene Expression","VIPER Activity","WGCNA clusters"),pch=c(16,17,18),pt.cex=2,col=c("black","firebrick3","forestgreen"),bg="white")
axis(1,at=c(1:length(means)),labels=names(means),cex.axis=0.83)

abline(h=0.5)
dev.off()
```
