## Supplementary Code for "Pan-Cancer and Single-Cell modelling of genomic alterations through gene expression": 026b_removeReads_TP53_BRCA.R

```
setwd("D:/Dropbox/projects/pancancer")

### Setup
source("../shared/functions/qol.R")
source("../shared/functions/geneids.R")
library(viper)
library(WGCNA)
library(caret)
library(pROC)
library(affy)
library(gplots)
library(DESeq2)


### Function to model
model<-function(mutrack,varmat,plot=TRUE,method="gbm",main="",colroc="black"){
    # Convert 1/0 into mut/wt categories for binary classification
    mutrack<-as.vector(mutrack)
    mutrack<-ifelse(mutrack==1,"mut","wt")
    # Filter Variables by Variance
    evars<-apply(varmat,1,var)
    varmat<-varmat[evars>0,]
    varmat<-varmat[names(sort(evars,dec=TRUE)[1:min(c(1000,nrow(varmat)))]),]
    # Build data object
    df<-cbind(t(varmat),mutrack)
    df<-as.data.frame(df)
    df$mutrack<-as.factor(df$mutrack)
    # Many caret functions model the probability of the first factor level
    df$mutrack<-relevel(df$mutrack,ref="mut")
    # We want to predict the outcome (the mutations), from a set of predictors (gene expression data), which we code here
    outcomeName<-"mutrack"
    predictorsNames<-colnames(df)[colnames(df)!=outcomeName]
    # Set a seed to use, to make the CV steps reproducible
    set.seed(1)
    # Split the data into training and test sets (0.6, 0.8, try different ones?)
    splitIndex<-createDataPartition(df[,outcomeName],p=0.75,list=FALSE,times=1)
    trainDF<-df[splitIndex,]
    testDF<-df[-splitIndex,]
    # Train Model Parameters (10-fold CV is acceptable)
    objControl<-trainControl(method="repeatedcv",number=10,returnResamp='none',summaryFunction=twoClassSummary,classProbs=TRUE)
    objModel<-train(data.matrix(trainDF[,predictorsNames]),trainDF[,outcomeName], 
                    method=method, 
                    trControl=objControl,  
                    metric="ROC",verbose=FALSE
    )
    # Validating the Model
    # It will predict using the best model: objModel$finalModel
    predictions<-predict(object=objModel,data.matrix(testDF[,predictorsNames]))
    probabilities<-predict(object=objModel,data.matrix(testDF[,predictorsNames]),type="prob")
    rocCurve<-roc(response=testDF$mutrack,predictor=probabilities[,"mut"])
    # Confusion matrix
    confMatrix<-confusionMatrix(predictions,testDF$mutrack)
    # Plot the ROC Curve with AUC
    if(plot){
        plot(rocCurve,col=colroc,cex.lab=2,cex.main=2,print.auc=FALSE,lwd=6,main=main)
        mtext("10-fold CV",side=1)
        grid(col="lightgrey")
        legend(0.3,0.2,legend=paste0("AUC= ", signif(rocCurve$auc,4)))
    }
    # Extract predictors from objModel$finalModel
    if(method=="glmnet"){
        str(objModel$finalModel)
        objModel$finalModel$tuneValue # alpha=1 is lasso
        coefs<-coef(objModel$finalModel,objModel$bestTune$lambda)[,1]
        coefs<-coefs[coefs!=0]
    }
    
    return(rocCurve$auc)
}


################ TP53 in BRCA #################
tum<-"BRCA"
gene<-any2entrez("TP53")

### Take inputs
load(paste0("tums010/",tum,"/",tum,"-tf-regulon.rda"))
load(paste0("tums010/",tum,"/",tum,"-vipermat_tumor.rda"))
colnames(vipermat)<-substr(colnames(vipermat),1,15)
vipermat<-vipermat[intersect(names(regul),rownames(vipermat)),]
load(paste0("tums010/",tum,"/",tum,"-rawcounts.rda"))
colnames(rawcounts)<-substr(colnames(rawcounts),1,15)
load(paste0("tums010/",tum,"/",tum,"-wgcnamat.rda"))
colnames(wgcnamat)<-substr(colnames(wgcnamat),1,15)
load(paste0("tums010/",tum,"/",tum,"-wgcna.rda")) # Clusters
wgcna<-sort(setNames(paste0("w",wgcna),names(wgcna)))
load(paste0("tums010/",tum,"/",tum,"-mutations.rda"))
### Keep same-samples (no normals)
commonsamples<-intersect(colnames(mutations),colnames(vipermat))
rawcounts<-rawcounts[,commonsamples]
vipermat<-vipermat[,commonsamples]
wgcnamat<-wgcnamat[,commonsamples]
mutations<-mutations[,commonsamples]
mutrack<-mutations[gene,]

# Models with original data
load("results/026_BRCA_TP53_aurocs0.rda")

### Then we shall remove reads from the expression matrix
### and we will generate viper and wgcna matrices on the noisy expmat
### until we see a drop in AUROC. Hopefully it will be faster for expmat


### Functions to remove reads
# https://stackoverflow.com/questions/53397861/reduce-total-sum-of-vector-elements-in-r#53397912
### Downsample with Beta Distribution reduction
# \frac{1}{B(\alpha,\beta)} x^{\alpha-1}(1-x)^{\beta-1}\
downsample<-function(x,finalreads=NULL){
    sumx<-sum(x)
    if(finalreads>=sumx){
        return(x)
    }
    rfactor<-finalreads/sumx # reduction factor
    #cfactor<-c(0:2000*rfactor)/1000 # chaos factor (uniform from 0 to 2*rfactor)
    bfactor<-0.1 # Beta distribution factor 
    shape1<-bfactor*rfactor/(1-rfactor)
    cfactor<-rbeta(length(x),shape1,bfactor) # chaos factor (beta distribution centered around rfactor) 
    #plot(density(cfactor));abline(v=rfactor)
    y<-sapply(x,function(xx){
        xx<-round(xx*sample(cfactor,1))
    })
    return(y)
}


# Show how downsampling functions work
if(TRUE){
    #x<-sample(1:1000,10000,replace=TRUE)
    x<-rawcounts[,1]
    set.seed(1)
    png("plots/026b_removeReads_finalreads.png",w=8000,h=6000,p=100)
    par(mfrow=c(3,4))
    for(finalreads in c(5E7,4E7,3E7,2E7,1E7,8E6,5E6,3E6,1E6,5E5,1E5,5E4)){
        y<-downsample(x,finalreads=finalreads)
        plot(log10(x+0.1),log10(y+0.1),pch=20,
             ylab=paste0("Log10 Downsampled gene counts (total: ",sum(y),")"),
             xlab=paste0("Log10 Original gene counts (total: ",sum(x),")"),
             main=paste0("Aimed downsampling: ",signif(finalreads/1E6,3),"M reads"))
        mtext(paste0("Effective downsampling: ",signif(sum(y)/1E6,3),"M reads"),cex=0.8)
        abline(0,1)
    }
    dev.off()
}

# Show beta function distribution on a sample
if(TRUE){
    finalreads<-10E6
    x<-rawcounts[,1]
    set.seed(1)
    sumx<-sum(x)
    if(finalreads>=sumx){
        return(x)
    }
    rfactor<-finalreads/sumx # reduction factor
    bfactor<-0.1 # Beta distribution factor 
    shape1<-bfactor*rfactor/(1-rfactor)
    cfactor<-rbeta(length(x),shape1,bfactor) # chaos factor (beta distribution centered around rfactor) 

    png("plots/026b_removeReads_beta.png",w=2000,h=2000,p=60)
    x1<-signif(sumx/1E6,2)
    x2<-signif(finalreads/1E6,2)
    plot(density(cfactor),lwd=6,main=paste0("Downsampling ",x1,"M to ",x2,"M reads"))
    mtext("Beta function",cex=0.8)
    abline(v=rfactor)
    dev.off()
}

# Computation of output!!!
finalreads<-c(50,40,30,20,15,10,8,5,3,2,1,0.5,0.1,0.05,0.01)
nperm<-10
for(finalread in rev(finalreads)){
    for(inp in 1:nperm){
        fname<-paste0("results/026b_removeReads/",gene,"_",tum,"_betareads_",finalread,"_aurocs_",inp,".rda")
        if(!file.exists(fname)){
            message("Doing perm ",inp," of finalreads ",finalread,"M")
            # Remove reads from raw counts
            set.seed(inp)
            rawcounts2<-apply(rawcounts,2,downsample,finalreads=finalread*1E6)
            
            
            ### VST may crash when there are fewer reads
            # rawcounts3<-rawcounts2[apply(rawcounts2,1,sum)>=100,]+1
            rawcounts3<-rawcounts2
            rawcounts3[rawcounts3==0]<-sample(c(1,2),sum(rawcounts3==0),replace=TRUE)
            conditions<-matrix(rep("TCGA",ncol(rawcounts3)),ncol=1)
            colnames(conditions)<-"Sample"
            rownames(conditions)<-colnames(rawcounts3)
            dds<-DESeqDataSetFromMatrix(countData=rawcounts3,colData=conditions,design=~1)
            vsd<-varianceStabilizingTransformation(dds,blind=TRUE,fitType="mean")
            expmat2<-assay(vsd)
            
            
            
            
            # Generate finalread vipermat
            signature<-viperSignature(expmat2,expmat2,method="mean")
            vipermat2<-viper(signature,regul,pleiotropy=FALSE,minsize=10)
            
            # Generate finalread wgcnamat
            wgcnamat2<-matrix(NA,nrow=length(unique(wgcna)),ncol=ncol(vipermat))
            colnames(wgcnamat2)<-colnames(vipermat)
            rownames(wgcnamat2)<-unique(wgcna)
            for(iw in unique(wgcna)){
                wgenes<-names(which(wgcna==iw))
                if(length(wgenes)==1){
                    wtukey<-expmat2[wgenes,]
                } else {
                    wsubmat<-expmat2[wgenes,]
                    wtukey<-apply(wsubmat,2,tukey.biweight)
                }
                wgcnamat2[iw,]<-wtukey
            }
            
            # Generate models on these reduced matrices
            varmat<-expmat2
            auc<-model(mutrack=mutrack,varmat=varmat,method="gbm",plot=FALSE)
            aurocs<-auc
            
            varmat<-vipermat2
            auc<-model(mutrack=mutrack,varmat=varmat,method="gbm",plot=FALSE)
            aurocs<-c(aurocs,auc)
            
            varmat<-wgcnamat2
            auc<-model(mutrack=mutrack,varmat=varmat,method="gbm",plot=FALSE)
            aurocs<-c(aurocs,auc)
            
            names(aurocs)<-c("exp","viper","wgcna")
            save(aurocs,file=fname)
        }
    }
}


# Load results
nperm<-100
finalreads<-c(50,40,30,20,15,10,8,5,3,2,1,0.5,0.1,0.05,0.01)

results_wgcna<-matrix(NA,nrow=nperm,ncol=length(finalreads))
colnames(results_wgcna)<-as.character(finalreads)
results_exp<-results_viper<-results_wgcna
for(finalread in as.character(finalreads)){
    for(inp in 1:nperm){
        fname<-paste0("results/026b_removeReads/",gene,"_",tum,"_betareads_",finalread,"_aurocs_",inp,".rda")
        if(file.exists(fname)){
            load(fname)
            results_exp[inp,finalread]<-aurocs["exp"]
            results_viper[inp,finalread]<-aurocs["viper"]
            results_wgcna[inp,finalread]<-aurocs["wgcna"]
        }
    }
}


# Plot
png(paste0("plots/026b_removeReads_",tum,"_mut_",eg2sym(gene),".png"),w=3000,h=2000,p=60)
par(mar=c(4,4,1,0))
results<-results_exp
means<-c(aurocs0[1],apply(results,2,mean,na.rm=TRUE));names(means)[1]<-"All"
sds<-c(0,apply(results,2,sd,na.rm=TRUE));names(sds)[1]<-"All"
plotCI(means,uiw=sds,liw=0,pch=16,type="o",lwd=6,col="black",cex=1.5,xaxt="n",ylab="AUROC",xlab="Nr. Reads (M)",ylim=c(0.4,1.0),gap=0,
       main="TP53 mutations - classification model in Breast Cancer",add=FALSE)

results<-results_viper
means<-c(aurocs0[2],apply(results,2,mean,na.rm=TRUE));names(means)[1]<-"All"
sds<-c(0,apply(results,2,sd,na.rm=TRUE));names(sds)[1]<-"All"
plotCI(means,uiw=sds,liw=0,pch=17,type="o",lwd=6,col="firebrick3",cex=1.5,gap=0,add=TRUE)

results<-results_wgcna
means<-c(aurocs0[3],apply(results,2,mean,na.rm=TRUE));names(means)[1]<-"All"
sds<-c(0,apply(results,2,sd,na.rm=TRUE));names(sds)[1]<-"All"
plotCI(means,uiw=sds,liw=0,pch=18,type="o",lwd=6,col="forestgreen",cex=1.5,gap=0,add=TRUE)

grid()
legend("topright",legend=c("Gene Expression","VIPER Activity","WGCNA clusters"),pch=c(16,17,18),pt.cex=2,col=c("black","firebrick3","forestgreen"),bg="white")
axis(1,at=c(1:length(means)),labels=names(means),cex.axis=0.83)

abline(h=0.5)
dev.off()
```
