## Supplementary Code for "Pan-Cancer and Single-Cell modelling of genomic alterations through gene expression": 027_rocNoise.R

```
### Roc Curves of TP53 models with increasing noise (A proof of concept)


### Setup
library(viper)
library(WGCNA)
library(caret)
library(pROC)
library(affy)
library(gplots)
source("../shared/functions/geneids.R")


### Function to model
model<-function(mutrack,varmat,plot=TRUE,method="gbm",main="",colroc="black",returnROC=FALSE){
    # Convert 1/0 into mut/wt categories for binary classification
    mutrack<-as.vector(mutrack)
    mutrack<-ifelse(mutrack==1,"mut","wt")
    # Filter Variables by Variance
    evars<-apply(varmat,1,var)
    varmat<-varmat[evars>0,]
    varmat<-varmat[names(sort(evars,dec=TRUE)[1:min(c(1000,nrow(varmat)))]),]
    # Build data object
    df<-cbind(t(varmat),mutrack)
    df<-as.data.frame(df)
    df$mutrack<-as.factor(df$mutrack)
    # Many caret functions model the probability of the first factor level
    df$mutrack<-relevel(df$mutrack,ref="mut")
    # We want to predict the outcome (the mutations), from a set of predictors (gene expression data), which we code here
    outcomeName<-"mutrack"
    predictorsNames<-colnames(df)[colnames(df)!=outcomeName]
    # Set a seed to use, to make the CV steps reproducible
    set.seed(1)
    # Split the data into training and test sets (0.6, 0.8, try different ones?)
    splitIndex<-createDataPartition(df[,outcomeName],p=0.75,list=FALSE,times=1)
    trainDF<-df[splitIndex,]
    testDF<-df[-splitIndex,]
    # Train Model Parameters (10-fold CV is acceptable)
    objControl<-trainControl(method="repeatedcv",number=10,returnResamp='none',summaryFunction=twoClassSummary,classProbs=TRUE)
    objModel<-train(data.matrix(trainDF[,predictorsNames]),trainDF[,outcomeName], 
                    method=method, 
                    trControl=objControl,  
                    metric="ROC",verbose=FALSE
    )
    # Validating the Model
    # It will predict using the best model: objModel$finalModel
    predictions<-predict(object=objModel,data.matrix(testDF[,predictorsNames]))
    probabilities<-predict(object=objModel,data.matrix(testDF[,predictorsNames]),type="prob")
    rocCurve<-roc(response=testDF$mutrack,predictor=probabilities[,"mut"])
    # Confusion matrix
    confMatrix<-confusionMatrix(predictions,testDF$mutrack)
    # Plot the ROC Curve with AUC
    if(plot){
        plot(rocCurve,col=colroc,cex.lab=2,cex.main=2,print.auc=FALSE,lwd=6,main=main)
        mtext("10-fold CV",side=1)
        grid(col="lightgrey")
        legend(0.3,0.2,legend=paste0("AUC= ", signif(rocCurve$auc,4)))
    }
    # Extract predictors from objModel$finalModel
    if(method=="glmnet"){
        str(objModel$finalModel)
        objModel$finalModel$tuneValue # alpha=1 is lasso
        coefs<-coef(objModel$finalModel,objModel$bestTune$lambda)[,1]
        coefs<-coefs[coefs!=0]
    }
    
    if(returnROC){
        return(rocCurve)
    }
    
    return(rocCurve$auc)
}


################ TP53 in BRCA #################
tum<-"BRCA"
gene<-any2entrez("TP53")

### Take inputs
load(paste0("tums010/",tum,"/",tum,"-tf-regulon.rda"))
load(paste0("tums010/",tum,"/",tum,"-vipermat_tumor.rda"))
colnames(vipermat)<-substr(colnames(vipermat),1,15)
vipermat<-vipermat[intersect(names(regul),rownames(vipermat)),]
load(paste0("tums010/",tum,"/",tum,"-expmat.rda"))
colnames(expmat)<-substr(colnames(expmat),1,15)
load(paste0("tums010/",tum,"/",tum,"-wgcnamat.rda"))
colnames(wgcnamat)<-substr(colnames(wgcnamat),1,15)
load(paste0("tums010/",tum,"/",tum,"-wgcna.rda")) # Clusters
wgcna<-sort(setNames(paste0("w",wgcna),names(wgcna)))
load(paste0("tums010/",tum,"/",tum,"-mutations.rda"))
### Keep same-samples (no normals)
commonsamples<-intersect(colnames(mutations),colnames(vipermat))
expmat<-expmat[,commonsamples]
vipermat<-vipermat[,commonsamples]
wgcnamat<-wgcnamat[,commonsamples]
mutations<-mutations[,commonsamples]
mutrack<-mutations[gene,]

### Test Model with no noise
fname<-"results/027_rocNoise.rda"
if(!file.exists(fname)){
    varmat<-expmat
    roc0<-model(mutrack=mutrack,varmat=varmat,method="gbm",plot=FALSE,returnROC=TRUE)
    noises<-c(0.01,0.1,0.5,1,2,4,6,8,10,12,14,16,18,20,25,30,35,40,50,100)
    rocs<-list()
    for(noise in noises){
        set.seed(1)
        message("Calculating model at noise ",noise)
        varmat<-expmat+rnorm(nrow(expmat)*ncol(expmat),sd=noise)
        roc<-model(mutrack=mutrack,varmat=varmat,method="gbm",plot=FALSE,returnROC=TRUE)
        rocs[[length(rocs)+1]]<-roc
    }
    names(rocs)<-noises
    save(roc0,rocs,file=fname)
} else {
    load(fname)
}

### Plot
#noises<-c(0.01,0.1,0.5,1,2,4,6,8,10,12,14,16,18,20,25,30,35,40,50,100)
noises<-c(1,2,4,6,8,10)
subrocs<-rocs[as.character(noises)]
colroc<-colorpanel(length(subrocs)+1,"red3","orange","navy")
png("plots/027_rocNoise.png",w=2000,h=2000,p=50)
plot(roc0,col=colroc[1],cex.lab=2,cex.main=2,print.auc=FALSE,lwd=10,main="TP53 gbm model with increasing noise")
for(i in 1:length(subrocs)){
    r<-subrocs[[i]]
    lines(r,col=colroc[i],lwd=10)
}
aucs<-signif(c(roc0$auc,sapply(subrocs,function(x){x$auc})),3)
legend("bottomright",lty=1,lwd=10,col=colroc,title="Noise Level (AUC)",cex=1.1,
       legend=paste0(c(0,names(subrocs))," (",aucs,")")
)
dev.off()
```
