## Supplementary Code for "Pan-Cancer and Single-Cell modelling of genomic alterations through gene expression": 029_geneplots.R

```
### Extra plots
# % Mutated vs. AUROC
# Cancer gene census genes vs AUROC vs mutsig-blacklisted


source("../shared/functions/geneids.R")
library(beeswarm)
library(vioplot)

# Setup
if(TRUE){
    raw<-matrix(c(
        "ACC","Adrenocortical Carcinoma",
        "BLCA","Bladder Urothelial Carcinoma",
        "BRCA","Breast Invasive Carcinoma",
        "CESC","Cervical Squamous Cell carcinoma",
        "CHOL","Cholangiocarcinoma",
        "COAD","Colon Adenocarcinoma",
        "DLBC","Diffuse Large B-cell Lymphoma",
        "ESCA","Esophageal Carcinoma",
        "GBM","Glioblastoma Multiforme",
        "HNSC","Head and Neck Squamous Cell Carcinoma",
        "KICH","Kidney Chromophobe",
        "KIRC","Kidney Renal Clear Cell Carcinoma",
        "KIRP","Kidney Renal Papillary Cell Carcinoma",
        "LAML","Acute Myeloid Leukemia",
        "LGG","Brain Lower Grade Glioma",
        "LIHC","Liver Hepatocellular Carcinoma",
        "LUAD","Lung Adenocarcinoma",
        "LUSC","Lung Squamous Cell Carcinoma",
        "MESO","Mesothelioma",
        "OV","Ovarian Serous Cystadenocarcinoma",
        "PAAD","Pancreatic Adenocarcinoma",
        "PCPG","Pheochromocytoma and Paraganglioma",
        "PRAD","Prostate Adenocarcinoma",
        "READ","Rectum Adenocarcinoma",
        "SARC","Sarcoma",
        "SKCM","Skin Cutaneous Melanoma",
        "STAD","Stomach Adenocarcinoma",
        "TGCT","Testicular Germ Cell Tumors",
        "THCA","Thyroid Carcinoma",
        "THYM","Thymoma",
        "UCEC","Uterine Corpus Endometrial Carcinoma",
        "UCS","Uterine Carcinosarcoma",
        "UVM","Uveal Melanoma"
    ),ncol=2,byrow = TRUE
    )
    tums<-setNames(raw[,2],raw[,1])
    rm(raw)
} # Tumor Acronyms
if(TRUE){
    # https://stackoverflow.com/questions/15282580/how-to-generate-a-number-of-most-distinctive-colors-in-r
    library(RColorBrewer)
    brewer.pal.info
    n<-length(tums)
    qual_col_pals<-brewer.pal.info[brewer.pal.info$category == 'qual',]
    cols<-unlist(mapply(brewer.pal, qual_col_pals$maxcolors, rownames(qual_col_pals)))
    set.seed(3)
    tumcols<-setNames(sample(cols, n),names(tums))
    #pie(rep(1,n),col=tumcols)
} # Tumor Colors

###### % Mutated vs. AUROC #######
resultsfile<-"results/029_percmutated.rda"
if(!file.exists(resultsfile)){
    vecmut<-vecamp<-vecdel<-c()
    aucmut<-aucamp<-aucdel<-c()
    for(tum in names(tums)){
        message("Doing ",tum)
        ### Mutations
        fname<-paste0("results/024_aurocs/",tum,"-mut_aurocs.rda")
        if(file.exists(fname)){
            load(fname)
            load(paste0("tums010/",tum,"/",tum,"-mutations.rda"))
            mutations<-mutations[names(aurocs),]
            freqhere<-apply(mutations,1,function(x){sum(x)/length(x)})
            names(freqhere)<-paste0(tum,"_",names(freqhere))
            vecmut<-c(vecmut,freqhere)
            aucmut<-c(aucmut,aurocs)
        }
        ### Amplifications
        fname<-paste0("results/024_aurocs/",tum,"-amp_aurocs.rda")
        if(file.exists(fname)&tum!="LAML"){ # LAML has a single event
            load(fname)
            load(paste0("tums010/",tum,"/",tum,"-fcnv.rda"))
            mutations<-fcnv
            mutations[fcnv>=0.5]<-1
            mutations[fcnv<0.5]<-0
            mutations<-mutations[names(aurocs),]
            freqhere<-apply(mutations,1,function(x){sum(x)/length(x)})
            names(freqhere)<-paste0(tum,"_",names(freqhere))
            vecamp<-c(vecamp,freqhere)
            aucamp<-c(aucamp,aurocs)
        }
        ### Deletions
        fname<-paste0("results/024_aurocs/",tum,"-del_aurocs.rda")
        if(file.exists(fname)&tum!="LAML"){ # LAML has a single event
            load(fname)
            load(paste0("tums010/",tum,"/",tum,"-fcnv.rda"))
            mutations<-fcnv
            mutations[fcnv<=(-0.5)]<-1
            mutations[fcnv>(-0.5)]<-0
            mutations<-mutations[names(aurocs),]
            freqhere<-apply(mutations,1,function(x){sum(x)/length(x)})
            names(freqhere)<-paste0(tum,"_",names(freqhere))
            vecdel<-c(vecdel,freqhere)
            aucdel<-c(aucdel,aurocs)
        }
    }
    save(vecmut,vecamp,vecdel,aucmut,aucamp,aucdel,file=resultsfile)
} else {load(resultsfile)}

# Plot
png("plots/029_percmutated.png",w=6000,h=2000,p=90)
par(mfrow=c(1,3))

x<-vecmut;y<-aucmut
plot(x,y,main="Mutation Models in Pan-Cancer Dataset",xlab="%Mutation in Dataset",ylab="Model AUROC",pch=20,type="n",xlim=c(0,1),ylim=c(0,1))
smoothScatter(x,y,colramp=colorRampPalette(c("white","black")),nrPoints=Inf,add=TRUE,pch=20,nbin=512,col="black")
smoothingSpline<-smooth.spline(x,y,spar=0.9)
lines(smoothingSpline,lwd=6,col="black")
pcc<-cor.test(x,y)
mtext(paste0("PCC=",signif(pcc$estimate,3)),cex=0.8)

x<-vecamp;y<-aucamp
plot(x,y,main="Amplification Models in Pan-Cancer Dataset",xlab="%Amplification in Dataset",ylab="Model AUROC",pch=20,type="n",xlim=c(0,1),ylim=c(0,1))
smoothScatter(x,y,colramp=colorRampPalette(c("white","red")),nrPoints=Inf,add=TRUE,pch=20,nbin=512,col="red3")
smoothingSpline<-smooth.spline(x,y,spar=0.9)
lines(smoothingSpline,lwd=6,col="red3")
pcc<-cor.test(x,y)
mtext(paste0("PCC=",signif(pcc$estimate,3)),cex=0.8)

x<-vecdel;y<-aucdel
plot(x,y,main="Deletion Models in Pan-Cancer Dataset",xlab="%Deletion in Dataset",ylab="Model AUROC",pch=20,type="n",xlim=c(0,1),ylim=c(0,1))
smoothScatter(x,y,colramp=colorRampPalette(c("white","blue")),nrPoints=Inf,add=TRUE,pch=20,nbin=512,col="navy")
smoothingSpline<-smooth.spline(x,y,spar=0.9)
lines(smoothingSpline,lwd=6,col="navy")
pcc<-cor.test(x,y)
mtext(paste0("PCC=",signif(pcc$estimate,3)),cex=0.8)

dev.off()


############################################################################
################# Behavior of Cancer Gene Census Genes #####################
### Load Cancer Gene Census
load("../shared/genelists/CancerGeneCensus/cgc_genelists.rda")
if(TRUE){ # Remove genes that are both TSG and oncogenes
    tmp<-setdiff(oncogenes,tsg)
    tsg<-setdiff(tsg,oncogenes)
    oncogenes<-tmp
    rm(tmp)
}


### Load MutSig blackllisted
mutsig<-any2entrez(read.delim("../shared/genelists/mutSig_blacklist.txt",as.is=TRUE,header=FALSE)[,1])


### Here we will compare genes with each other in separated pan-cancer contexts, so we will calculate ranks
resultsfile<-"results/029_rankmutated.rda"
if(!file.exists(resultsfile)){
    rankmut<-rankamp<-rankdel<-c()
    for(tum in names(tums)){
        message("Doing ",tum)
        ### Mutations
        fname<-paste0("results/024_aurocs/",tum,"-mut_aurocs.rda")
        if(file.exists(fname)){
            load(fname)
            out<-rank(aurocs)/length(aurocs)
            names(out)<-paste0(tum,"_",names(out))
            rankmut<-c(rankmut,out)
        }
        ### Amplifications
        fname<-paste0("results/024_aurocs/",tum,"-amp_aurocs.rda")
        if(file.exists(fname)&tum!="LAML"){ # LAML has a single event
            load(fname)
            out<-rank(aurocs)/length(aurocs)
            names(out)<-paste0(tum,"_",names(out))
            rankamp<-c(rankamp,out)
        }
        ### Deletions
        fname<-paste0("results/024_aurocs/",tum,"-del_aurocs.rda")
        if(file.exists(fname)&tum!="LAML"){ # LAML has a single event
            load(fname)
            out<-rank(aurocs)/length(aurocs)
            names(out)<-paste0(tum,"_",names(out))
            rankdel<-c(rankdel,out)
        }
    }
    save(rankmut,rankamp,rankdel,file=resultsfile)
} else {load(resultsfile)}


######### Plot
png("plots/029_CGC.png",w=2500,h=1000,p=45)
par(mar=c(10,4,3,1),mfrow=c(1,3))

### Muts
x<-rankmut
g_allcgc<-x[gsub(".+_","",names(x))%in%allcgc]
g_tsg<-x[gsub(".+_","",names(x))%in%tsg]
g_oncogenes<-x[gsub(".+_","",names(x))%in%oncogenes]
g_mutsig<-x[gsub(".+_","",names(x))%in%mutsig]
g_others<-x[!gsub(".+_","",names(x))%in%allcgc
            & !gsub(".+_","",names(x))%in%tsg
            & !gsub(".+_","",names(x))%in%oncogenes
            & !gsub(".+_","",names(x))%in%mutsig
            ]
results<-list(g_others,g_mutsig,g_oncogenes,g_tsg)
names(results)<-c("Other Genes","MutSig Blacklist","Oncogenes","Tumor Suppressors")
plot(1,xlim=c(0,length(results)+1),ylim=c(0,1),type="n",xaxt="n",ylab="Model Rank",main="Model Performance in Gene Types",xlab="")
for(is in 1:length(results)){
    plotme<-unique(results[[is]])
    boxplot(plotme,add=TRUE,at=is,lwd=4,pch=NA,whisklty=0,staplelty=0,width=1.5)
    vioplot(plotme,add=TRUE,at=is,drawRect=FALSE,wex=0.8,border=NA,
            col=rgb(col2rgb("black")[1]/256,col2rgb("black")[2]/256,col2rgb("black")[3]/256,alpha=0.2))
    
    bs<-beeswarm(plotme,pch=20,cex=0.5,add=TRUE,at=is,
                 col=rgb(col2rgb("black")[1]/256,col2rgb("black")[2]/256,col2rgb("black")[3]/256,alpha=0.6),
                 spacing=1.5)
}
text(cex=1.5,x=1:length(results),y=-0.05,names(results),xpd=TRUE,srt=45,pos=2,offset=0)
abline(v=c(1:length(results)))
mtext("Mutated Genes",cex=0.8,font=2)
# Significant Differences
wilcox.test(g_others,g_oncogenes,alternative="greater") # 0.04957
#text(x=3,y=-0.01,"*",cex=3)


### Amps
x<-rankamp
g_allcgc<-x[gsub(".+_","",names(x))%in%allcgc]
g_tsg<-x[gsub(".+_","",names(x))%in%tsg]
g_oncogenes<-x[gsub(".+_","",names(x))%in%oncogenes]
g_mutsig<-x[gsub(".+_","",names(x))%in%mutsig]
g_others<-x[!gsub(".+_","",names(x))%in%allcgc
            & !gsub(".+_","",names(x))%in%tsg
            & !gsub(".+_","",names(x))%in%oncogenes
            & !gsub(".+_","",names(x))%in%mutsig
            ]
results<-list(g_others,g_oncogenes,g_tsg)
names(results)<-c("Other Genes","Oncogenes","Tumor Suppressors") # MutSig removed: no MutSig genes amplified or deleted in enough smaples to calculate model
plot(1,xlim=c(0,length(results)+1),ylim=c(0,1),type="n",xaxt="n",ylab="Model Rank",main="Model Performance in Gene Types",xlab="")
for(is in 1:length(results)){
    plotme<-unique(results[[is]])
    boxplot(plotme,add=TRUE,at=is,lwd=4,pch=NA,whisklty=0,staplelty=0,width=1.5)
    vioplot(plotme,add=TRUE,at=is,drawRect=FALSE,wex=0.8,border=NA,
            col=rgb(col2rgb("red3")[1]/256,col2rgb("red3")[2]/256,col2rgb("red3")[3]/256,alpha=0.2))
    
    bs<-beeswarm(plotme,pch=20,cex=0.5,add=TRUE,at=is,
                 col=rgb(col2rgb("red3")[1]/256,col2rgb("red3")[2]/256,col2rgb("red3")[3]/256,alpha=0.6),
                 spacing=1.5)
}
text(cex=1.5,x=1:length(results),y=-0.05,names(results),xpd=TRUE,srt=45,pos=2,offset=0)
abline(v=c(1:length(results)))
mtext("Amplified Genes",cex=0.8,font=2)


### Dels
x<-rankdel
g_allcgc<-x[gsub(".+_","",names(x))%in%allcgc]
g_tsg<-x[gsub(".+_","",names(x))%in%tsg]
g_oncogenes<-x[gsub(".+_","",names(x))%in%oncogenes]
g_mutsig<-x[gsub(".+_","",names(x))%in%mutsig]
g_others<-x[!gsub(".+_","",names(x))%in%allcgc
            & !gsub(".+_","",names(x))%in%tsg
            & !gsub(".+_","",names(x))%in%oncogenes
            & !gsub(".+_","",names(x))%in%mutsig
            ]
results<-list(g_others,g_oncogenes,g_tsg)
names(results)<-c("Other Genes","Oncogenes","Tumor Suppressors") # MutSig removed: no MutSig genes amplified or deleted in enough smaples to calculate model
plot(1,xlim=c(0,length(results)+1),ylim=c(0,1),type="n",xaxt="n",ylab="Model Rank",main="Model Performance in Gene Types",xlab="")
for(is in 1:length(results)){
    plotme<-unique(results[[is]])
    boxplot(plotme,add=TRUE,at=is,lwd=4,pch=NA,whisklty=0,staplelty=0,width=1.5)
    vioplot(plotme,add=TRUE,at=is,drawRect=FALSE,wex=0.8,border=NA,
            col=rgb(col2rgb("navy")[1]/256,col2rgb("navy")[2]/256,col2rgb("navy")[3]/256,alpha=0.2))
    
    bs<-beeswarm(plotme,pch=20,cex=0.5,add=TRUE,at=is,
                 col=rgb(col2rgb("navy")[1]/256,col2rgb("navy")[2]/256,col2rgb("navy")[3]/256,alpha=0.6),
                 spacing=1.5)
}
text(cex=1.5,x=1:length(results),y=-0.05,names(results),xpd=TRUE,srt=45,pos=2,offset=0)
abline(v=c(1:length(results)))
mtext("Deleted Genes",cex=0.8,font=2)
# Significant Differences
wilcox.test(g_others,g_oncogenes,alternative="greater") # 0.003705
text(x=2,y=-0.01,"**",cex=3)
wilcox.test(g_others,g_tsg,alternative="less") # 0.0005019
text(x=3,y=1.01,"**",cex=3)


dev.off()
```
