## Supplementary Code for "Pan-Cancer and Single-Cell modelling of genomic alterations through gene expression": 040_cropseq.R

```
### We will prove that Gene Networks are better than expression at prediction mutations


# ARACNe networks based on CCLE
load("../shared/ccle/ccle-cotf-regulon.rda")
load("../shared/ccle/ccle-tf-regulon.rda")


### Dataset https://www.ncbi.nlm.nih.gov/geo/query/acc.cgi?acc=GSE92872
### Paper CROPSeq Datligner et al., 2017 Nature Methods
### Jurkat cells
fname<-"cropseq/scdataset.rda"
if(!file.exists(fname)){
    scrawcounts<-read.csv(gzfile("cropseq/CROP-seq_Jurkat_TCR.digital_expression.500genes.only_assigned.csv.gz"),skip=6,row.names=1,as.is=TRUE,header=FALSE)
    dim(scrawcounts) # 36722  5905
    scgrnas<-read.csv(gzfile("cropseq/CROP-seq_Jurkat_TCR.digital_expression.500genes.only_assigned.csv.gz"),skip=3,nrow=1,header=FALSE,as.is=TRUE)
    scgenes<-read.csv(gzfile("cropseq/CROP-seq_Jurkat_TCR.digital_expression.500genes.only_assigned.csv.gz"),skip=4,nrow=1,header=FALSE,as.is=TRUE)
    scgrnas<-as.character(scgrnas[2:length(scgrnas)])
    scgenes<-as.character(scgenes[2:length(scgenes)])
    scsamples<-paste0(scgenes,"_",1:ncol(scrawcounts))
    colnames(scrawcounts)<-scsamples
    screadcounts<-apply(scrawcounts,2,sum)
    dim(scrawcounts) # 36722  5905
    save(scrawcounts,scgenes,scgrnas,screadcounts,file=fname)
    ### RPMs (VST is not designed for many zeroes)
    scrpm<-apply(scrawcounts,2,function(x){1E6*x/sum(x)})
    save(scrawcounts,scrpm,scgenes,scgrnas,screadcounts,file=fname)
} else {load(fname)}
rm(scrpm)


### Seurat pipeline (regress out cell cycle)
source("../shared/functions/geneids.R")
source("../shared/functions/qol.R")
library(Seurat)
library(Rtsne)

## Create Seurat object
# We will keep all genes expressed in >=3 cells and all cells with at least 200 detected genes:
seuset <- CreateSeuratObject(
    raw.data = scrawcounts,
    min.cells = 3, 
    min.genes = 200
)

## Normalize with Seurat
# By default, Seurat employs a global-scaling normalization method LogNormalize that
# normalizes the gene expression measurements for each cell by the total expression,
# multiplies this by a scale factor (10,000 by default), and log-transforms the result:
seuset <- NormalizeData(
    object = seuset, 
    normalization.method = "LogNormalize", 
    scale.factor = 10000
)

# Mean Variability plot for genes ?seurat
expmat<-as.matrix(seuset@data)
avgexp<-apply(expmat,1,ExpMean)
dispersion<-apply(expmat,1,LogVMR) # variance of mean ratio

# Get Highly variable genes for clustering
seuset<-FindVariableGenes(seuset,mean.function=ExpMean,dispersion.function=LogVMR,do.plot=FALSE)
hvargenes<-seuset@var.genes

## Regress vs UMI counts
seuset <- ScaleData(
    object = seuset, 
    vars.to.regress = c("nUMI")
)
expmat<-as.matrix(seuset@scale.data)

## Regress vs Cell Cycle markers
# Cell cycle markers, from Tirosh et al, 2015
ccgenes <- readLines(con = "cropseq/regev_lab_cell_cycle_genes.txt")
ccgenes<-eg2sym(sym2eg(ccgenes))
setdiff(ccgenes,rownames(rawcounts))

# We can segregate this list into markers of G2/M phase and markers of S phase
s.genes <- ccgenes[1:43]
g2m.genes <- ccgenes[44:97]
s.genes<-intersect(s.genes,rownames(expmat))
g2m.genes<-intersect(g2m.genes,rownames(expmat))

## Assign Cell-Cycle Scores
seuset<-CellCycleScoring(seuset,s.genes=s.genes,g2m.genes=g2m.genes,set.ident=TRUE)
# View cell cycle scores and phase assignments
head(seuset@meta.data)

# TSNE on cell cycle markers
tsnemat<-expmat[c(s.genes,g2m.genes),]
fname<-paste0("results/040_tsne_cellCycle_Datlinger.rda")
if(!file.exists(fname)){ttt<-Rtsne(t(tsnemat),max_iter=1000);save(ttt,file=fname)}else{load(fname)}
x<-ttt$Y[,1]
y<-ttt$Y[,2]

phases<-as.character(seuset@meta.data$Phase)
mycols<-phases
mycols[phases=="G1"]<-"salmon"
mycols[phases=="G2M"]<-"seagreen"
mycols[phases=="S"]<-"cornflowerblue"

# Plot
png(paste0("plots/040_tsne_cellCycle_Datlinger.png"),w=1500,h=1000,p=30)
plot(x,y,pch=20,xlab="TSNE1",ylab="TSNE2",main=paste0("CROP-Seq cells"),col=mycols,xlim=c(min(x),max(x)*1.5))
mtext(paste0(length(x)," cells"),cex=0.8,font=2)
grid()
legend("bottomright",pch=19,col=c("salmon","seagreen","cornflowerblue"),legend=c("G1","G2M","S"))
dev.off()

### Regress out cell cycle scores during data scaling
# We now attempt to subtract (‘regress out’) this source of heterogeneity from the data.
seuset<-ScaleData(seuset,vars.to.regress=c("S.Score","G2M.Score"))
expmat<-as.matrix(seuset@scale.data)

# TSNE on cell cycle markers
tsnemat<-expmat[c(s.genes,g2m.genes),]
fname<-paste0("results/040_tsne_regout_Datlinger.rda")
if(!file.exists(fname)){ttt<-Rtsne(t(tsnemat),max_iter=1000);save(ttt,file=fname)}else{load(fname)}
x<-ttt$Y[,1]
y<-ttt$Y[,2]

# Plot
png(paste0("plots/040_tsne_cellCycleRemoved_Datlinger.png"),w=1500,h=1000,p=30)
plot(x,y,pch=20,xlab="TSNE1",ylab="TSNE2",main=paste0("CROP-Seq cells"),col=mycols,xlim=c(min(x),max(x)*1.5))
mtext(paste0(length(x)," cells"),cex=0.8,font=2)
grid()
legend("bottomright",pch=19,col=c("salmon","seagreen","cornflowerblue"),legend=c("G1","G2M","S"))
dev.off()

## Clustering based on Cas9 and RUNX1T1
scexpmat<-as.matrix(seuset@scale.data)
save(scexpmat,scgenes,scgrnas,file="cropseq/scexpmat.rda")
```
