## Supplementary Code for "Pan-Cancer and Single-Cell modelling of genomic alterations through gene expression": 041_scnetworks.R

```
library(viper)
library(caret)
library(Rtsne)


setwd("D:/Dropbox/projects/pancancer/")

# ARACNe networks based on CCLE
# load("../shared/ccle/ccle-cotf-regulon.rda")
load("../shared/ccle/ccle-tf-regulon.rda")

# Datlinger dataset
load("cropseq/scexpmat.rda") # scexpmat scgenes scgrnas

# Calculate VIPER mat
fname<-"cropseq/scvipermat.rda"
if(!file.exists(fname)){
    scvipermat<-viper(scexpmat,regul=regul)
    save(scvipermat,file=fname)
} else {load(fname)}

dim(scexpmat) # 26623  5905
dim(scvipermat) # 1115 5905

# barplot
x<-sort(table(scgenes))
png("plots/041_scgenes.png",w=2000,h=1500,p=55)
par(las=2)
barplot(x,main="Datlinger CROP-Seq dataset, mutations")
grid()
dev.off()


# TSNE
fname<-"results/sctsne.rda"
if(!file.exists(fname)){
    ttte<-Rtsne(t(scexpmat),max_iter=1000)
    tttv<-Rtsne(t(scvipermat),max_iter=1000)
    save(ttte,tttv,file=fname)
} else {load(fname)}


png("plots/041_tsne_viper.png",h=11000,w=3000,pointsize=60)
x<-setNames(tttv$Y[,1],colnames(scvipermat))
y<-setNames(tttv$Y[,2],colnames(scvipermat))
par(mfrow=c(11,3))
for(scgene in unique(scgenes)){
    plot(x[which(scgenes=="CTRL")],main=scgene,y[which(scgenes=="CTRL")],xlab="TSNE1",ylab="TSNE2",pch=20,col="grey")
    if(scgene!="CTRL"){
        points(x[which(scgenes==scgene)],y[which(scgenes==scgene)],col="black",pch="x")
    }
    mtext("Viper Activity",cex=0.8)
}
dev.off()

png("plots/041_tsne_expression.png",h=11000,w=3000,pointsize=60)
x<-setNames(ttte$Y[,1],colnames(scexpmat))
y<-setNames(ttte$Y[,2],colnames(scexpmat))
par(mfrow=c(11,3))
for(scgene in unique(scgenes)){
    plot(x[which(scgenes=="CTRL")],main=scgene,y[which(scgenes=="CTRL")],xlab="TSNE1",ylab="TSNE2",pch=20,col="grey")
    if(scgene!="CTRL"){
        points(x[which(scgenes==scgene)],y[which(scgenes==scgene)],col="black",pch="x")
    }
    mtext("Gene Expression",cex=0.8)
}
dev.off()


###### Build Models

### Function to generate the models
probmodel<-function(testgene,varmat_annotation,ctrlname,varmat,method="gbm"){
    # Define what is mutated and what is ctrl
    mutrack<-varmat_annotation
    mutrack[varmat_annotation==testgene]<-1
    mutrack[varmat_annotation==ctrlname]<-0
    mutrack[!mutrack%in%c("1","0")]<-NA
    mutrack<-as.numeric(mutrack)
    # Convert 1/0 into mut/wt categories for binary classification
    mutrack<-ifelse(mutrack==1,"mut","wt")
    # Subset
    varmat<-varmat[,!is.na(mutrack)]
    mutrack<-mutrack[!is.na(mutrack)]
    
    # Filter Variables by Variance
    evars<-apply(varmat,1,var)
    varmat<-varmat[evars>0,]
    varmat<-varmat[names(sort(evars,dec=TRUE)[1:min(c(100,nrow(varmat)))]),]
    # Build data object
    df<-cbind(t(varmat),mutrack)
    df<-as.data.frame(df)
    df$mutrack<-as.factor(df$mutrack)
    # Many caret functions model the probability of the first factor level
    df$mutrack<-relevel(df$mutrack,ref="mut")
    # We want to predict the outcome (the mutations), from a set of predictors (gene expression data), which we code here
    outcomeName<-"mutrack"
    predictorsNames<-colnames(df)[colnames(df)!=outcomeName]
    # Split the data into training and test sets (0.6, 0.8, try different ones?)
    splitIndex<-createDataPartition(df[,outcomeName],p=0.75,list=FALSE,times=1)
    trainDF<-df[splitIndex,]
    testDF<-df[-splitIndex,]
    submutrack<-mutrack[-splitIndex]
    # Train Model Parameters (10-fold CV is acceptable)
    objControl<-trainControl(method="repeatedcv",number=10,returnResamp='none',summaryFunction=twoClassSummary,classProbs=TRUE)
    objModel<-train(data.matrix(trainDF[,predictorsNames]),trainDF[,outcomeName], 
                    method=method, 
                    trControl=objControl,  
                    metric="ROC",verbose=FALSE
    )
    predictions<-predict(object=objModel,testDF)
    probabilities<-predict(object=objModel,testDF,type="prob")
    output<-cbind(probabilities,submutrack)
    return(output)
}


### Each gene, multiple training/test splits
testgenes<-setdiff(unique(scgenes),"CTRL")
ncontrol<-table(scgenes)["CTRL"]
nsplit<-10
inputmat<-scexpmat
vinputmat<-scvipermat
for(testgene in testgenes){
    set.seed(1)
    png(paste0("plots/041_kotests/041_",testgene,".png"),w=2000,h=1500,pointsize=40)
    ntest<-table(scgenes)[testgene]
    par(mfrow=c(1,2))
    
    ### Test expression model on single cell data
    probs<-matrix(nrow=0,ncol=3)
    colnames(probs)<-c("mut","wt","submutrack")
    for (ns in 1:nsplit){
        message("Doing ",testgene," expression split ",ns)
        newprobs<-probmodel(testgene,scgenes,"CTRL",inputmat)
        probs<-rbind(probs,newprobs)
    }
    prob_ctrl<-100*probs[probs[,3]=="wt","mut"]
    prob_test<-100*probs[probs[,3]=="mut","mut"]
    boxplot(prob_ctrl,prob_test,cex.axis=0.8,names=c(paste0("CTRL (",ncontrol,")"),paste0("KO (",ntest,")")),pch=NA,col="mediumseagreen",main="Gene Expression",ylab="KO predicted Probability (%)")
    stripchart(list(prob_ctrl,prob_test),jitter=0.1,vertical=TRUE,method="jitter",pch=20,col=c("darkgreen","black"),bg="darkgreen",add=TRUE)
    wt<-wilcox.test(prob_ctrl,prob_test,alt="less")
    mtext(paste0(testgene," model, p=",signif(wt$p.value,4)),cex=0.8)
    
    ### Test viper model on single cell data
    probs<-matrix(nrow=0,ncol=3)
    colnames(probs)<-c("mut","wt","submutrack")
    for (ns in 1:nsplit){
        message("Doing ",testgene," viper split ",ns)
        newprobs<-probmodel(testgene,scgenes,"CTRL",vinputmat)
        probs<-rbind(probs,newprobs)
    }
    prob_ctrl<-100*probs[probs[,3]=="wt","mut"]
    prob_test<-100*probs[probs[,3]=="mut","mut"]
    boxplot(prob_ctrl,prob_test,cex.axis=0.8,names=c(paste0("CTRL (",ncontrol,")"),paste0("KO (",ntest,")")),pch=NA,col="cornflowerblue",main="VIPER Activity")
    stripchart(list(prob_ctrl,prob_test),jitter=0.1,vertical=TRUE,method="jitter",pch=20,col=c("navy","black"),bg="darkgreen",add=TRUE)
    wt<-wilcox.test(prob_ctrl,prob_test,alt="less")
    mtext(paste0(testgene," model, p=",signif(wt$p.value,4)),cex=0.8)
    dev.off()
}
```
