## Supplementary Code for "Pan-Cancer and Single-Cell modelling of genomic alterations through gene expression": 042_shifrut.R

```
source("../shared/functions/geneids.R")

### Shifrut Dataset
# https://www.ncbi.nlm.nih.gov/geo/query/acc.cgi?acc=GSE119450
# GSM3375483	D1_NoStim_10x
# GSM3375484	D1_Stim_10x
# GSM3375485	D2_NoStim_10x
# GSM3375486	D2_Stim_10x
# GSM3375487	D1_NoStim_ReAmp
# GSM3375488	D1_Stim_ReAmp
# GSM3375489	D2_NoStim_ReAmp
# GSM3375490	D2_Stim_ReAmp
# Pooled CRISPR screen with single cell RNASeq readout (Perturb-Seq / CROP-Seq) in primary human T cells.
# Dataset includes CD8 T cells from two donors, for two conditions: with TCR stimulation or No stimulation.

### Format the two unstimulated datasets

if(FALSE){
    # Loading 10x matrices
    # https://support.10xgenomics.com/single-cell-gene-expression/software/pipelines/latest/output/matrices
    library(Matrix)
    library(Seurat)
    
    ### D1N expression
    matrix_dir<-"shifrut/GSM3375483_D1N_matrix/"
    barcode.path<-paste0(matrix_dir,"barcodes.tsv.gz")
    genes.path<-paste0(matrix_dir,"genes.tsv.gz")
    matrix.path<-paste0(matrix_dir,"matrix.mtx.gz")
    rawcounts1<-readMM(file=matrix.path)
    gene.names<-read.delim(genes.path,header=FALSE,stringsAsFactors=FALSE)
    barcode.names<-read.delim(barcode.path,header=FALSE,stringsAsFactors=FALSE)
    colnames(rawcounts1)<-barcode.names$V1
    rownames(rawcounts1)<-gene.names$V1
    # D1N annotation
    anno1<-read.csv("shifrut/GSM3375487_D1N_CellBC_sgRNA.csv",as.is=TRUE)
    #intersect(gsub("-1","",colnames(rawcounts1)),anno1[,1])
    png("plots/042_grnas_D1N.png",w=1000,h=800,p=30)
    par(las=2)
    barplot(sort(table(gsub(".+\\.","",anno1[,2]))),ylab="nr. cells",main="D1N")
    dev.off()
    
    ### D2N expression
    matrix_dir<-"shifrut/GSM3375485_D2N_matrix/"
    barcode.path<-paste0(matrix_dir,"barcodes.tsv.gz")
    genes.path<-paste0(matrix_dir,"genes.tsv.gz")
    matrix.path<-paste0(matrix_dir,"matrix.mtx.gz")
    rawcounts2<-readMM(file=matrix.path)
    gene.names<-read.delim(genes.path,header=FALSE,stringsAsFactors=FALSE)
    barcode.names<-read.delim(barcode.path,header=FALSE,stringsAsFactors=FALSE)
    colnames(rawcounts2)<-barcode.names$V1
    rownames(rawcounts2)<-gene.names$V1
    # D2N annotation
    anno2<-read.csv("shifrut/GSM3375489_D2N_CellBC_sgRNA.csv",as.is=TRUE)
    #intersect(gsub("-1","",colnames(rawcounts2)),anno2[,1])
    png("plots/042_grnas_D2N.png",w=1000,h=800,p=30)
    par(las=2)
    barplot(sort(table(gsub(".+\\.","",anno2[,2]))),ylab="nr. cells",main="D2N")
    dev.off()
    
    # Guides in common of both expriments
    intersect(gsub(".+\\.","",anno1[,2]),gsub(".+\\.","",anno2[,2]))
    
    ### Seurat normalization
    rawcounts<-rawcounts1
    seuset<-CreateSeuratObject(raw.data=rawcounts,min.cells=3,min.genes=200)
    seuset<-NormalizeData(object=seuset,normalization.method="LogNormalize",scale.factor=10000)
    seuset<-FindVariableGenes(seuset,mean.function=ExpMean,dispersion.function=LogVMR,do.plot=FALSE)
    # Regress vs. UMI counts
    seuset<-ScaleData(object=seuset,vars.to.regress=c("nUMI")) 
    # Regress vs. cell cycle
    ccgenes<-readLines(con="cropseq/regev_lab_cell_cycle_genes.txt")
    ccgenes<-eg2ens(sym2eg(ccgenes))
    s.genes<-ccgenes[1:43]
    g2m.genes<-ccgenes[44:97]
    s.genes<-intersect(s.genes,rownames(rawcounts))
    g2m.genes<-intersect(g2m.genes,rownames(rawcounts))
    seuset<-CellCycleScoring(seuset,s.genes=s.genes,g2m.genes=g2m.genes,set.ident=TRUE)
    seuset<-ScaleData(seuset,vars.to.regress=c("S.Score","G2M.Score"))
    scexpmat1<-as.matrix(seuset@scale.data)
    save(scexpmat1,anno1,file="shifrut/scexpmat_D1N.rda")

    rawcounts<-rawcounts2
    seuset<-CreateSeuratObject(raw.data=rawcounts,min.cells=3,min.genes=200)
    seuset<-NormalizeData(object=seuset,normalization.method="LogNormalize",scale.factor=10000)
    seuset<-FindVariableGenes(seuset,mean.function=ExpMean,dispersion.function=LogVMR,do.plot=FALSE)
    # Regress vs. UMI counts
    seuset<-ScaleData(object=seuset,vars.to.regress=c("nUMI")) 
    # Regress vs. cell cycle
    ccgenes<-readLines(con="cropseq/regev_lab_cell_cycle_genes.txt")
    ccgenes<-eg2ens(sym2eg(ccgenes))
    s.genes<-ccgenes[1:43]
    g2m.genes<-ccgenes[44:97]
    s.genes<-intersect(s.genes,rownames(rawcounts))
    g2m.genes<-intersect(g2m.genes,rownames(rawcounts))
    seuset<-CellCycleScoring(seuset,s.genes=s.genes,g2m.genes=g2m.genes,set.ident=TRUE)
    seuset<-ScaleData(seuset,vars.to.regress=c("S.Score","G2M.Score"))
    scexpmat2<-as.matrix(seuset@scale.data)
    save(scexpmat2,anno2,file="shifrut/scexpmat_D2N.rda")
}


### VIPER-transform those matrices (beware of ENSG)
library(viper)

# CCLE ARACNe network
fname<-"../shared/ccle/ccle-tf-regulon_ensg.rda"
if(!file.exists(fname)){
    load("../shared/ccle/ccle-tf-regulon.rda")
    names(regul)<-eg2ens(sym2eg(names(regul)))
    regul<-lapply(regul,function(x){
        tfmod<-x$tfmode
        names(tfmod)<-eg2ens(sym2eg(names(x$tfmode)))
        return(list(tfmode=tfmod,likelihood=x$likelihood))
    })
    save(regul,file=fname)
} else {load(fname)}


load("shifrut/scexpmat_D1N.rda")
scvipermat1<-viper(scexpmat1,regul)
save(scvipermat1,file="shifrut/scvipermat1_D1N.rda")

load("shifrut/scexpmat_D2N.rda")
scvipermat2<-viper(scexpmat2,regul)
save(scvipermat2,file="shifrut/scvipermat2_D2N.rda")
```
