## Supplementary Code for "Pan-Cancer and Single-Cell modelling of genomic alterations through gene expression": 043c_datlinger_models.R

```
setwd("D:/Dropbox/projects/pancancer")

library(caret)
library(pROC)

# Datlinger dataset
load("cropseq/scexpmat.rda") # scexpmat scgenes scgrnas

# VIPER matrix calculated on CCLE network
load("cropseq/scvipermat.rda")
scvipermat<-scvipermat[!is.na(rownames(scvipermat)),]
anno<-setNames(scgenes,colnames(scexpmat))

# Submatrices
dim(scexpmat) # 26623  5905
rowvars<-apply(scexpmat,1,var)
subexpmat<-scexpmat[names(sort(rowvars,dec=TRUE)[1:1000]),]
dim(scvipermat) # 1115 5905
rowvars<-apply(scvipermat,1,var)
subvipermat<-scvipermat[names(sort(rowvars,dec=TRUE)[1:1000]),]


# Pick genes
obs<-sort(table(anno),dec=TRUE)
igenes<-names(obs)[obs>=10]
ctrlgenes<-grep("^CTRL",igenes,value=TRUE)
igenes<-grep("^CTRL",igenes,value=TRUE,invert=TRUE)


#######################################################
### Hic sunt models
#######################################################
if(FALSE){
    method="gbm"
    source("code/scmodel.R")
    eaurocs<-vaurocs<-list()
    for(i in 1:1000){
        varmat<-subexpmat
        aurocs<-c()
        for(igene in igenes){
            message("Doing ",igene)
            scmodel()
        }
        names(aurocs)<-igenes
        eaurocs[[i]]<-aurocs
        
        varmat<-subvipermat
        aurocs<-c()
        for(igene in igenes){
            message("Doing ",igene)
            scmodel()
        }
        names(aurocs)<-igenes
        vaurocs[[i]]<-aurocs
        save(eaurocs,vaurocs,file="results/043_naurocs_datlinger.rda")
    }
    emat<-eaurocs[[1]]
    vmat<-vaurocs[[1]]
    for(i in 2:100){
        emat<-rbind(emat,eaurocs[[i]])
        vmat<-rbind(vmat,vaurocs[[i]])
    }
    save(emat,vmat,file="results/043_naurocs_datlinger.rda")
    
}


### And plot!
load("results/043_naurocs_datlinger.rda")

# Means
eaurocs<-apply(emat,2,mean)
vaurocs<-apply(vmat,2,mean)
# Error bars
esds<-apply(emat,2,sd)
vsds<-apply(vmat,2,sd)
sds<-rbind(esds,vsds)

# Barplot
toplot<-rbind(eaurocs,vaurocs)
toplot<-toplot[,order(-toplot[2,])]

png("plots/043_datlinger.png",w=3000,h=2000,p=80)
par(las=2)
bp<-barplot(toplot,beside=TRUE,ylim=c(0.4,0.8),xpd=FALSE,col=c("salmon","navy"),main="Datlinger Dataset",ylab="AUROC")
legend("topright",col=c("salmon","navy"),pch=15,legend=c("Gene Expression","VIPER Activity"),lty=c(1,2),lwd=6)
abline(h=mean(eaurocs),col="salmon",lty=1,lwd=6)
abline(h=mean(vaurocs),col="navy",lty=2,lwd=6)
abline(h=0.5,col="darkgrey",lty=3,lwd=3)
par(las=1)
wt<-wilcox.test(as.vector(emat),as.vector(vmat),paired=TRUE)
mtext(paste0("p=",signif(wt$p.value,2)),cex=0.8)
# Err bar
segments(bp,toplot+sds,bp,toplot,lwd=3)
arrows(bp,toplot+sds,bp,toplot,lwd=3,angle=90,code=3,length=0.2)
dev.off()
```
