## Supplementary Code for "Pan-Cancer and Single-Cell modelling of genomic alterations through gene expression": geneids.R

```
require(AnnotationDbi)
require(org.Hs.eg.db)
require(biomaRt)
require(org.Mm.eg.db)


### annotategene function
annotategene<-function(x){
    tab<-AnnotationDbi::select(org.Hs.eg.db, keys=x, columns=c("GENENAME"), keytype="ENTREZID")
    tab<-tab[!duplicated(tab[,"ENTREZID"]),]
    out<-setNames(tab[,"GENENAME"],tab[,"ENTREZID"])
    out<-out[x]
    return(out)
}

### sym2eg function
sym2eg<-function(ids){
    list_symbol2eg <- as.character(org.Hs.egALIAS2EG[mappedkeys(org.Hs.egALIAS2EG)])
    ids <- as.character(ids)
    outlist <- list_symbol2eg[ids]
    names(outlist) <- ids
    outlist[is.na(outlist)] <- paste("unknown.", ids[is.na(outlist)], sep = "")
    outlist <- gsub("unknown.unknown.", "", outlist)
    return(outlist)
}


### any2entrez function
any2entrez<-function(x){
    tab<-AnnotationDbi::select(org.Hs.eg.db, keys=x, columns=c("ENTREZID"), keytype="ALIAS")
    symbols<-tab[,1]
    entrez<-tab[,2]
    dups<-which(duplicated(symbols))
    if(length(dups)>0){
        symbols<-symbols[-dups]
        entrez<-entrez[-dups]
    }
    out<-setNames(entrez,symbols)
    return(out)
}

### ens2eg function
ens2eg<-function(x){
    if(!exists("ens2egmap")){
        ens2egmap<<-as.list(org.Hs.egENSEMBL2EG)        
    }
    out<-ens2egmap[x]
    names(out)<-x
    out2<-sapply(out,function(x){
        if(is.null(x)){
            return(NA)
        } else {
            return(x[1])
        }
    })
    out3<-unlist(out2)
    out<-setNames(out3,names(out))
    return(out)
}

### eg2ens function
eg2ens<-function(x){
    if(!exists("eg2ensmap")){
        eg2ensmap<<-as.list(org.Hs.egENSEMBL[mappedkeys(org.Hs.egENSEMBL)])        
    }
    out<-eg2ensmap[x]
    names(out)<-x
    out2<-sapply(out,function(x){
        if(is.null(x)){
            return(NA)
        } else {
            return(x[1])
        }
    })
    out3<-unlist(out2)
    out<-setNames(out3,names(out))
    return(out)
}

### eg2sym function
eg2sym<-function(x){
    if(!exists("eg2symmap")){
        eg2symmap<<-as.list( org.Hs.egSYMBOL[mappedkeys( org.Hs.egSYMBOL)])        
    }
    x<-as.character(x)
    out<-eg2symmap[x]
    names(out)<-x
    out2<-sapply(out,function(x){
        if(is.null(x)){
            return(NA)
        } else {
            return(x[1])
        }
    })
    out3<-unlist(out2)
    out<-setNames(out3,names(out))
    return(out)
}


### eg2refseq function
eg2refseq<-function(x){
  if(!exists("ens2egmap")){
    eg2refseqmap<<-as.list(org.Hs.egREFSEQ)        
  }
  out<-eg2refseqmap[x]
  names(out)<-x
  out2<-sapply(out,function(x){
    if(is.null(x)){
      return(NA)
    } else {
      return(x[1])
    }
  })
  out3<-unlist(out2)
  out<-setNames(out3,names(out))
  return(out)
}


# Basic function to convert human to mouse gene names
convertHumanGeneList <- function(x){
    # BiocInstaller::biocLite('grimbough/biomaRt')
    require(biomaRt)
    human = useMart("ensembl", dataset = "hsapiens_gene_ensembl")
    mouse = useMart("ensembl", dataset = "mmusculus_gene_ensembl")
    genesV2 = getLDS(attributes = c("hgnc_symbol"), filters = "hgnc_symbol", values = x , mart = human, attributesL = c("mgi_symbol"), martL = mouse, uniqueRows=T)
    mousex <- setNames(genesV2[,2],genesV2[,1])
    return(mousex)
}


### eg2sym function for Mouse
eg2sym_Mm<-function(x){
    if(!exists("eg2symmap_Mm")){
        eg2symmap_Mm<<-as.list( org.Mm.egSYMBOL[mappedkeys( org.Mm.egSYMBOL)])        
    }
    out<-eg2symmap_Mm[x]
    names(out)<-x
    out2<-sapply(out,function(x){
        if(is.null(x)){
            return(NA)
        } else {
            return(x[1])
        }
    })
    out3<-unlist(out2)
    out<-setNames(out3,names(out))
    return(out)
}


# # Transcript level ENST to refseq
# library(org.Hs.eg.db)
# library("biomaRt")
# ensembl<-  useMart("ensembl", dataset="hsapiens_gene_ensembl")
# 
# values<- c("NM_001243076","NM_001949")
# getBM(attributes=c("refseq_mrna", "ensembl_gene_id", "ensembl_transcript_id", "hgnc_symbol"),
#       filters = "refseq_mrna", values = values, mart= ensembl)
# 
# 
# 
# values<-keys(org.Hs.egREFSEQ2EG)
# mapped<-getBM(attributes=c("refseq_mrna", "ensembl_gene_id", "ensembl_transcript_id", "hgnc_symbol"),
#               filters = "refseq_mrna", values = values, mart= ensembl)
# length(values)
# length(!is.na(mapped$ensembl_transcript_id))
```
