## Supplementary Code for "Pan-Cancer and Single-Cell modelling of genomic alterations through gene expression": scmodel.R

```
scmodel<-function(){

    # Gene track
    genetrack<-ifelse(anno==igene,"mut","wt")
    varmat<-varmat[,anno%in%c(igene,ctrlgenes)]
    genetrack<-genetrack[colnames(varmat)]
    
    # Build data object
    df<-cbind(t(varmat),genetrack)
    df<-as.data.frame(df)
    df$genetrack<-as.factor(df$genetrack)
    df$genetrack<-relevel(df$genetrack,ref="mut")
    
    # Set up caret
    outcomeName<-"genetrack"
    predictorsNames<-colnames(df)[colnames(df)!=outcomeName]
    
    # Split the data into training and test sets
    splitIndex<-createDataPartition(df[,outcomeName],p=0.75,list=FALSE,times=1)
    trainDF<-df[splitIndex,]
    testDF<-df[-splitIndex,]
    
    # Train Model Parameters (10-fold CV is acceptable)
    objControl<-trainControl(method="repeatedcv",number=10,returnResamp='none',summaryFunction=twoClassSummary,classProbs=TRUE)
    objModel<-train(data.matrix(trainDF[,predictorsNames]),trainDF[,outcomeName], 
                    method=method, 
                    trControl=objControl,  
                    metric="ROC",verbose=FALSE
    )
    
    # Validating the Model
    # It will predict using the best model: objModel$finalModel
    predictions<-predict(object=objModel,data.matrix(testDF[,predictorsNames]))
    probabilities<-predict(object=objModel,data.matrix(testDF[,predictorsNames]),type="prob")
    rocCurve<-roc(response=testDF$genetrack,predictor=probabilities[,"mut"])
    
    # Confusion matrix
    # confusionMatrix(predictions,testDF$gene.mut)
    
    # # Plot the ROC Curve with AUC
    # png(paste0("plots/024_modelROCs/ROC_",tum,"_MUT_",eg2sym(gene),".png"),w=1400,h=1000,p=35)
    # plot(rocCurve,col="red3",cex.lab=2,cex.main=1.1,print.auc=FALSE,lwd=2,
    #      main=paste0(method," Model predicting ",igene," Mutations"))
    # mtext("10-fold CV",side=1)
    # grid(col="lightgrey")
    # legend(0.2,0.2,legend=paste0("AUC= ", round(rocCurve$auc,4)))
    # dev.off()
    
    # Update auroc object
    aurocs<<-c(aurocs,rocCurve$auc)
}
```
